# WrapEvoFS enables auditable feature compression with regret-constrained representative locking

**DOI:** 10.64898/2026.09.28.755112

**Authors:** Junseo Park, Emma Schmidt, Shuxin Chi, Mostafa Fatehi Hassanabad, Leonard J. Foster, Huan Zhong, the Alzheimer’s Disease Neuroimaging Initiative

**Author notes:** Correspondence Huan Zhong, corresponding author. Correspondence Leonard J. Foster, co-corresponding author. Data used in preparation of this article were obtained from the Alzheimer’s Disease Neuroimaging Initiative (ADNI) database (adni.loni.usc.edu). The investigators within ADNI contributed to the design and implementation of ADNI and/or provided data but did not participate in the analysis or writing of this report. The complete investigator list is available in the ADNI acknowledgement list. **Author email addresses** Junseo Park, Emma Schmidt, Shuxin Chi, Mostafa Fatehi Hassanabad.

## Abstract

WrapEvoFS provides a deterministic, auditable rule for choosing one feature set from repeated stochastic searches. It retains candidates within a prespecified development-score gap, selects the most representative set by feature overlap, and records the decision before held-out evaluation. A corrected genetic objective preserves score ordering lost through zero truncation. In a fully nested four-class cancer analysis, all 30 locked signatures met the 0.01 tolerance, and 11 differed from the highest-scoring candidate; every changed selection increased mean feature overlap within the eligible pool. Paired differences in area under the receiver operating characteristic curve against recursive feature elimination with cross-validation all had 95% intervals crossing zero. No selector dominated matched comparisons on feature count and predictive performance, and one-time evaluation of frozen signatures at held-out centers yielded heterogeneous effects. WrapEvoFS standardizes selection within a fixed candidate bank under an empirical score-gap constraint, enabling reproducible reporting of performance, compression, and feature-agreement trade-offs without implying predictive superiority or stability under changes in participant composition.

## 1 Introduction

High-dimensional biomedical datasets often contain far more predictors than participants, making fitted models and selected signatures sensitive to preprocessing, finite samples, and stochastic search paths [1–3]. Feature selection can improve interpretability and reduce the number of measurements required for a model, but its evaluation is compromised if held-out outcomes inform preprocessing, hyperparameters, the stochastic seed, or the reported feature set [4–7]. Selecting the run with the most favorable held-out result introduces this bias even when each individual search uses development data only.

Recursive feature elimination with cross-validation (RFECV) provides an auditable elimination path and a target feature count, whereas genetic search explores nonnested subsets but introduces run-to-run variability [8–11]. Stability selection and ensemble feature selection address selection variability through resampling or aggregation [12, 13]. A distinct decision remains when repeated searches yield different feature sets with similar development scores and a single signature must be retained. Choosing the highest-scoring candidate ignores agreement with the alternatives, whereas choosing the most representative set without a score constraint can incur a substantial development-score loss.

WrapEvoFS addresses this issue by restricting selection to candidates within a prespecified score gap of the best observed candidate and choosing the set with the greatest average feature overlap among those candidates. The selected set is recorded and fixed before held-out evaluation, a step termed *locking*. For a fixed candidate bank and recorded scores, the selection rule is deterministic and its decision can be audited (Figure 1).

**Figure 1.**
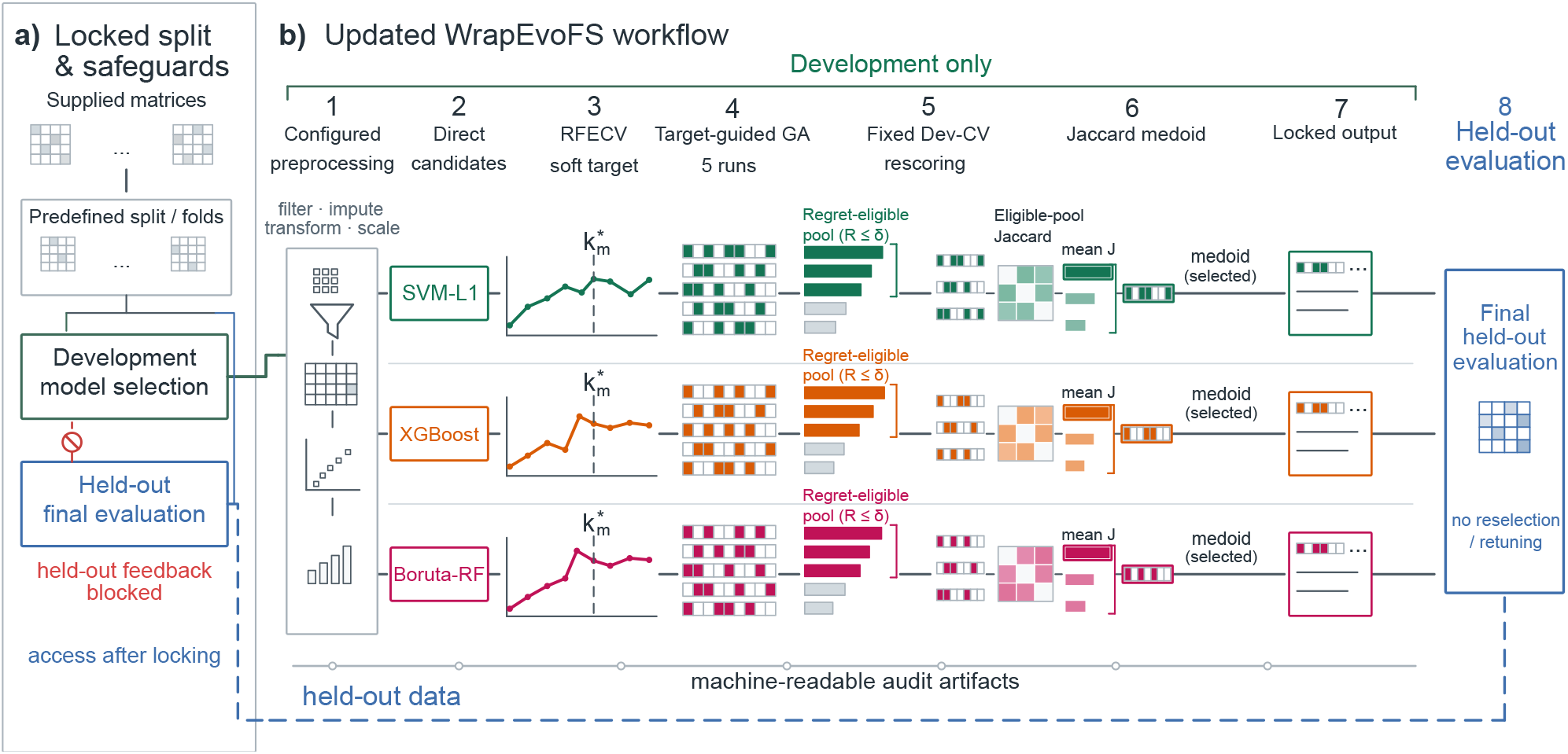
Development-only feature selection and representative locking. ia) Predefined partitions separate development-only selection from held-out evaluation. b) Preprocessing and initial feature screening (Direct) define three feature universes; RFECV supplies a target count, and five genetic searches generate candidate sets in each branch. Fixed development-CV rescoring identifies candidates within the allowed score gap (the eligible pool). The candidate with the greatest average Jaccard overlap within that pool (the medoid) is locked before held-out evaluation. The blue boundary marks held-out access after locking; the lower trace represents machine-readable audit records. The final estimator is fitted on development data after locking; held-out outcomes trigger no reselection or retuning.

This approach is related to model multiplicity and Rashomon-set analysis, in which near-optimal models can differ in their predictions or feature reliance [14, 15]. It differs from multiobjective evolutionary feature selection, which searches performance–cardinality trade-offs during candidate generation [16]. WrapEvoFS constrains the observed score loss after candidate generation and then selects a representative candidate. Nested selection bias and adaptive reuse of cross-validation scores remain as separate concerns [4, 6, 17], as do agreement across stochastic runs and stability under participant resampling [18].

An audit of the original WrapEvoFS analyses also identified a defect in the search objective: truncating a size-penalized score at zero made distinct negative scores indistinguishable. We correct this loss of ordering and formalize the subsequent selection step as regret-constrained representative locking. We evaluate the resulting workflow using fully nested resampling from supplied matrices, matched selector-layer and random-bank controls, and one-time evaluation of frozen signatures at held-out centers. These analyses assess the reproducibility of the selection decision and its performance, compression, and feature-agreement trade-offs; they do not assume a predictive advantage over alternative selectors.

We evaluate the corrected objective and locking rule under development-only, post-freeze held-out, and fully nested designs, while retaining archived analyses as explicitly labeled demonstrations.

## 2 Results

### 2.1 Objective flattening and corrected-objective verification

The corrected-objective configuration reduced aggregate deviation from the target feature count, with the largest reduction in the prespecified stress condition.

Eighteen of 36 original AMP-AD configurations triggered at least one zero-truncation warning; none of six original CGGA configurations did. The most severe condition was AMP-AD Rush/SVM-L1/Small: target *k*^∗^ = 3, locked count 84, absolute deviation 81, four of five zero-valued run-best fitness, and 245 all-zero generations. This provides a concrete loss-of-ordering mechanism beyond the general observation that genetic search is variable (Supplementary Fig. S1; Supplementary Tables S1–S2).

The matched development-only comparison comprised 120 completed GA runs in 24 combinations of center, branch, and Small or Reference cap (Figure 2; Supplementary Fig. S2; Supplementary Tables S1–S2). Aggregate absolute target deviation decreased from 216 to 137; the corrected-objective configuration was better in 12 conditions, unchanged in 6, and worse in 6. The prespecified Rush/SVM-L1/Small stress condition accounted for 56 of the 79 reduced deviation units; excluding it, deviation decreased from 135 to 112. All-zero generations decreased from 673 to 333 overall and from 428 to 267 without the stress condition.

**Figure 2.**
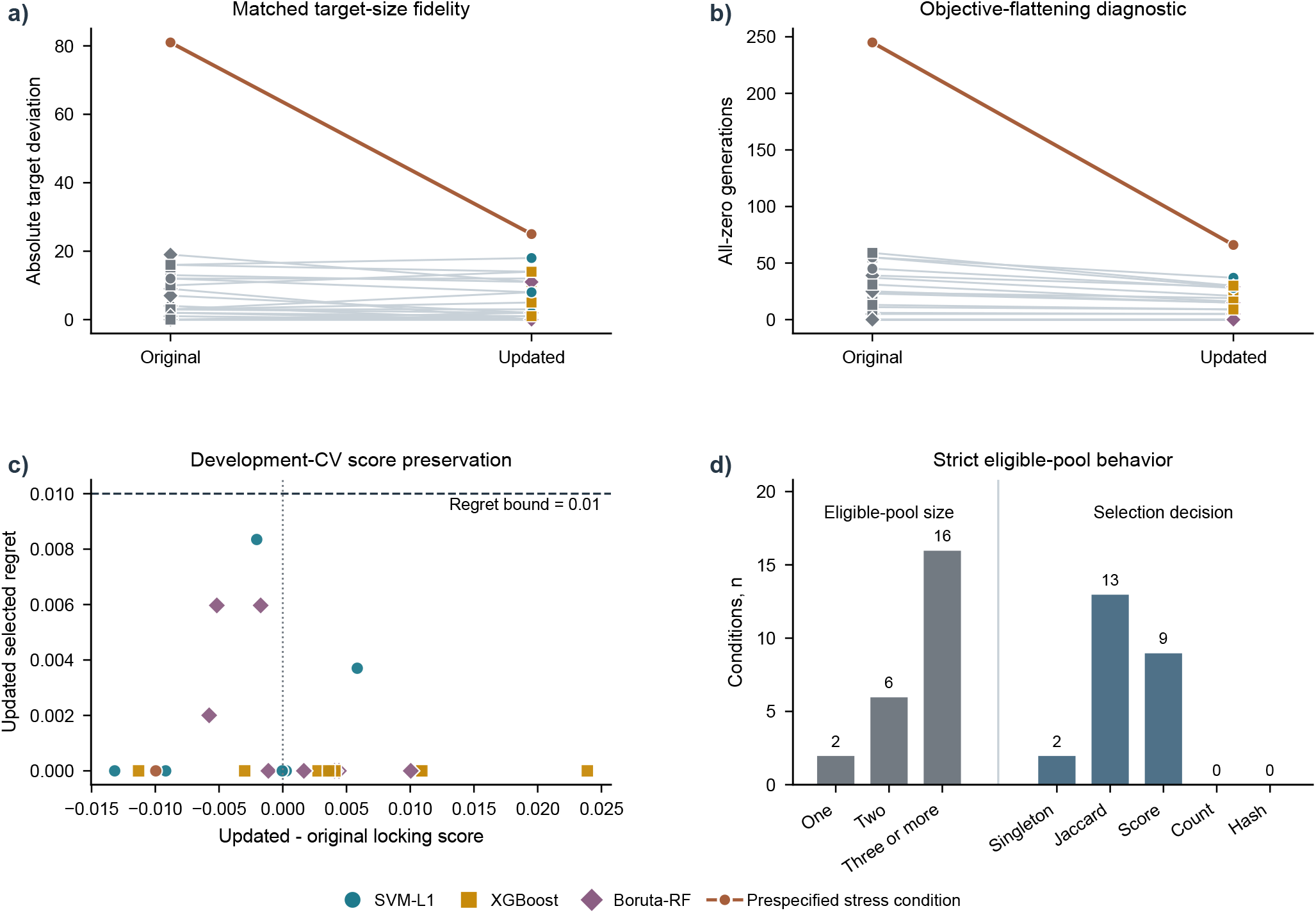
Corrected-objective search and score-constrained locking in the development-only comparison. The comparison includes 24 center–branch–cap conditions and 120 completed GA runs. a,b) Absolute deviation from the RFECV target (a) and all-zero-generation counts (b). Gray denotes original observations, colors denote corrected-objective branches, and the highlighted condition is Rush/SVM-L1/Small. c) Corrected-minus-original development-CV locking score versus the selected candidate’s score gap from the best candidate (empirical regret). Shapes identify branches; the dashed line marks the 0.01 tolerance. d) Eligible-pool sizes and final selection-decision stages; labels give condition counts. Candidate generation and locking both changed between configurations. No held-out outcomes informed this comparison.

All 24 locked sets satisfied the prespecified tolerance of 0.01 for empirical regret, the development-score gap from the best candidate in the same bank; the maximum selected regret was 0.00835 (Figure 2c). Corrected-objective-minus-original fixed locking-score differences had mean +0.00098, median +0.00096, and range −0.01319 to +0.02389; uniform parent-sampling fallbacks were zero. This comparison verifies configuration-level target fidelity and score-gap behavior but does not isolate either component. Held-out outcomes were accessed only after the protocol and all 24 selected masks were frozen.

### 2.2 Representative locking respects the development-score tolerance

Representative locking selected only candidates within the prespecified development-score tolerance. These candidates form the eligible pool; its medoid is the candidate with the greatest average Jaccard overlap with the other eligible sets. The four locking rules answer distinct decision questions (Table 2). The highest-locking-score candidate guarantees zero empirical regret but does not optimize representativeness. The unrestricted full-bank medoid optimizes full-bank representativeness without a nontrivial score-gap guarantee, whereas the legacy top-three medoid substitutes a rank cutoff for a metric-scale constraint. The regret-constrained medoid first enforces eligibility and only then optimizes within-pool mean Jaccard.

The existing-artifact 2 by 2 cross-lock analysis separated candidate-bank and locking-rule effects (Supplementary Table S3). Within 24 original-objective banks, the regret-constrained medoid changed 10 masks and reduced mean and maximum selected regret from 0.00261 and 0.02111 to 0.00060 and 0.00729, while total target deviation changed from 216 to 227. Within the 24 corrected-objective banks, it changed 8 masks and reduced mean and maximum regret from 0.00351 and 0.01525 to 0.00108 and 0.00835, while target deviation changed from 135 to 137. The rule therefore bounds configured score gaps without optimizing target fidelity.

At *δ* = 0.01, 2 of the 24 corrected-objective conditions had singleton eligible pools, 6 had two candidates, and 16 had at least three (Figure 2d; Supplementary Table S4). Unique Jaccard medoids determined 13 selections, higher locking score resolved 9, and singleton selection resolved 2; neither feature count nor the stable mask hash, a content-derived identifier used as the final tie-breaker, was needed. No exact duplicate mask occurred in the 48 complete banks, and all recovered Rush selections matched their archived and frozen-evaluation records.

Across 36 archived AMP-AD banks, retrospective regret-constrained re-locking compressed the corresponding Direct sets by a median 74.1% (range 15.8%–80.0%); median empirical score gap was 0 and the maximum was 0.0091. Across six CGGA banks, median compression was 57.9% (range 34.3%–70.0%), median gap was 0.00065, and maximum gap was 0.00355 (Supplementary Fig. S3; Supplementary Table S5). Recovery of the complete five-mask ADNI banks enabled the same audit without rerunning selection or accessing held-out outcomes. At absolute *δ* = 0.01, all three eligible pools were singletons and the selected empirical regret was zero. The rule retained the legacy XGBoost selection but changed the SVM-L1 and Boruta-RF selections, for which the legacy score gaps were 0.0140 and 0.0225, respectively (Supplementary Table S8).

### 2.3 Tolerance sensitivity and seed agreement

The tolerance controlled how many alternatives could be considered, while overlap among repeated-search outputs often remained modest.

Increasing absolute tolerance enlarged the strict eligible pool in AMP-AD and CGGA (Supplementary Fig. S4), and all 168 selections met their declared threshold without pool expansion. We fixed *δ* = 0.01 before locking. This absolute tolerance is defined on the reported metric scale and is not a universal threshold. Complete five-mask matrices yielded median selected mean Jaccard values of 0.204 and 0.424, respectively; median Nogueira coefficients were negative and 0.076. These values quantify stochastic agreement on fixed development samples, not participant-resampling or biomarker stability.

### 2.4 Fully nested benchmark reveals selector-dependent trade-offs

The fully nested benchmark produced compact, traceable signatures, but no selector consistently performed best across branches.

A separate TCGA GBM/LGG experiment moved the complete selection pipeline inside two repeated five-fold outer-CV partitions for 491 participants. The design yielded 150 candidates in 30 banks without outer-test access during selection or locking. Every selected candidate satisfied the prespecified 0.01 empirical-regret tolerance; the maximum was 0.00978. The regret-constrained medoid differed from the canonical highest-locking-score candidate in 11 banks, and every change increased eligible-pool mean Jaccard (median gain 0.0429; Figure 3a,b; Supplementary Fig. S5).

**Figure 3.**
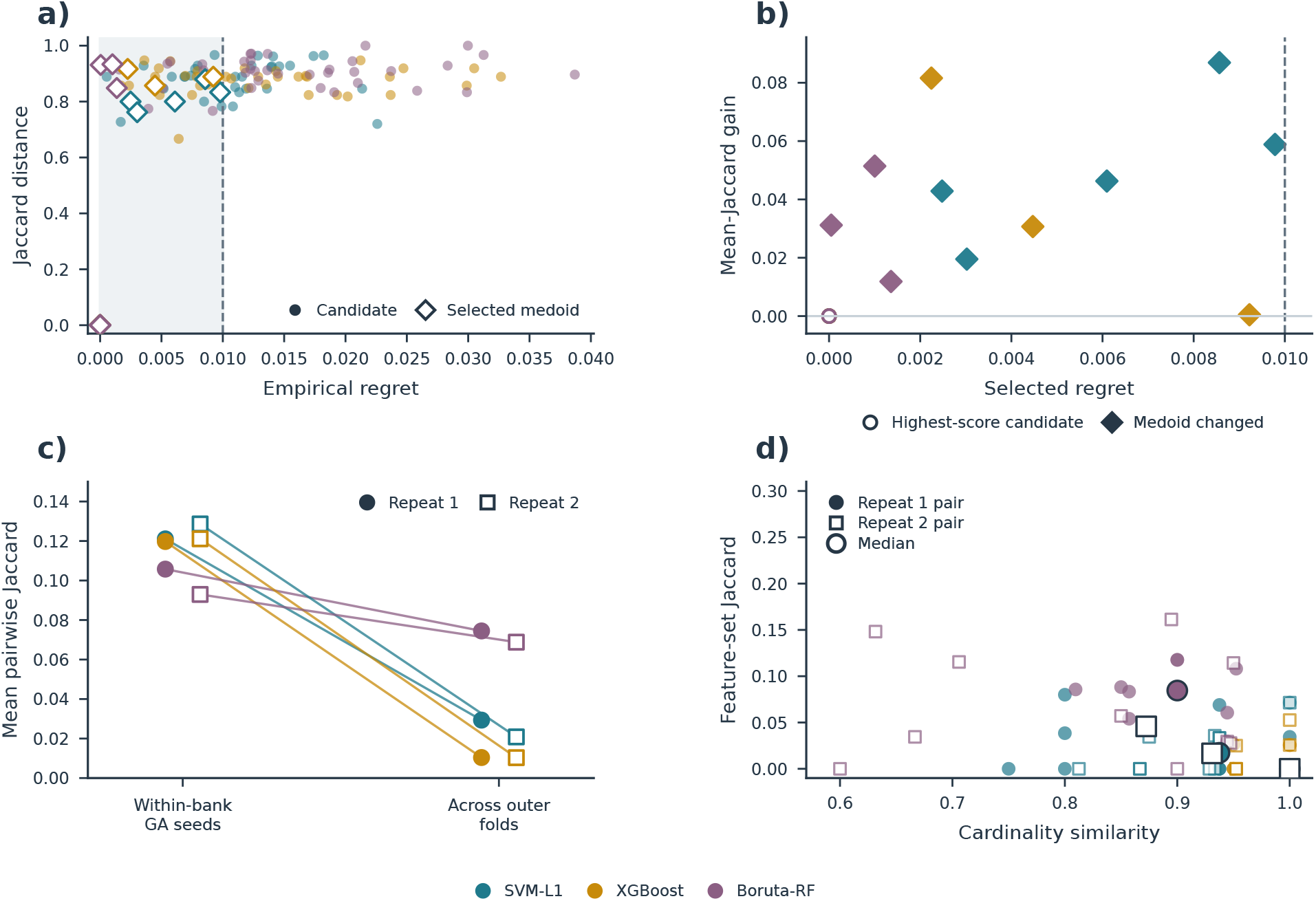
Representative locking and feature-set agreement in the fully nested TCGA analysis. Two five-fold outer-CV repeats across three Direct branches yield 30 banks of five candidates from 491 participants. a) Candidate development-score gaps and Jaccard distances from the highest-scoring candidate. Shading and the dashed line mark eligibility at *δ* = 0.01; open diamonds denote selected medoids. Highest-scoring candidates lie at the origin. b) Selected-minus-highest-scoring mean Jaccard overlap within each of 24 nonsingleton eligible pools. Filled diamonds denote 11 changed selections; open circles denote retained highest-scoring candidates. Singleton pools are excluded because this gain is undefined. c) Mean pairwise Jaccard overlap within candidate banks and among locked sets across outer folds, by repeat and branch. d) Feature-count similarity, min(*k*_1_, *k*_2_)*/* max(*k*_1_, *k*_2_), versus Jaccard overlap for 60 outer-fold pairs; large symbols denote coordinate-wise branch-repeat medians. Outer-test data were excluded from selection and locking. Overlap measures describe search and participant-partition sensitivity, not independent biomarker replication.

Mean within-bank pairwise Jaccard among the five GA candidates was 0.115, whereas mean Jaccard among locked signatures across outer folds was 0.0357 (Figure 3c). Across outer partitions, similar signature size did not imply similar feature identity: 42 of 60 paired locked signatures had cardinality similarity at least 0.9, yet 33 of those 42 pairs had feature-set Jaccard at most 0.05 (Figure 3d). Supplementary Fig. S6 reports chance-corrected candidate agreement, all dependent outer-fold pairwise similarities, and RNA/SCNV composition. These are descriptive measures of stochastic-search agreement and participant-partition sensitivity within this cohort, not estimates of biomarker stability.

We next assessed how these locked signatures compared with alternative selectors in predictive performance and feature count. The regret-constrained-medoid signatures contained a mean of 14.6, 20.0, and 18.0 features in the SVM-L1, XGBoost, and Boruta-RF branches, respectively, compared with 81, 100, and 100 Direct features. Mean repeated out-of-fold (OOF) macro one-vs-rest AUROC was 0.827, 0.827, and 0.828. Corresponding paired AUROC differences between WrapEvoFS and RFECV-only were −0.0108 (95% CI −0.0259 to 0.0035), 0.0012 (−0.0113 to 0.0134), and 0.0054 (−0.0125 to 0.0225); all intervals included zero (Figure 4a).

**Figure 4.**
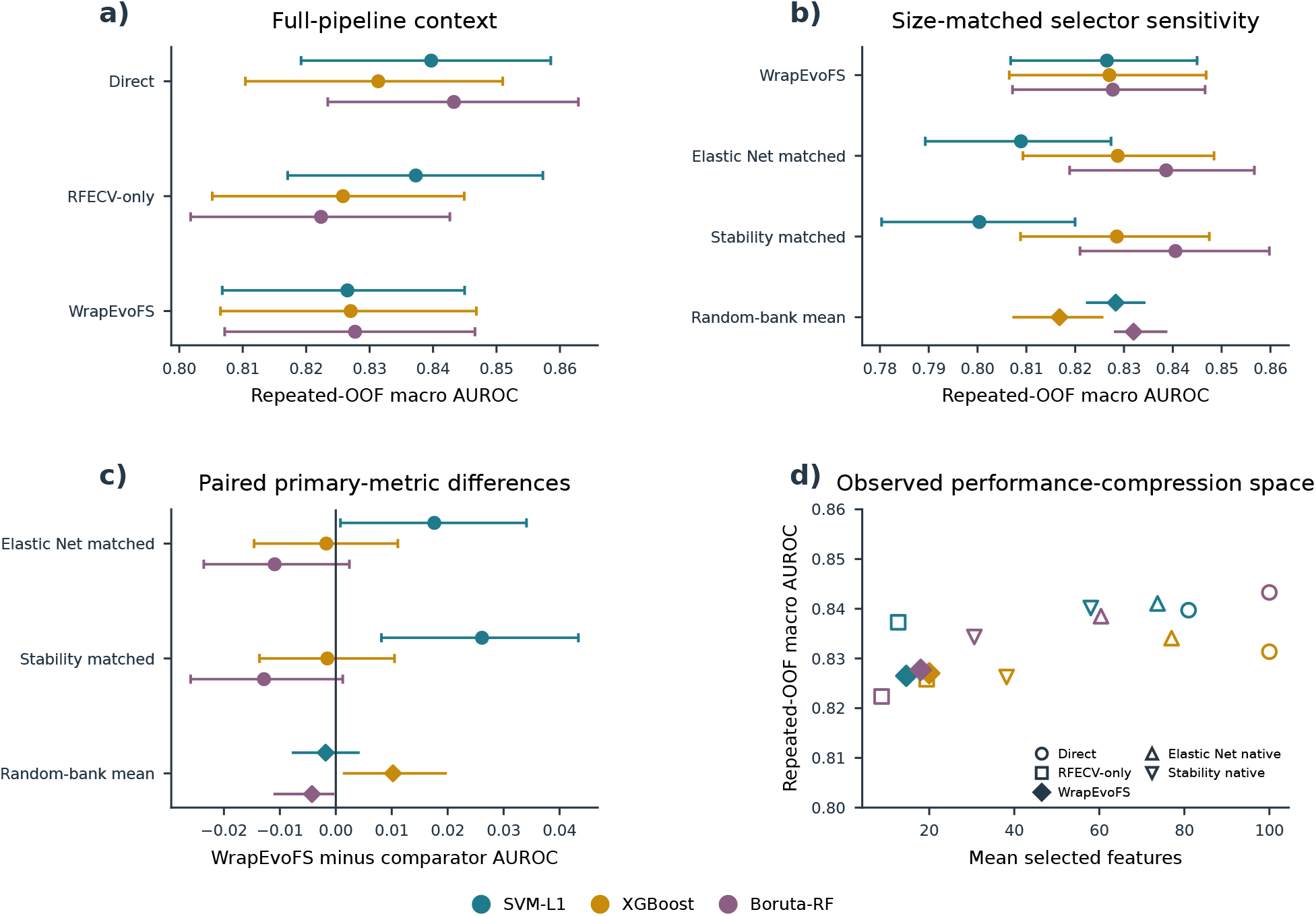
Predictive performance and feature counts in the fully nested TCGA benchmark. Analyses use the same 491 participants, two stratified five-fold outer-CV repeats, training-only selection, and a fixed final random forest. Colors identify Direct branches. a) Repeated out-of-fold macro one-vs-rest AUROC for Direct, RFECV-only, and WrapEvoFS. b) AUROC for WrapEvoFS, cardinality-matched Elastic Net and stability rankings, and five random-bank replicates (diamond, mean; range across replicates). c) Paired AUROC differences between WrapEvoFS and each comparator. Elastic Net and stability-selection 95% intervals use 2,000 participant-clustered, class-stratified bootstrap resamples; random-bank symbols show the range across five replicate banks. d) Mean feature count versus AUROC, including native Elastic Net and stability supports. Selector-layer comparisons share the post-Direct feature universe and do not establish standalone pipeline equivalence.

A retrospective selector-layer analysis conducted under a frozen protocol compared these signatures with nested multinomial Elastic Net, subsampling stability selection, and five independently seeded random banks within each identical post-Direct outer-training universe (Figure 4b–d; Supplementary Figs. S7–S8; Supplementary Table S6). At matched cardinality, SVM-L1 WrapEvoFS exceeded Elastic Net by 0.0176 (95% CI 0.0008 to 0.0341) and stability selection by 0.0261 (0.0082 to 0.0434). XGBoost differences were −0.0017 ( −0.0146 to 0.0111) and −0.0015 ( −0.0136 to 0.0105). Boruta-RF differences were −0.0109 ( −0.0236 to 0.0024) and −0.0128 ( −0.0260 to 0.0012). Native Elastic Net and stability supports used mean branch-specific counts of 60–77 and 31–58 features, whereas WrapEvoFS used 15–20. No selector dominated all branches on both performance and feature count. WrapEvoFS provides a compact, traceable hand-off subject to the regret constraint; predictive performance remained branch dependent.

### 2.5 Frozen selections undergo one-time multicenter evaluation

Predictive differences varied across held-out centers and configurations despite a common protocol that prohibited reselection or retuning.

After the 24 corrected-objective AMP-AD locks and the evaluation protocol were frozen, each signature was evaluated once in its corresponding held-out center without reselection or parameter changes (Figure 5; Supplementary Fig. S9; Supplementary Table S7). Pooled differences ranged from −0.0404 to +0.0480 against RFECV-only and from −0.0311 to +0.0176 against the legacy top-three medoid. Four of six RFECV-only contrasts and five of six legacy-top-three contrasts had 95% intervals including zero. Of the three non-zero-crossing intervals, the two RFECV-only contrasts favored opposite directions across branches, whereas the single legacy-top-three contrast favored the legacy rule. Because the masks and evaluation protocol were frozen before held-out outcome access, this analysis evaluates protocol adherence and center transport. The heterogeneous effects do not support a uniform predictive claim.

**Figure 5.**
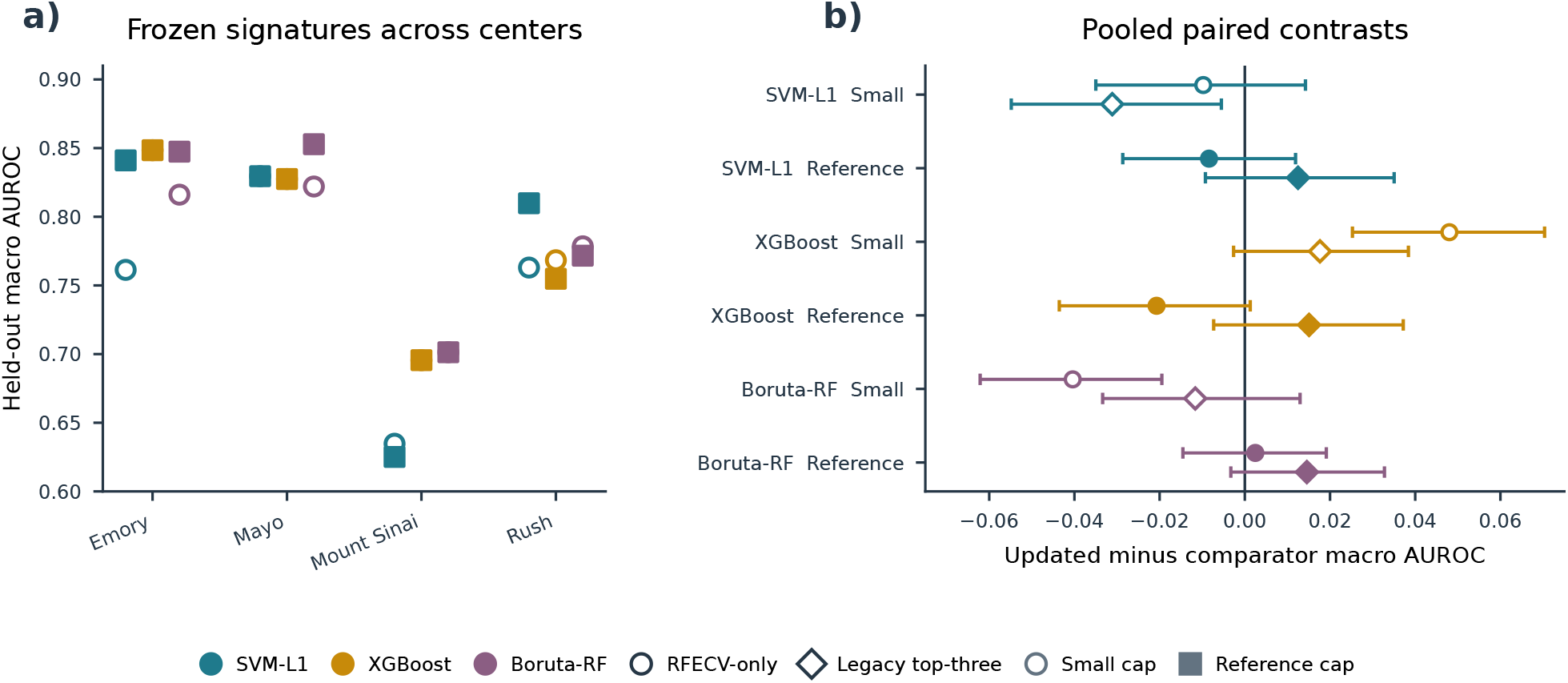
One-time held-out-center evaluation of frozen AMP-AD signatures. a) Macro one-vs-rest AUROC for 24 center–branch–cap conditions. Colors identify branches; open circles and filled squares denote Small and Reference caps, respectively. b) Pooled AUROC differences relative to RFECV-only (circles) and the legacy top-three medoid (diamonds), with 95% intervals from 2,000 common resamples stratified by held-out center and class (seed 42). Open and filled symbols denote Small and Reference caps. Masks, parameters, and the evaluation protocol were frozen before held-out outcomes were accessed; no result triggered reselection or retuning.

### 2.6 Archived and cross-modal analyses define empirical scope

The archived cohorts and radiomics analysis broadened the tested settings but did not establish a uniform predictive benefit.

Archived original-configuration demonstrations remain fully reported in the Supplement. ADNI locked signatures reduced Direct counts by 44%–70%, but all nine primary paired held-out intervals crossed zero (Supplementary Figs. S10–S12; Supplementary Tables S8–S9). Original AMP-AD effects varied by center, branch, cap, metric, and comparator (Supplementary Figs. S13–S14; Supplementary Tables S10–S12). CGGA showed 54%–73% compression and variable five-run agreement without a uniform predictive advantage in its coherent same-split benchmark (Supplementary Figs. S15–S19; Supplementary Tables S13–S18).

The VGH brain-tumour radiomics cohort (UBC REB H20-02354, “Brain Tumor Image Analysis”) provided a supplementary internal test in a distinct feature modality. The same 197 MGMT-linked participants were analyzed under a frozen 137/60 development/held-out split in full and label-independent perturbation-filtered feature spaces (Supplementary Fig. S20). All six locks had zero empirical development-CV regret at *δ* = 0.01, with 10.0%–81.0% compression relative to Direct sets (median 75.6%). Paired held-out locked-minus-Direct AUROC estimates ranged from −0.007 to 0.069, and all 95% intervals included zero (Supplementary Table S19). This analysis broadens the tested modality but reuses participants across feature spaces, relies on unadjudicated automatic regions, and remains internal.

### 2.7 Secondary held-out trade-offs and interoperability

The secondary analyses showed that a locked feature set can be reused reproducibly downstream, although results remained method dependent.

Secondary tuned-model, computational, random-subset, and Elastic Net analyses are reported in Supplementary Figs. S21–S23 and Supplementary Table S16. Bayesian, STABL, and BLiP analyses used the locked ADNI outputs as reproducible downstream candidate sets without reopening held-out-informed selection (Supplementary Figs. S24–S27; Supplementary Tables S20–S22). Five of eight features selected by training-only STABL overlapped the 78-feature WrapEvoFS locked-signature union, yielding an 81-feature candidate set for restricted downstream analyses (Supplementary Figs. S26–S27; Supplementary Table S21). The corresponding 81-feature restricted Laplace Bayesian model yielded a held-out macro one-vs-rest AUROC of 0.803 (95% CI 0.744–0.856), compared descriptively with 0.780 for the 7,411-feature Bayesian reference, for which a confidence interval was unavailable (Supplementary Figs. S24–S25 and S27; Supplementary Tables S20–S22). Directions varied by method, and agreement is not independent replication because the analyses reuse the same development boundary.

### 2.8 Controlled candidate-bank locking simulation

The simulation reproduced the empirical score-gap guarantee, but no locking rule consistently minimized loss in the hidden utility used to generate candidate scores.

A prespecified simulation isolated the locking layer under hidden utility across 4,995,000 synthetic banks (Supplementary Fig. S28; Supplementary Table S23). In the primary *R* = 5, *δ* = 0.01 design, the regret-constrained medoid had no empirical-regret violation and increased mean full-bank Jaccard from 0.424 to 0.446 relative to highest-locking-score selection, but mean hidden-oracle regret was 0.00259 versus 0.00222. It combined lower oracle regret with higher representativeness in 91 of 162 primary scenarios, with relative behavior depending on score noise, consensus alignment, bank size, and tolerance. The simulation quantifies the feasibility–representativeness trade-off; it does not show uniformly lower oracle regret.

## 3 Discussion

WrapEvoFS separates stochastic candidate generation from the decision to retain one feature set. Its corrected objective preserves penalized-objective ordering that zero truncation can erase, and regret-constrained representative locking imposes an explicit empirical development-CV score-gap constraint before selecting the eligible-pool medoid. The fully nested benchmark adds a practical result: no tested selector dominated every Direct branch on both primary predictive performance and feature count. WrapEvoFS retained 15–20 features; compared with equally sized Elastic Net and stability rankings, it had higher estimates in the SVM-L1 branch, similar estimates in the XGBoost branch, and lower point estimates with zero-crossing intervals in the Boruta-RF branch. The post-freeze AMP-AD analysis further demonstrated that masks and parameters can be fixed before held-out-center outcomes are accessed.

The comparator results clarify when the additional stochastic search and locking layer may be useful. Native Elastic Net and stability supports were substantially larger, while their AUROC relative to matched-cardinality supports varied by branch and selector. At equal cardinality, the direction depended on the upstream Direct universe, and random-bank means likewise varied by branch. Pipeline comparison and locking-rule comparison therefore answer different questions: RFECV, Elastic Net, and stability selection generate feature sets, whereas the highest-locking-score candidate, legacy top-three medoid, unrestricted full-bank medoid, and regret-constrained medoid choose among a supplied stochastic bank. The proposed rule is most relevant when nonnested candidate exploration is scientifically justified and one compact, traceable output must be retained.

Five saved seeds reveal dispersion across run-best masks, while the nested TCGA experiment shows a distinct participant-partition effect: similar selected counts can coexist with markedly different feature identities across outer partitions (Figure 3c,d; Supplementary Fig. S6). The nested analysis therefore separates signature-size consistency from feature-identity sensitivity. Representative locking standardizes the candidate passed downstream, but it does not resolve sensitivity to participant composition. Seed agreement and participant-resampling stability should therefore be reported as different estimands [12, 13, 18].

The formal guarantee is finite-bank and empirical: it concerns only the configured development-CV score gap. In archived and fixed-partition analyses, preprocessing and Direct screening were fitted on each full development partition rather than within locking-score folds, and repeated fold reuse makes locking scores adaptive internal selection scores [17]. Five seeds are sparse. In TCGA, all study-fitted steps from the supplied matrix were nested, while the new selector comparison is conditional on post-Direct universes, remains internal, and was performed retrospectively under a frozen protocol after the parent TCGA results existed. The radiomics analysis used one clinical cohort with automatic, non-adjudicated regions and no repeat segmentation, and post-freeze center effects were heterogeneous. Broader cohorts, participant-resampling designs, and biological follow-up are needed to establish transportability and biomarker reproducibility; these boundaries do not alter the proved score-gap guarantee or reproducible selection audit.

WrapEvoFS provides an auditable rule that standardizes selection of one representative subset from a stochastic candidate bank under an explicit empirical development-CV score-gap tolerance while reporting signature size and feature-identity sensitivity separately.

## 4 Methods

### 4.1 Overall analytical workflow and leakage boundary

Figure 1 separates development-only selection from one-time held-out evaluation. Preprocessing and Direct screening precede RFECV target estimation; five genetic searches per branch generate retained run-best subsets that a common development-CV evaluator rescores. A prespecified tolerance defines the regret-eligible pool, whose Jaccard medoid is locked under deterministic tie-breaking. Held-out outcomes cannot determine the strategy, tolerance, objective, metric, tie rule, or selected run, and trigger no reselection. Complete pseudocode and executable settings are provided in the Supplementary Methods and Supplementary Table S24.

### 4.2 RFECV target and primary estimands

For each Direct branch (SVM-L1, XGBoost, or Boruta-RF) [9, 19, 20], RFECV follows a recursive elimination path on development folds and supplies a soft target: the smallest feature count attaining the best mean score among counts allowed by the configured cap. Compression is the fraction of Direct features removed by a selected set.

Let *F*_*r*_ be the feature set of retained candidate *r* in a bank of *R* candidates. All candidates are evaluated on the same *K*_*L*_ development folds using a fixed-specification random forest [21] and a metric oriented so that larger values are better. If 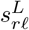 is candidate *r*’s score on validation fold *£*, its locking score *L*_*r*_, empirical regret *R*_*r*_, and eligible pool *E*_*δ*_ are

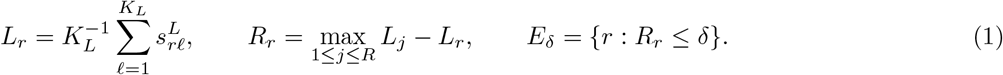

Here *δ* ≥0 is the prespecified absolute score-gap tolerance, and *E*_*δ*_ contains candidate indices. Each fold score is obtained by fitting on the remaining folds and scoring the validation fold. All candidates share the fold partition, metric, and estimator specification; run-specific estimator seeds remain in the audit record. The search-stage metric may differ from the locking metric: search generates candidates, whereas the common locking evaluator compares them. Run-to-run agreement is summarized by pairwise Jaccard overlap, the selected candidate’s mean overlap, and the chance-corrected Nogueira coefficient only when a complete run-by-feature matrix exists [18]. These quantities describe stochastic agreement on one fixed development sample, not biomarker stability under participant resampling. Held-out performance differences are secondary and are calculated only after locking. Full notation for RFECV targeting, compression, locking scores, and held-out contrasts is provided in Supplementary Methods, Section S1.2.

### 4.3 Legacy-compatible and corrected genetic objectives

A nonempty binary chromosome **z** represents feature set *F*_*m*_(**z**) in Direct branch *m*. Let *S*_*G,m*_(**z**; *ξ*) be its mean search-stage score across fixed development folds with recorded seed *ξ*, 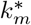 the RFECV target, and *λ* the penalty per feature of absolute target deviation. The penalized objective and its legacy-compatible transformation are

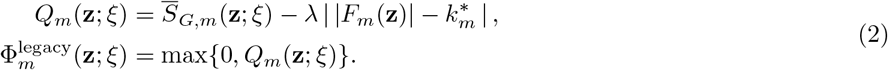

Zero truncation makes distinct negative objectives indistinguishable. The corrected mode therefore uses *Q*_*m*_ directly for reporting, ranking, elite selection, and run-best updating. A nonnegative shift is applied only to proportional parent-sampling weights and preserves the ordering of finite objectives. Uniform parent sampling is recorded when finite scores cannot define valid weights; empty chromosomes are prohibited. These scoring folds evaluate a frozen development-derived feature matrix: preprocessing and Direct screening are not refitted within each search-scoring fold. The package records zero-truncation, target-deviation, sampling-fallback, objective-diversity, and mask-diversity diagnostics. Fold-level definitions, sampling probabilities, and edge-case rules are provided in Supplementary Methods, Section S1.2.

### 4.4 Regret-constrained medoid locking

The eligible pool in Equation (1) is never expanded with a candidate outside the declared threshold. Because a highest-scoring candidate has zero regret, the pool is nonempty. If it contains one candidate, that candidate is selected. Otherwise, each eligible candidate’s mean Jaccard similarity, defined as its average feature overlap with the other eligible candidates, is

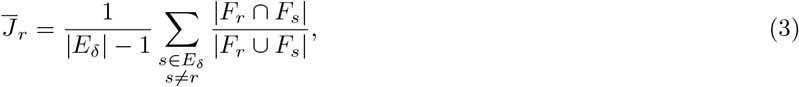

The candidate with the largest mean similarity is the eligible-pool Jaccard medoid. Remaining ties are resolved, in order, by higher locking score, fewer features, and a lexicographically smaller stable mask hash. The hash resolves otherwise identical selection criteria deterministically and is not a scientific score. Exact duplicate candidates retain their multiplicity as voting candidates, and source-run provenance is used only after the scientific feature set has been selected.

Selection is restricted to the eligible pool, so the output always satisfies the declared empirical score-gap tolerance. At zero tolerance, only highest-scoring candidates are eligible; tied maxima still undergo medoid selection. If the tolerance includes every candidate, the rule becomes the full-bank medoid. These guarantees concern the configured empirical development-CV score gap, not expected predictive risk, generalization error, or statistical uncertainty. Complete propositions, proofs, duplicate-policy details, and implementation correspondence are given in the Supplementary Methods and Supplementary Table S25.

With *R* retained candidates, *E* eligible candidates, canonical-universe size *p*, and average selected-set size *s*, the package’s dense-mask, sparse-Jaccard implementation has expected time *O*(*Rp*+*E*^2^*s*+*R* log *R*) and memory *O*(*Rp*+*E*^2^). For the present *R* = 5 design, at most 10 unordered eligible pairs are compared; this use case does not establish large-bank scalability. Detailed representation-specific derivations are in the Supplementary Methods.

The corrected-objective configuration fixes absolute *δ* = 0.01 before locking. It is a metric-scale tolerance, not a universal threshold. Development-only sensitivity evaluates 0, 0.005, 0.01, 0.02, and a best-run-SE-scaled threshold where fold vectors exist. The scaled sensitivity is not the conventional paired one-standard-error rule and stops when required fold vectors are absent.

### 4.5 Controlled candidate-bank locking simulation

The frozen simulation protocol evaluated highest-locking-score selection, the legacy top-three medoid, the unrestricted full-bank Jaccard medoid, and the regret-constrained medoid under hidden synthetic utility. The primary design crossed three candidate topologies, three heterogeneity levels, three proxy strengths, and six score-noise ratios at *R* = 5 and *δ* = 0.01, with 10,000 independently indexed banks per scenario. Prespecified sensitivities varied *R, δ*, score-noise distribution, and duplicate rate, yielding 513 scenarios and 4,995,000 banks in total. Primary summaries weight the 162 scenarios equally; the scenario grid is not treated as a population prior. Oracle-regret failure was defined as hidden-utility regret greater than 0.02. The simulation used no participant-level data, classifier fitting, RFECV, Direct selection, or GA. Complete utility, candidate-generation, tie-path, execution-integrity, and claim-boundary details are provided in the Supplementary Methods, Supplementary Fig. S28, and Supplementary Table S23.

### 4.6 Development-only objective comparison

The comparison covered four AMP-AD held-out-center development partitions, three Direct branches, two feature caps, and five seeds per condition (24 conditions; 120 full GA runs). The matched original configuration used the archived zero-truncated objective and legacy top-three medoid. The corrected configuration used *λ* = 0.015, untruncated ranking and elitism, shifted parent-sampling weights, fixed five-fold development-CV macro one-vs-rest AUROC rescoring, and regret-constrained locking at *δ* = 0.01. Because candidate generation and locking both changed, this was a configuration-level comparison. Held-out-center outcomes were not accessed.

### 4.7 One-time post-freeze AMP-AD held-out evaluation

All 24 masks, locking scores, hyperparameters, metric orientation, preprocessing boundary, final estimator, bootstrap plan, and source provenance were frozen in a prespecified protocol before guarded one-time evaluation at the corresponding held-out center. Each signature was fitted on its complete development partition using the frozen 150-tree random-forest specification; no result triggered reselection or configuration change. Macro one-vs-rest AUROC was primary, with macro AUPRC, balanced accuracy, macro F1, and accuracy secondary. Comparisons with RFECV-only and legacy-top-three-medoid predictions used 2,000 common resamples stratified by center and class (seed 42). The four centers represent four distribution shifts; the 24 conditions are not independent cohorts.

### 4.8 CGGA coherent benchmark and saved-bank nested sensitivity

The coherent benchmark reconstructed the archived 214/92 CGGA split and verified participant IDs, labels, and compact-matrix values against the raw source. RFECV-only, size-matched Elastic Net [22], highest-score, legacy-top-three-medoid, unrestricted-full-bank-medoid, and regret-constrained-medoid signatures were evaluated with the same fixed 500-tree random forest. AUROC intervals and paired contrasts used 2,000 common class-stratified resamples (seed 42). No feature-selection stage was rerun.

For the nested sensitivity, the five saved candidate banks from a reduced-budget CGGA analysis were re-locked at *δ* = 0.01 while preserving outer-fold preprocessing, feature universes, masks, and candidate scores. A 500-tree random forest was refitted in each archived outer-training fold. This post hoc internal sensitivity used 214 participants, five outer folds, and a historical 20-individual, 12-generation GA budget; it did not access the 92-sample held-out partition or replace the primary analysis.

### 4.9 Fully nested TCGA GBM/LGG analysis

The TCGA analysis used a supplied workbook containing 491 participants, 20,118 processed RNA-seq features, and 5,000 cleaned SCNV features. The provider confirmed label-independent variance filtering; a label-blind audit recovered 4,999 features by ranking the 24,776-feature uncleaned sheet, with an exact variance tie at the cutoff. This upstream reduction was the declared empirical starting boundary. Untreated primary GBM, astrocytoma, oligodendroglioma, and oligoastrocytoma counts were 66, 147, 165, and 113 [23–25]. Modality prefixes were retained; clinical columns, RPPA, and uncleaned SCNV were excluded.

Two shuffled stratified five-fold outer-CV repeats used random states 20260810 and 20260811. Within every outer-training partition, median imputation, zero-variance filtering, Direct screening, RFECV target estimation, five GA runs, fixed candidate rescoring, and locking were refitted without outer-test access. Post-Direct universes were capped deterministically at 100 features. RFECV, GA fitness, and locking used macro one-vs-rest AUROC. Each GA used 50 individuals, 50 generations, seeds 42–46, the corrected untruncated shifted-linear objective, and *λ* = 0.015. The five saved run-best candidates were rescored using a fixed five-fold, 500-tree random-forest evaluator and locked with absolute *δ* = 0.01, strict eligible-only pooling, and canonical deterministic tie-breaking. This produced 30 outer-training candidate banks and 150 GA runs.

After locking, Direct, RFECV-only, highest-locking-score, legacy top-three-medoid, unrestricted-full-bank-medoid, and regret-constrained-medoid signatures were evaluated on the corresponding outer-test fold using the same 500-tree random forest and a common deterministic seed within each repeat–fold–branch condition. Mean repeat-specific OOF macro one-vs-rest AUROC was the primary predictive summary. Confidence intervals and paired differences between the regret-constrained medoid and each comparator used 2,000 participant-clustered, class-stratified bootstrap resamples that retained both repeat-specific predictions for each sampled participant. The pairwise outer-fold Jaccard values share training observations and were therefore summarized descriptively rather than treated as independent replicates. Complete execution hashes, agreement definitions, duplicate handling, and source-table correspondence are reported in the Supplementary Methods.

The selector-layer sensitivity conducted under the frozen protocol reused each verified post-Direct outer-training universe without rerunning prior selection stages. Nested multinomial Elastic Net produced native and matched-cardinality supports; subsampling stability selection used 50 class-stratified half-samples and a 0.80 frequency threshold. Five random five-mask banks per condition matched feature count and locking-evaluation budget. Feature hashes preceded outer-test access, and signatures shared the condition-specific final-estimator seed. Because TCGA outcomes had already been analyzed, this is a retrospective nested sensitivity. Complete grids, seeds, integrity records, and reporting boundaries are in the Supplementary Methods and Supplementary Table S6.

### 4.10 Software implementation and audit artifacts

WrapEvoFS v0.2.0 supports Python 3.10–3.12 through an API and command-line interface. Configurations select legacy-compatible or corrected-objective search and locking modes; archived results are never silently replaced. Machine-readable outputs record scores, regrets, eligibility, overlap, selected masks, seeds, tie paths, fingerprints, and GA diagnostics. Pickle-free atomic checkpoints reject incompatible state. Complete configuration, schema, hashing, checkpoint, and serialization details are provided in the Supplementary Methods and repository documentation.

### 4.11 Empirical datasets and analysis boundary

The five empirical designs are summarized in Table 1, with preprocessing boundaries and archived original settings detailed in Supplementary Tables S26–S27. The ADNI-derived multiclass analysis contains 532 development and 133 held-out participants with 7,411 processed predictors [26, 27]. AMP-AD contains 527 participants and 13,136 supplied frontal-proteomic and metabolomic predictors from the AMP-AD Diverse Cohorts Study in four fixed leave-one-center-out evaluations [28]. The CGGA MGMT-methylation analysis contains 214 development and 92 held-out samples with 24,326 aligned gene-expression predictors [29]. The fully nested TCGA GBM/LGG analysis contains 491 participants with 20,118 supplied RNA-seq and 5,000 cleaned SCNV predictors [23, 24]. The VGH brain-tumour radiomics analysis contains 137 development and 60 held-out participants with 1,781 full-space features and a 1,346-feature perturbation-filtered sensitivity space. These are internal demonstrations beginning from supplied matrices; independent external validation was not performed.

**Table 1.**
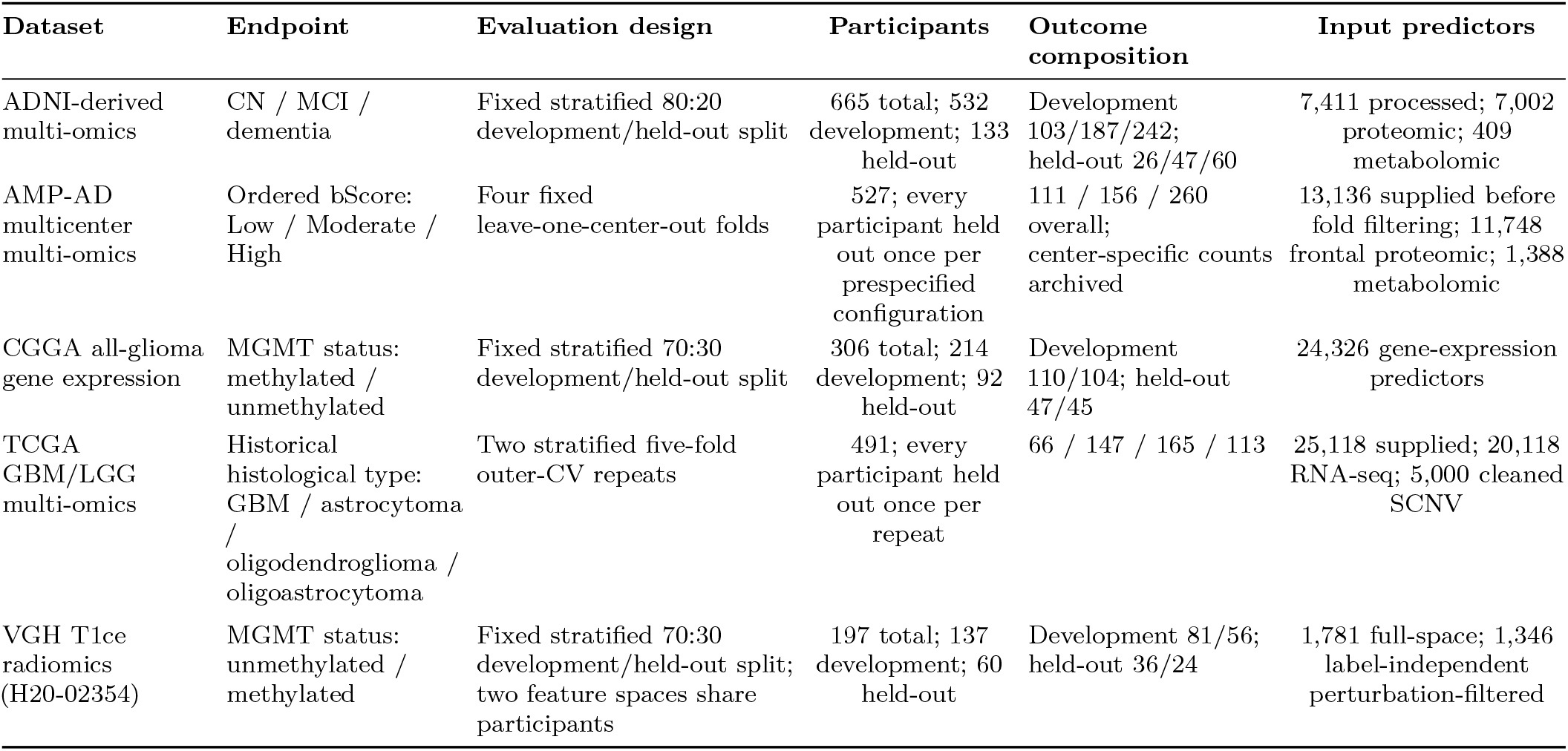
Empirical datasets, evaluation designs, outcome composition, and predictor inventories.

**Table 2.** Decision rules for selecting one feature set from a finite stochastic candidate bank.

| Rule | Admissible pool | Primary decision | Empirical-regret guarantee | Representativeness target |
| --- | --- | --- | --- | --- |
| Highest-locking-score candidate | Full bank | Maximize locking score | Zero | Not explicitly optimized |
| Legacy top-three medoid | Three highest-ranked candidates | Maximize mean Jaccard within the rank-restricted pool | No metric-scale bound | Rank-restricted pool |
| Unrestricted full-bank medoid | Full bank | Maximize mean Jaccard within the full bank | No nontrivial bound | Full candidate bank |
| Regret-constrained medoid | $E_\delta = \{r : R_r = L_{\text{best}} - L_r \leq \delta\}$ | Maximize mean Jaccard within $E_\delta$ | $R_{\text{selected}} \leq \delta$ | Strict regret-eligible pool |

The ADNI-derived source data were obtained from the Alzheimer’s Disease Neuroimaging Initiative database (https://adni.loni.usc.edu/). ADNI was launched in 2003 as a public–private partnership led by Principal Investigator Michael W. Weiner, MD, to evaluate whether serial imaging, biological markers, and clinical and neuropsychological assessment can jointly measure progression of mild cognitive impairment and early Alzheimer’s disease.

For the VGH brain-tumour radiomics analysis, prespecified header and geometry rules selected one eligible T1ce series per participant without MGMT access. DICOM series were converted to pseudonymized NIfTI with dcm2niix [30]; RaidionicsRADS and RaidionicsSeg generated automatic brain and tumor-core candidate regions [31], which were not manually corrected or treated as ground truth. MGMT linkage followed series selection, conversion, segmentation, configuration freezing, PyRadiomics extraction [32], and label-independent filtering. Of 326 source participants, 262 were T1ce eligible, 259 completed extraction, and 197 had unique validated MGMT linkage (117 unmethylated and 80 methylated); no labels were inferred for unlinked records. A fixed stratified 70:30 split yielded 137 development and 60 held-out participants. A separate deterministic 15-participant pilot froze the perturbation-filtered space using original, 1-mm-eroded, and 1-mm-dilated masks; retention required ICC(2,1) ≥0.85 and median relative range ≤0.25 without outcome labels [33]. Supplementary Table S28 reports the frozen extraction settings, complete source flow, and validation boundary; IBSI provides the feature-definition reference framework [34].

Within each feature space, SVM-L1, XGBoost, and Boruta-RF Direct screening and RFECV target estimation used development data only. Five GA searches used seeds 42–46, 50 individuals, 50 generations, five-fold ROC-AUROC scoring, the untruncated shifted-linear objective, and *λ* = 0.015. The five run-best sets were rescored by fixed five-fold development-CV ROC AUROC and locked with absolute *δ* = 0.01, strict eligible-only pooling, and canonical deterministic tie-breaking. All six locks were completed before a 500-tree random forest was fitted on the full development partition and evaluated once on held-out data. AUROC and AUPRC intervals and paired AUROC differences used 2,000 class-stratified bootstrap resamples with seed 42. The two feature spaces share participants, split, and outcomes; their contrast is a feature-definition sensitivity analysis, not external validation.

Retrospective sensitivities used only available masks, scores, and fold vectors; missing candidate artifacts were not reconstructed. Complete cohort-specific availability is reported in the Supplement.

Except in the fully nested TCGA experiment, preprocessing and label-dependent Direct screening were fitted once on each full development partition and were not refitted inside every locking-score fold. RFECV, GA evaluation, and locking reused configured development folds during adaptive selection; the locking evaluator used fixed stratified five-fold construction with random state 42. Candidate evaluators used run-derived estimator seeds, so seed agreement may reflect search and estimator randomness. Development-CV scores are adaptive internal selection scores and do not provide unbiased generalization estimates. In TCGA, all study-fitted selection steps beginning from the supplied matrix were refitted inside each outer-training partition, and only the resulting outer-test predictions contributed to repeated OOF performance summaries. The provider’s label-independent 5,000-feature SCNV variance filter preceded outer CV and defines the supplied-data boundary.

### 4.12 Stage-metric audit and secondary analyses

Archived configurations used mixed stage metrics: ADNI and AMP-AD combined weighted multiclass AUROC, balanced accuracy, and the locking metric; CGGA combined RFECV AUROC, GA accuracy, and locking AUROC. These original workflows are staged heuristics, not unified optimizers. The new binary radiomics analysis aligned RFECV, GA, and locking on ROC AUROC under the corrected-objective configuration. The package’s scoring.unified_metric: macro_ovr_auroc option maps to binary ROC AUROC or multiclass macro one-vs-rest AUROC when compatible; it does not retroactively alter archived runs.

Held-out results were joined only after each development-only rule was fixed. A held-out estimate for a newly selected rule was reused only when its selected run exactly matched an archived evaluated run; otherwise it remains unavailable. The CGGA paired AUROC contrasts between the archived legacy top-three medoid and RFECV-only used 2,000 common class-stratified bootstrap resamples (seed 42) from the frozen aligned held-out predictions. These post hoc intervals did not affect candidate generation, locking, or the choice of *δ*; no held-out value compared locking strategies during selection.

Bayesian uncertainty quantification, training-only STABL, and BLiP are secondary interoperability analyses [35–39]. They show that development-derived locked outputs can define a downstream candidate set for posterior inference, resampling-oriented selection, and expected-FDR decisions. Detailed priors, diagnostics, stability paths, posterior summaries, and BLiP sensitivities are provided in the Supplementary Information. Cross-method overlap is not interpreted as biomarker validation.

### 4.13 Ethics and consent

This study performed secondary methodological analyses of existing data and involved no new participant recruitment, intervention, or recontact. ADNI protocols were approved by the institutional review board or research ethics board at each participating site, and written informed consent was obtained from participants or their authorized representatives. The CGGA source collection was approved by the Beijing Tiantan Hospital Institutional Review Board, was conducted in accordance with the Declaration of Helsinki, and obtained written informed consent; the CGGA resource identifies collection protocol KY2013-017-01. The AMP-AD analysis used de-identified post-mortem multi-omics data from the AMP-AD Diverse Cohorts Study contributed by Emory University, Mayo Clinic, Mount Sinai, and Rush University and accessed as supplied analysis matrices. The source study reports approval by the institutional review board at each respective institution and consent from all participants or next of kin [28]; the supplied matrices did not retain a single cross-cohort protocol identifier, and none is asserted here.

The TCGA GBM/LGG analysis used de-identified coded genomic data from an existing public research resource and involved no participant recontact. TCGA source collections followed NCI and NHGRI human-subject protection, informed-consent, privacy, and data-access policies (https://www.cancer.gov/ccg/research/genome-sequencing/tcga/history/ethics-policies); the source studies describe the collection and genomic characterization procedures [23, 24]. The historical histological labels used here are an analysis endpoint and are not asserted to represent current WHO integrated diagnoses.

The VGH brain-tumour radiomics analysis used existing clinical DICOM images and MGMT metadata under University of British Columbia research ethics approval H20-02354 (“Brain Tumor Image Analysis”; principal investigator Stephen Yip). The requirement for informed consent was waived for this retrospective study, and secondary analysis was permitted. DICOM data were de-identified during restricted preprocessing before radiomics analysis; direct identifiers remained confined to restricted linkage records, and participant-level data are not redistributed.

### 4.14 Use of generative AI

OpenAI’s ChatGPT was used for coding assistance and grammar editing. The authors reviewed and verified all resulting changes and take full responsibility for the submitted work.

## 5 Data availability

Participant-level data are not redistributed; access requires the applicable source procedures. ADNI data are available to approved investigators through the LONI Image and Data Archive at https://ida.loni.usc.edu/login.jsp?project=ADNI&page=HOME. AMP-AD Diverse Cohorts data are available through the AD Knowledge Portal (https://adknowledgeportal.synapse.org/DataAccess;syn51732482; https://doi.org/10.7303/9618093) [40]. CGGA mRNAseq_325 data are available at https://www.cgga.org.cn/download.jsp, and TCGA GBM and lower-grade-glioma source data through the NCI Genomic Data Commons at https://portal.gdc.cancer.gov/. Redistribution permission has not been established for the provider-organized processed TCGA workbook; the public reproducibility repository supplies nonrestricted checksums and provenance manifests. VGH participant-level matrices, identifiers, images, masks, and prediction records cannot be redistributed because of participant-privacy and institutional ethics restrictions. Requests for controlled access should be directed to the corresponding author and remain subject to institutional and ethics approval; only non-identifying aggregate and audit materials are shared publicly.

## 6 Code availability

WrapEvoFS v0.2.0 is publicly available under the BSD 3-Clause License at https://github.com/junseoparkX/wrapevofs-package; the tagged release is available at https://github.com/junseoparkX/wrapevofs-package/releases/tag/v0.2.0. The verified wheel and source distribution are available from https://pypi.org/project/wrapevofs/0.2.0/; install with pip install wrapevofs==0.2.0. Nonrestricted manuscript source, analysis code, figure source data, checksums, provenance manifests, and aggregate audit materials are publicly available at https://github.com/junseoparkX/wrapevofs-manuscript-reproducibility; restricted participant-level data are not included.

## Acknowledgements

We thank Selina Parmar for providing the CGGA data, Ruihan Xu for providing the ADNI-derived and AMP-AD data, and Cecilia Liang for identifying, organizing, and sharing the processed TCGA GBM/LGG multi-omics dataset used in this study.

Data used in preparation of this article were obtained from the Alzheimer’s Disease Neuroimaging Initiative (ADNI) database. The investigators within ADNI contributed to the design and implementation of ADNI and/or provided data but did not participate in the analysis or writing of this report. Data collection and sharing for ADNI is funded by the National Institute on Aging (National Institutes of Health Grant U19AG024904); the grantee organization is the Northern California Institute for Research and Education. ADNI has also received support from the National Institute of Biomedical Imaging and Bioengineering, the Canadian Institutes of Health Research, and private-sector contributions facilitated by the Foundation for the National Institutes of Health, including AbbVie; Alzheimer’s Association; Alzheimer’s Drug Discovery Foundation; Araclon Biotech; BioClinica; Biogen; Bristol-Myers Squibb; CereSpir; Cogstate; Eisai; Elan Pharmaceuticals; Eli Lilly; EuroImmun; F. Hoffmann-La Roche and Genentech; Fujirebio; GE Healthcare; IXICO; Janssen Alzheimer Immunotherapy Research and Development; Johnson & Johnson Pharmaceutical Research and Development; Lumosity; Lundbeck; Merck; Meso Scale Diagnostics; NeuroRx Research; Neurotrack Technologies; Novartis Pharmaceuticals; Pfizer; Piramal Imaging; Servier; Takeda Pharmaceutical; and Transition Therapeutics. Data collection and sharing for the metabolomics component were also funded by the Alzheimer’s Disease Metabolomics Consortium through National Institute on Aging grants R01AG046171, RF1AG051550, and 3U01AG024904-09S4.

The results published here are in whole or in part based on data obtained from the AD Knowledge Portal. The data available through the portal would not be possible without the participation of research volunteers and the contributions of collaborating researchers. Data generation for the AMP-AD Diverse Cohorts Study was supported by National Institutes of Health grants U01AG046139, U01AG046170, U01AG061357, U01AG061356, U01AG061359, and R01AG067025.

## Author contributions

J.P. performed conceptualization, methodology, software development, validation, formal analysis, investigation, data curation, DICOM de-identification, radiomics data generation, visualization, and writing of the original draft. L.J.F. and H.Z. provided supervision; H.Z. also provided project administration and serves as corresponding author, and L.J.F. serves as co-corresponding author. M.F.H., E.S., and S.C. contributed the patient DICOM dataset and associated clinical metadata for the VGH brain-tumour radiomics cohort, supported the ethics and consent documentation, and contributed to resources and data curation. S.C. additionally coordinated collaboration with the VGH team. J.P., L.J.F., M.F.H., E.S., and H.Z. contributed to review and editing. All authors reviewed and approved the final manuscript and take responsibility for the work.

## References

[1] Ambroise, C. & McLachlan, G. J. Selection bias in gene extraction on the basis of microarray gene-expression data. Proc. Natl Acad. Sci. USA 99, 6562–6566 (2002).

[2] He, Z. & Yu, W. Stable feature selection for biomarker discovery. Comput. Biol. Chem. 34, 215–225 (2010).

[3] Ein-Dor, L., Kela, I., Getz, G., Givol, D. & Domany, E. Outcome signature genes in breast cancer: is there a unique set? Bioinformatics 21, 171–178 (2005).

[4] Vaxrma, S. & Simon, R. Bias in error estimation when using cross-validation for model selection. BMC Bioinformatics 7, 91 (2006).

[5] Steyerberg, E. W. & Harrell, F. E. J. Prediction models need appropriate internal, internal-external, and external validation. J. Clin. Epidemiol. 69, 245–247 (2016).

[6] Cawley, G. C. & Talbot, N. L. C. On over-fitting in model selection and subsequent selection bias in performance evaluation. J. Mach. Learn. Res. 11, 2079–2107 (2010).

[7] Kapoor, S. & Narayanan, A. Leakage and the reproducibility crisis in machine-learning-based science. Patterns 4, 100804 (2023).

[8] Pedregosa, F. et al. Scikit-learn: machine learning in python. J. Mach. Learn. Res. 12, 2825–2830 (2011).

[9] Guyon, I., Weston, J., Barnhill, S. & Vapnik, V. Gene selection for cancer classification using support vector machines. Mach. Learn. 46, 389–422 (2002).

[10] Siedlecki, W. & Sklansky, J. A note on genetic algorithms for large-scale feature selection. Pattern Recognit. Lett. 10, 335–347 (1989).

[11] Goldberg, D. E. Genetic Algorithms in Search, Optimization, and Machine Learning (Addison-Wesley, Reading, MA, 1989).

[12] Meinshausen, N. & Bühlmann, P. Stability selection. J. R. Stat. Soc. Series B Stat. Methodol. 72, 417–473 (2010).

[13] Abeel, T., Helleputte, T., Van de Peer, Y., Dupont, P. & Saeys, Y. Robust biomarker identification for cancer diagnosis with ensemble feature selection methods. Bioinformatics 26, 392–398 (2010).

[14] Marx, C., Calmon, F. & Ustun, B. Predictive multiplicity in classification. In Proceedings of the 37th International Conference on Machine Learning, vol. 119 of Proceedings of Machine Learning Research, 6765–6774 (PMLR, 2020). URL https://proceedings.mlr.press/v119/marx20a.html.

[15] Fisher, A., Rudin, C. & Dominici, F. All models are wrong, but many are useful: Learning a variable’s importance by studying an entire class of prediction models simultaneously. Journal of Machine Learning Research 20, 1–81 (2019). URL https://www.jmlr.org/papers/v20/18-760.html.

[16] Xue, B., Cervante, L., Shang, L., Browne, W. N. & Zhang, M. Multi-objective evolutionary algorithms for filter based feature selection in classification. International Journal on Artificial Intelligence Tools 22, 1350024 (2013).

[17] Moscovich, A. & Rosset, S. On the cross-validation bias due to unsupervised preprocessing. J. R. Stat. Soc. Series B Stat. Methodol. 84, 1474–1502 (2022).

[18] Nogueira, S., Sechidis, K. & Brown, G. On the stability of feature selection algorithms. J. Mach. Learn. Res. 18, 1–54 (2018).

[19] Chen, T. & Guestrin, C. Xgboost: A scalable tree boosting system. In Proceedings of the 22nd ACM SIGKDD International Conference on Knowledge Discovery and Data Mining, 785–794 (2016).

[20] Kursa, M. B. & Rudnicki, W. R. Feature selection with the boruta package. J. Stat. Softw. 36, 1–13 (2010).

[21] Breiman, L. Random forests. Mach. Learn. 45, 5–32 (2001).

[22] Zou, H. & Hastie, T. Regularization and variable selection via the elastic net. J. R. Stat. Soc. Series B Stat. Methodol. 67, 301–320 (2005).

[23] The Cancer Genome Atlas Research Network. Comprehensive genomic characterization defines human glioblastoma genes and core pathways. Nature 455, 1061–1068 (2008).

[24] The Cancer Genome Atlas Research Network. Comprehensive, integrative genomic analysis of diffuse lower-grade gliomas. N. Engl. J. Med. 372, 2481–2498 (2015).

[25] Ceccarelli, M. et al. Molecular profiling reveals biologically discrete subsets and pathways of progression in diffuse glioma. Cell 164, 550–563 (2016).

[26] Weiner, M. W. et al. The alzheimer’s disease neuroimaging initiative: progress report and future plans. Alzheimers Dement. 6, 202–211.e7 (2010).

[27] Petersen, R. C. et al. Alzheimer’s disease neuroimaging initiative (adni): clinical characterization. Neurology 74, 201–209 (2010).

[28] Reddy, J. S., Heath, L., Vander Linden, A., Allen, M. et al. Bridging the gap: Multi-omics profiling of brain tissue in alzheimer’s disease and older controls in multi-ethnic populations. Alzheimer’s & Dementia 20, 7174–7192 (2024).

[29] Xiong, Y. et al. Multi-dimensional omics characterization in glioblastoma identifies the purity-associated pattern and prognostic gene signatures. Cancer Cell Int. 20, 37 (2020).

[30] Li, X., Morgan, P. S., Ashburner, J., Smith, J. & Rorden, C. The first step for neuroimaging data analysis: DICOM to NIfTI conversion. J. Neurosci. Methods 264, 47–56 (2016).

[31] Bouget, D. et al. Raidionics: an open software for pre- and postoperative central nervous system tumor segmentation and standardized reporting. Sci. Rep. 13, 15570 (2023).

[32] van Griethuysen, J. J. M. et al. Computational radiomics system to decode the radiographic phenotype. Cancer Res. 77, e104–e107 (2017).

[33] Zwanenburg, A. et al. Assessing robustness of radiomic features by image perturbation. Sci. Rep. 9, 614 (2019).

[34] Zwanenburg, A., Vallières, M., Abdalah, M. A. et al. The image biomarker standardisation initiative: Standardized quantitative radiomics for high-throughput image-based phenotyping. Radiology 295, 328–338 (2020).

[35] Papaspiliopoulos, O., Roberts, G. O. & Sköld, M. A general framework for the parametrization of hierarchical models. Stat. Sci. 22, 59–73 (2007).

[36] Hédou, J., Marić, I., Bellan, G. et al. Discovery of sparse, reliable omic biomarkers with stabl. Nat. Biotechnol. 42, 1581–1593 (2024).

[37] Park, T. & Casella, G. The bayesian lasso. J. Am. Stat. Assoc. 103, 681–686 (2008).

[38] Spector, A. & Janson, L. Controlled discovery and localization of signals via bayesian linear programming. J. Am. Stat. Assoc. 120, 460–471 (2025).

[39] pyblip developers. pyblip software repository and documentation. https://github.com/amspector100/pyblip (2026). Accessed July 26, 2026.

[40] AD Knowledge Portal. Diverse cohorts study. Synapse (2026). URL 10.7303/9618093.

