## Supplementary Information for "WrapEvoFS enables auditable feature compression with regret-constrained representative locking"

##### Author email addresses

| | | | |
|---|---|---|---|
| Junseo Park | | Emma Schmidt | |
| Shuxin Chi | | Mostafa Fatehi Hassanabad | |
| Leonard J. Foster | | Huan Zhong | |

##### Correspondence


[‡]Data used in preparation of this article were obtained from the Alzheimer's Disease Neuroimaging Initiative (ADNI) database (adni.loni.usc.edu). The investigators within ADNI contributed to the design and implementation of ADNI and/or provided data but did not participate in the analysis or writing of this report. The complete investigator list is available in the ADNI acknowledgement list.

##### Contents

Supplementary Methods; Supplementary Figures S1–S28; Supplementary Tables S1–S28; Supplementary References.

This Supplementary Information provides the complete algorithm, formal locking guarantees, computational complexity, archived and secondary analyses, one-time post-freeze AMP-AD evaluation details, a coherent CGGA benchmark, a reduced-budget saved-bank nested sensitivity, a controlled candidate-bank locking simulation, a fully nested TCGA GBM/LGG analysis, a VGH brain-tumour radiomics internal demonstration, and machine-readable audit correspondence. The package preserves the original algorithms as legacy-compatible modes, and archived results are not relabeled as corrected-objective results. Complete candidate- and metric-level audits are supplied as machine-readable CSV files with analysis provenance. Supplementary Figures S21 to S22 provide descriptive tuned-evaluator sensitivity and computational provenance and are not used for a core claim.

### S1 Supplementary Methods

#### S1.1 Complete WrapEvoFS algorithm

**Complete development-only WrapEvoFS search and regret-constrained representative locking.**

**Input.** Development data $(X_D, y_D)$; Direct candidates $\mathcal{C}_m$; RFECV configuration; GA mode and base seed $\xi_0$; population size $N$; generations $G$; runs $R$; elite count $e$; crossover probability $p_c$; per-bit mutation probability $p_\mu$; initial off-ratio $\rho_0$; locking metric and seed; tolerance mode and $\delta$.

**Output.** RFECV target $k_m^*$; locked set $F_m^{\text{lock}}$; complete audit $A_m$.

Fit preprocessing and Direct selection on development data only.

Evaluate the eligible RFECV path and select $k_m^*$; resolve score ties toward the smaller count.

Construct fixed stratified development folds for GA evaluation. Use zero-based indices $(r, g, i, \ell)$ in the seed definitions below.

**For each** independent run $r = 0, \ldots, R-1$, initialize the genetic-operator generator with seed $\xi_0 + r$.

  **For each** of $N$ initial chromosomes, set exactly $\lfloor \rho_0 p_m \rfloor$ entries to zero and the remainder to one, uniformly shuffle their positions, and activate one uniformly sampled feature if an empty mask would otherwise result.

  **For each** generation $g = 0, \ldots, G-1$:

    **For each** chromosome at population-list index $i$, compute its $K_G$ fold scores with random-forest seed $\xi_{rgi\ell} = \xi_0 + 10000r + 100g + 1009i + \ell$ in fold $\ell$.

    Compute the mean base score, target deviation, penalty, raw objective, and zero-truncated legacy fitness.

    **If corrected-objective mode:** rank and update run best by raw objective, base score, smaller deviation, smaller count, and stable mask hash.

    **If legacy-compatible mode:** reproduce zero-truncated ranking and update behavior.

    Copy the first $e$ ranked chromosomes unchanged to the next generation.

    For the remaining $N - e$ positions, compute shifted proportional weights in corrected-objective mode (zero-truncated nonnegative weights in legacy-compatible mode) and normalize them to parent probabilities.

    **If** weights are nonfinite, effectively equal, or have invalid or effectively zero total, set all parent probabilities to $1/N$ and record the fallback.

    Sample $N - e$ parent indices independently with replacement and pair the sampled parents in list order.

    For each pair, perform one-point crossover with probability $p_c$; when $p_m > 2$, sample the cut uniformly from $\{1, \ldots, p_m - 2\}$, otherwise copy both parents.

    Independently flip every child bit with probability $p_\mu$; if a child is empty, activate one uniformly sampled feature.

    Form the next size-$N$ population from the $e$ elites followed by mutated children; export generation diagnostics and warnings for zero truncation, deviation, sampling fallback, and mask collapse.

  **End for** generation; retain one run-best candidate.

**End for** run.

Construct one fixed stratified locking-fold partition and re-score every retained run-best candidate with the same locking metric, fold partition, and random-forest hyperparameters, using its recorded run-specific estimator seed, to obtain $L_r$ and fold scores.

Encode every candidate in the common ordered feature universe, compute its stable mask identifier, and compute $R_r = L_{\text{best}} - L_r$ (plus configured sensitivity quantities).

For the primary absolute rule, form $E_\delta = \{r : R_r \leq \delta\}$. Prespecified relative and best-run-SE-scaled modes form their corresponding sensitivity pools under their own declared thresholds. Never add a candidate outside the active threshold.

**If** $|E_\delta| = 1$, select its sole candidate.

**Otherwise,** compute pairwise Jaccard similarity and each candidate's mean Jaccard within $E_\delta$.

25 Select the medoid by larger mean Jaccard, larger $L_r$, smaller feature count, and lexicographically smaller stable mask hash; use source-run ID only as post-selection provenance among exact duplicate masks.

26 Assert that the selected candidate is in $E_\delta$ and satisfies the exact declared threshold; export canonical masks and hashes, duplicate multiplicity and provenance, scores, fold scores, regrets, eligibility, Jaccard values, tie path, seeds, version, and configuration hash.

27 **Return** $(F_m^{\text{lock}}, k_m^*, A_m)$. Held-out data are not inputs and are used only for one-time evaluation after the feature set, settings, and final model-development procedure are locked.

---

The algorithm makes the leakage boundary operational: held-out data are absent from search, candidate rescoring, eligibility, representative locking, and audit export. The main text gives the scientific definitions; the following sections provide the formal and engineering details. The executable v0.2.0 wrapper, RFECV, genetic-operator, evaluator, locking, and randomness specifications are consolidated in Supplementary Table S24; archived dataset-level metric and cap overrides remain in Supplementary Table S27.

##### S1.2 Complete notation and objective definitions

This section specifies the quantities used in candidate generation, rescoring, and representative locking. For Direct branch $m$, let $\mathcal{C}_m$ be the pre-GA candidate set with size $p_m$. RFECV chooses the smallest count attaining the largest mean development-fold score among counts permitted by the configured cap:

$$k_m^* = \min\left(\arg\max_{k\in\mathcal{K}_m^{\text{elig}}} \overline{S}_{R,m}(k)\right). \tag{1}$$

For selected set $F$, compression relative to the Direct set is

$$C(F) = 1 - \frac{|F|}{p_m}. \tag{2}$$

Every retained candidate is evaluated on the same $K_L$ fixed stratified folds. With $s_{r\ell}^L$ denoting the locking-metric score for candidate $r$ on fold $\ell$, its mean locking score, empirical regret, and strict eligible pool are

$$L_r = \frac{1}{K_L}\sum_{\ell=1}^{K_L} s_{r\ell}^L, \tag{3}$$

$$R_r = L_{\text{best}} - L_r, \qquad L_{\text{best}} = \max_j L_j, \tag{4}$$

$$E_\delta = \{r : R_r \le \delta\}. \tag{5}$$

The held-out contrast, calculated only after locking, is

$$\Delta_{\text{heldout}} = M(F^{\text{lock}}) - M(\mathcal{C}_m). \tag{6}$$

A nonempty binary chromosome $\mathbf{z} \in \{0,1\}^{p_m}$ represents feature set $F_m(\mathbf{z})$. Using $K_G$ fixed stratified folds and chromosome-level seed $\xi$, the base search score and size-penalized raw objective are

$$\overline{S}_{G,m}(\mathbf{z};\xi) = \frac{1}{K_G}\sum_{\ell=0}^{K_G-1} s_{m\ell}^G(\mathbf{z};\xi), \tag{7}$$

$$Q_m(\mathbf{z};\xi) = \overline{S}_{G,m}(\mathbf{z};\xi) - \lambda\,\big||F_m(\mathbf{z})| - k_m^*\big|. \tag{8}$$

The legacy-compatible objective truncates negative values,

$$\Phi_m^{\text{legacy}}(\mathbf{z};\xi) = \max\{0, Q_m(\mathbf{z};\xi)\}, \tag{9}$$

whereas the corrected mode uses $Q_m$ directly for ranking, elite selection, and run-best updating. For proportional parent sampling only, it maps each finite population objective $Q_i$ to the nonnegative weight below, where $Q_{\min}$ is the minimum finite objective in the evaluated population:

$$w_i = Q_i - Q_{\min} + \epsilon, \qquad \epsilon = 10^{-12}, \tag{10}$$

and samples with probability

$$\pi_i = \frac{w_i}{\sum_j w_j}. \tag{11}$$

Here $N$ is the population size. Nonfinite objectives receive zero weight. Uniform sampling, $\pi_i = 1/N$, is recorded when no objective is finite, the finite-objective range is at most $\epsilon$, or the weight total is invalid or effectively zero.

For a nonsingleton eligible pool, candidate $r$'s mean Jaccard similarity is

$$\overline{J}_r = \frac{1}{|E_\delta| - 1} \sum_{\substack{s \in E_\delta \\ s \neq r}} \frac{|F_r \cap F_s|}{|F_r \cup F_s|}. \tag{12}$$

Candidates are ordered by larger $\overline{J}_r$, larger $L_r$, smaller $|F_r|$, and lexicographically smaller canonical stable-mask hash. If $\widehat{r}$ is first in that ordering, the locked feature set is

$$F^{\text{lock}} = F_{\widehat{r}}. \tag{13}$$

These equations define the implemented quantities; the propositions below establish finite-bank feasibility and deterministic selection.

##### S1.3 Stochastic-agreement summary

For five saved GA runs, Nogueira stability was computed from the run-by-candidate binary selection matrix as one minus the mean observed per-feature sample variance divided by the Bernoulli variance expected from the mean selected proportion [1]. The candidate-universe size was the branch-specific Direct set size. With only five runs, these estimates are noisy and characterize stochastic agreement on one fixed development sample rather than stability under participant resampling.

##### S1.4 Formal properties of regret-constrained medoid locking

Let the finite nonempty retained candidate bank be a multiset indexed by $r \in \mathcal{R}$. Candidate $r$ has a nonempty feature set $F_r$ and finite larger-is-better locking score $L_r$. All candidates use one common ordered canonical universe $U = (u_1, \ldots, u_p)$ with unique feature names. The canonical mask is $z_{rj} = \mathbf{1}\{u_j \in F_r\}$, and its stable identifier is $H_r = \text{hash}(\text{uint8}(z_r).\,\text{tobytes}())$. We assume $\delta \geq 0$, well-defined Jaccard similarity, deterministic canonical encoding, and collision resistance of the stable identifier. Define

$$L_{\text{best}} = \max_{r \in \mathcal{R}} L_r, \qquad R_r = L_{\text{best}} - L_r, \qquad E_\delta = \{r \in \mathcal{R} : R_r \leq \delta\}.$$

For $|E_\delta| > 1$ define

$$\overline{J}_r = \frac{1}{|E_\delta| - 1} \sum_{\substack{s \in E_\delta \\ s \neq r}} \frac{|F_r \cap F_s|}{|F_r \cup F_s|}.$$

Selection is restricted to $E_\delta$ and ordered by larger $\overline{J}_r$, larger $L_r$, smaller $|F_r|$, and lexicographically smaller $H_r$. A singleton is selected directly and has undefined mean Jaccard. Duplicate masks are retained as multiple voting candidates; exact duplicates remaining tied after the hash represent one scientific feature set. A source run ID is then retained only as a provenance label, not as a scientific tie-break.

**Proposition 1 (nonempty eligibility).** For every finite nonempty bank and $\delta \geq 0$, $E_\delta$ is nonempty. *Proof.* At least one $r^*$ attains $L_{\text{best}}$, giving $R_{r^*} = 0 \leq \delta$ and hence $r^* \in E_\delta$. *QED.*

**Proposition 2 (strict empirical regret feasibility).** The selected record $\widehat{r}$ belongs to $E_\delta$ and satisfies $R_{\widehat{r}} \leq \delta$. *Proof.* A singleton rule returns the sole eligible record. For a nonsingleton, the medoid optimization and every score, size, hash, and duplicate-provenance tie path are restricted to $E_\delta$. No path can admit an ineligible record. Thus $\widehat{r} \in E_\delta$, which is equivalent to the stated inequality. *QED.*

**Proposition 3 (zero tolerance).** $E_0 = \{r : L_r = L_{\text{best}}\}$. A unique score maximizer is selected; if several candidates attain $L_{\text{best}}$, the rule selects the Jaccard medoid of the complete zero-regret tied pool under the canonical tie-break. *Proof.* Since $R_r \geq 0$, $R_r \leq 0$ iff $R_r = 0$ iff $L_r = L_{\text{best}}$. The singleton or nonsingleton conclusion follows. Therefore $\delta = 0$ does not always produce a singleton. *QED.*

**Proposition 4 (all-candidate limit).** If $\delta \geq \max_r R_r$, every candidate is eligible and the method is the full-bank Jaccard medoid under the same tie-break. *Proof.* The condition gives $R_r \leq \delta$ for all $r$, hence $E_\delta = \mathcal{R}$. Substitute this equality into the medoid definition. *QED.*

**Proposition 5 (determinism).** Given identical canonical masks, scores, source provenance, tolerance, larger-is-better orientation, and configuration, the selected feature set and canonical audit are deterministic. *Proof.* Maxima, regrets, exact eligibility, Jaccard values, and means are deterministic functions of the inputs. Canonical masks and stable identifiers are deterministic byte transformations. The tie key totally orders distinct masks under the collision-resistance assumption; exact duplicates have the same scientific output and a deterministic post-selection provenance label. Audit rows and configuration fields are canonically ordered. *QED.*

The digest is a stable ordering device, not a probabilistic scientific score. Collision resistance is an engineering assumption and is not the source of the regret bound, which follows from restricting selection to $E_\delta$.

**Proposition 6 (candidate-order permutation invariance).** Permuting candidate input records, without changing their content, provenance, or multiplicity, does not alter the selected feature set or canonical audit. *Proof.* The permutation preserves the score multiset, regrets, $E_\delta$, and every eligible-pool Jaccard sum. The remaining tie key depends only on content-derived quantities. Exact duplicate records yield the same feature set, provenance normalization is independent of row order, and canonical serialization restores the same audit order. *QED.*

Candidate content drives scientific selection, source-run provenance is kept for traceability, and input-row order is discarded. Because duplicates remain voting records, multiplicity is part of the bank: adding or removing a duplicate can change the medoid, whereas permuting it cannot.

**Proposition 7 (distinction from alternative rules).** The highest-score rule guarantees zero empirical regret but does not optimize representativeness. The unrestricted medoid optimizes full-bank mean Jaccard but can incur regret as large as $\max_r R_r$. A legacy top-$k$ medoid uses a rank cutoff, whose metric-scale score gap is unrestricted. Regret-constrained locking first imposes $R_r \leq \delta$ and then optimizes eligible-pool representativeness. These are candidate-bank statements, not claims of globally optimal feature selection.

**Claim boundary.** $R_r$ is an empirical development-CV score gap, not expected predictive risk, generalization error, or statistical uncertainty. The propositions do not establish predictive superiority, unbiased generalization, external validity, participant-resampling or biomarker stability, or clinical utility.

##### S1.5 ADNI archived-bank recovery and retrospective re-locking

The archived ADNI Drive source retained, for each Direct branch, five run-best binary masks, their feature tables, and development-CV summaries. Before re-locking, the audit required exactly five nonempty masks, a common branch-specific universe, finite scores, exact agreement between each binary mask and its decoded feature set, and reproduction of the archived legacy top-three medoid. The scientific larger-is-better locking score was the archived `cv_balanced_accuracy_mean`; GA fitness arrays and later weighted one-vs-rest AUROC summaries were not substituted for it. The recovered mask matrices had shapes $5 \times 63$, $5 \times 100$, and $5 \times 48$ for SVM-L1, XGBoost, and Boruta-RF, respectively, and contained no duplicate masks.

No GA, Direct selection, RFECV, candidate scoring, model fitting, or held-out evaluation was rerun. Absolute eligibility at $\delta = 0.01$ and the canonical content-derived tie-break were applied to the verified banks. All three eligible pools were singletons, so the highest-locking-score candidate was selected with empirical regret zero and neither a Jaccard-medoid nor stable-hash tie was reached. The selected GA ranks were 4, 2, and 1 for SVM-L1, XGBoost, and Boruta-RF. Exhaustive permutation of the five input records gave the same selected feature set in all 120 orders for each branch. Supplementary Table S8 reports the complete candidate audit; machine-readable scores, masks, pairwise Jaccard values, source provenance, and validation checks accompany the reproducibility package.

##### S1.6 VGH brain-tumour radiomics internal demonstration

The VGH brain-tumour radiomics analysis began from a heterogeneous clinical DICOM export handled in a restricted environment under UBC REB H20-02354 (“Brain Tumor Image Analysis”). A header-only preflight grouped records by patient, study, series, and instance identity; prespecified sequence and geometry rules selected one eligible T1ce series without access to MGMT. Selected series were de-identified and converted read-only to pseudonymized NIfTI with `dcm2niix` v1.0.20260724 [2]. RaidionicsRADS v1.3.2 and RaidionicsSeg v1.5.2 generated automatic brain and tumor-core candidate regions in T1ce space [3]. Empty masks, image–mask geometry mismatch, nonfinite voxels, out-of-bounds masks, and execution failure were hard technical failures. Morphology and acquisition findings were retained as review flags rather than automatic exclusions; no expert contour correction or adjudication was performed.

PyRadiomics v3.0.1 [4] used intensity normalization (scale 100), 1-mm isotropic B-spline image resampling, nearest-neighbor mask resampling, bin width 25, geometry tolerance $10^{-5}$, minimum three-dimensional and 64-voxel regions, pre-cropping, and a five-voxel pad. First-order, shape, GLCM, GLRLM, GLSZM, GLDM, and NGTDM features were enabled for Original, Wavelet, Laplacian-of-Gaussian (1–5 mm), Gradient, Square, SquareRoot, Logarithm, and Exponential image families. This produced two frozen supervised matrices with identical participant order and outcomes: a full 1,781-feature T1ce tumor-core space and an order-preserving 1,346-feature subset. The subset definition was frozen label-independently in a deterministic 15-participant pilot using features extracted from original, 1-mm-eroded, and 1-mm-dilated automatic tumor-core masks. Retention required ICC(2,1) at least 0.85 and median relative range at most 0.25. Outcome linkage occurred only after imaging selection, conversion, segmentation QC, configuration freezing, extraction, and stability filtering. Supplementary Table S28 records the complete non-identifying construction boundary. The manuscript package does not redistribute pilot-level perturbation measurements, scanner/site covariates, repeat segmentations, masks, or patient-level QC records, so those upstream checks cannot be independently recomputed from the submission bundle.

Both feature spaces used the same stratified 70:30 split (random state 42): 137 development participants and 60 held-out participants. For each space, SVM-L1, XGBoost, and Boruta-RF generated Direct candidates; five-fold RFECV supplied a soft target; and seeds 42–46 each completed a 50-generation, 50-individual GA using five-fold ROC-AUROC, the corrected untruncated shifted-linear objective, and a target-deviation penalty of 0.015. All 30 GA completion records stated that held-out data were unused and passed checksum, mask-hash, generation-count, and finite-score audits. The five run-best masks in each of six banks were then rescored by one fixed five-fold development-only random-forest evaluator and locked using the strict absolute regret constraint $\delta = 0.01$. The frozen package policy retained exact duplicate masks as multiple voting candidates while recording their common scientific feature set and source-run provenance separately.

All six development locks were written before held-out evaluation. A fixed 500-tree random forest was then fitted on the entire development partition for Direct, RFECV, and locked feature sets and applied once to the held-out partition. AUROC and AUPRC intervals and paired AUROC differences used 2,000 class-stratified bootstrap resamples. The second feature space reuses the same participants, outcomes, and split, so it is a feature-definition sensitivity analysis rather than an external validation. Five-seed agreement is likewise stochastic agreement on one development sample, not participant-resampling or biomarker stability. Participant-level predictions are retained outside the manuscript package; only non-identifying aggregate summaries and locking audits are distributed.

##### S1.7 AMP-AD post-freeze evaluation and Rush source recovery

The 24 corrected-objective AMP-AD development locks were enumerated in a frozen inventory containing the held-out center, branch, cap, selected run, selected feature list, fixed locking score, empirical regret, eligible-pool size, tie path, and source paths. Before any held-out-center file was read, the evaluation protocol fixed the primary and secondary metrics, final random-forest estimator, run-derived seeds, archived comparators, 2,000-resample paired bootstrap, and reporting rule. The protocol and inventory hashes were then checked at evaluation entry, and a completion sentinel prevented a second execution. All 24 conditions were reported irrespective of direction. This temporal freeze prevents held-out-guided selection within the analysis but does not make the evaluation prospective clinical validation or create 24 independent cohorts.

The authoritative Rush source was a Google Drive results folder rather than an original ZIP archive. All 30 run-best feature CSV files, the run-level summary, and the locking summary were recovered with their Drive file IDs and URLs. A local provenance ZIP was reconstructed without claiming it to be the original archive. Regret-constrained locking reproduced all six archived Rush selected runs and all six masks used by the frozen held-out evaluation; maximum selected empirical regret was zero, no exact duplicate masks occurred, and no selection reached the stable-hash tie stage. Recovery completed the corrected-objective cross-lock and duplicate audits for all 24 AMP-AD conditions.

##### S1.8 CGGA coherent benchmark and nested saved-bank re-locking

For the coherent benchmark, the raw CGGA expression and clinical files were reconstructed using the archived fixed split and train-fitted missingness, minimum-positive imputation, log transformation, and zero-variance filtering. Participant order, labels, and every value in the archived compact development and held-out matrices were required to match before analysis continued. Six existing development-derived signatures per branch were compared: RFECV-only, size-matched Elastic Net, highest fixed development-CV candidate, legacy top-three medoid, unrestricted full-bank medoid, and regret-constrained medoid. Every signature used the same 500-tree random forest and the same 92-sample held-out set. The 2,000 class-stratified bootstrap samples were common across methods, enabling paired regret-constrained-medoid contrasts. No feature-selection stage was rerun.

The nested saved-bank sensitivity reused five outer-fold candidate banks from an earlier development-only CGGA analysis that had been excluded from the primary release because its search budget was reduced to 20 individuals and 12 generations. Each bank was re-locked with the regret-constrained medoid at absolute $\delta = 0.01$ and the canonical tie-break; the GA, Direct selection, RFECV, candidate score computation, and original 92-sample held-out evaluation were not rerun. A fixed 500-tree random forest was fitted within each outer training fold using the newly selected existing candidate, and OOF predictions were summarized across 214 development participants. The regret-constrained medoid changed all five selections and reduced OOF AUROC relative to the legacy top-three medoid. This result is retained as a limitation-oriented sensitivity and does not replace the main fixed-split CGGA experiment.

##### S1.9 Fully nested TCGA GBM/LGG analysis

The TCGA analysis began from a provider-organized common-participant workbook. The included sheets contained 491 participants, 20,118 processed RNA-seq predictors, 5,000 cleaned SCNV predictors, and four historical histological labels. The data provider confirmed in writing that the 5,000 SCNV predictors were selected only by unsupervised variance filtering across the common multi-omics participants and that histological class labels were not used. A separate label-blind reconstruction ranked all 24,776 features in the uncleaned `SCNV_Common` sheet by sample variance, recovered 4,999 of the 5,000 supplied cleaned features, and found that the selected and excluded features at the cutoff had exactly equal variance. Thus, the sole reconstruction ambiguity was a deterministic tie at rank 5,000, not evidence of outcome-dependent screening. Each predictor retained an `RNA::` or `SCNV::` prefix. Clinical columns, RPPA, and the uncleaned SCNV sheet were excluded. The source values were not transformed during extraction. This provider-supplied reduction occurred before the present outer-CV experiment; the cleaned participant-by-feature matrix and labels were checksum-locked before modeling.

Two shuffled stratified five-fold outer-CV repeats used random states 20260810 and 20260811. Median imputation, zero-variance filtering, branch-specific Direct screening, deterministic post-Direct capping at 100 features, RFECV, GA search, fixed candidate rescoring, and locking were fitted exclusively within each outer-training fold. The three Direct branches were SVM-L1, XGBoost, and Boruta-RF. RFECV used five-fold macro one-vs-rest AUROC and branch caps of 15, 20, and 10, respectively. Each of the 30 repeat–fold–branch banks comprised five GA runs with seeds 42–46, 50 individuals, 50 generations, the untruncated shifted-linear objective, and a target-deviation penalty of 0.015. The locking evaluator used a 500-tree random forest, five fixed outer-training folds, larger-is-better macro one-vs-rest AUROC, strict absolute eligibility at $\delta = 0.01$, and the canonical content-derived tie-break. Exact duplicates would have retained multiplicity as voting candidates with source-run provenance stored separately; no within-bank duplicate masks occurred.

The complete candidate audit in Supplementary Fig. S5 plots all 150 candidates against empirical regret and Jaccard distance from the canonical highest-locking-score mask. Independent reconstruction verified every highest-locking-score mask, eligible pool, selected mask, selected regret, pairwise Jaccard, and mean Jaccard to numerical precision. All 30 selected regrets met the tolerance. Six pools were singletons. All five two-candidate pools were structural mean-Jaccard ties and retained the highest-locking-score mask. The regret-constrained medoid differed from the highest-locking-score mask in 11 banks, all with at least three eligible candidates and positive mean-Jaccard gain.

Candidate agreement and participant-partition sensitivity are expanded in Supplementary Fig. S6. Nogueira stability was computed separately within each outer-training bank on that bank's fixed post-Direct universe; panel a reports the unweighted mean of the five fold-level estimates for each repeat–branch combination rather than a pooled 25-mask coefficient. A small negative finite-sample value is valid. Outer-fold pairwise similarities share folds and are dependent, so their distributions are descriptive. Cross-branch Jaccard was not computed because the branch-specific candidate universes differ. Cardinality similarity was defined as $\min(k_1, k_2)/\max(k_1, k_2)$. SCNV composition largely reflects the upstream Direct universes and is not interpreted as biological modality preference.

Six signatures were evaluated within each repeat–fold–branch condition: Direct, RFECV-only, the highest-locking-score candidate, the legacy top-three medoid, the unrestricted full-bank medoid, and the regret-constrained medoid. All used the same 500-tree random forest and common deterministic seed $50000+1000\times\text{repeat}+100\times\text{fold}+10\times\text{branch index}$. A guarded common-seed refresh reused only the already selected feature sets and reran no GA, Direct, RFECV, or adaptive selection. All 150 GA jobs remained unchanged; 180 final method fits produced 17,676 participant–method outer-prediction records, and all completion and prediction checksums passed. Repeated-OOF metrics and the complete comparator audit are shown in Supplementary Fig. S7. Confidence intervals and paired differences used 2,000 participant-clustered, class-stratified bootstrap resamples that retained both repeat-specific OOF predictions for each sampled participant. The analysis estimates repeated internal OOF performance and participant-partition sensitivity in one TCGA-derived cohort; it is not independent external validation. The claim boundary in Supplementary Methods S1.4 applies; modern WHO diagnostic validity and causal biology were not assessed.

A checksum-frozen selector-layer extension reused the same 30 verified post-Direct outer-training universes and existing common-seed WrapEvoFS predictions. It did not rerun Direct, RFECV, GA search, candidate rescoring, or locking. For each branch and outer fold, multinomial Elastic Net used a training-fitted standard scaler, balanced class weights, the SAGA solver, and five-fold inner-CV macro one-vs-rest AUROC. The fixed grid crossed $C \in \{0.0003, 0.001, 0.003, 0.01, 0.03, 0.1, 0.3\}$ with $l_1$ ratios 0.25, 0.50, 0.75, and 1.00. Exact score ties favored smaller $C$ and then larger $l_1$ ratio. Native support was the union of nonzero class coefficients. The matched-cardinality sensitivity ranked features by maximum absolute standardized coefficient, with canonical feature name resolving exact ties; any matched positions with zero coefficients were counted and were not described as native Elastic-Net support.

Subsampling stability selection reused the chosen Elastic-Net configuration in 50 class-stratified half-samples without replacement. Native support required selection frequency at least 0.80; the matched-cardinality sensitivity ranked by frequency, mean absolute coefficient, and canonical feature name. This descriptive procedure did not claim formal per-family error control. Five prespecified random banks per condition contained five distinct, uniformly drawn masks with the same size as the existing regret-constrained-medoid signature. Each bank was evaluated with the fixed 25-fit locking evaluator and the same absolute $\delta = 0.01$ rule. Thus, the random control matched bank size and locking-evaluation budget, not full GA compute. Feature sets and hashes were written before outer-test values or outcomes were opened; the condition-specific final random-forest seed was shared by every comparator. All 30 conditions and output checksums passed validation. Extended results are shown in Supplementary Fig. S8 and Supplementary Table S6. Because the parent TCGA outcomes had already been analyzed and the new selectors were conditional on post-Direct universes, this is a retrospective nested selector-layer sensitivity rather than a prospective or standalone full-pipeline benchmark.

##### S1.10 Controlled candidate-bank locking simulation

The controlled simulation isolated finite-bank locking under a hidden utility that was unavailable to each locking rule. The canonical universe contained $p = 1000$ features arranged as 20 latent signal groups with one canonical feature and four proxies per group. Candidate sets contained approximately 20 features. For candidate set $F$, hidden utility was

$$T(F) = \text{clip}\left(0.84 + 0.05G(F) - 0.0005I(F) - 0.00025\,||F| - 20|\,,\ 0.75,\ 0.90\right), \tag{14}$$

where $G(F)$ is mean group-aware recovery and $I(F)$ is the number of selected features outside the 20 signal groups. A selected canonical member received group credit 1; proxies received credit 0, 0.6, or 0.9 under the none, moderate, or strong proxy conditions. The observed larger-is-better locking score was $L = T + \epsilon$, where $\epsilon$ followed the prespecified Gaussian distribution in the primary experiment and alternative Laplace, standardized Student-$t_3$, or quantized-Gaussian distributions in sensitivity analyses. Hidden $T$ is a stylized utility and is never interpreted as AUROC.

The primary grid crossed three candidate topologies (aligned, competing, and misleading), three calibrated heterogeneity levels, three proxy strengths, and six score-noise ratios (0, 0.25, 0.5, 1, 2, and 4). Each of the 162 primary scenarios contained 10,000 replicate-indexed banks with $R = 5$, absolute $\delta = 0.01$, Gaussian score noise, and no forced duplicates. In aligned banks the candidates shared the same canonical prototype before mutation; competing banks introduced a second proxy-based prototype; and misleading banks gave 80% of candidates a systematically lower-utility majority prototype. Prespecified sensitivities varied $R$ over 3, 10, 20, 50, and 100 with a matched $R = 5$ reference; varied $\delta$ over 0, 0.005, 0.01, and 0.02; replaced Gaussian score noise; and introduced duplicate rates 0.1 and 0.3. The complete workload comprised 513 scenarios and 4,995,000 banks. No participant-level data, classifier, preprocessing operation, RFECV, Direct selector, or GA was simulated or fitted.

The four rules were highest score, legacy top-three Jaccard medoid, unrestricted full-bank Jaccard medoid, and regret-constrained medoid. Oracle regret was $\max_r T_r - T_{\hat{r}}$, empirical regret was $\max_r L_r - L_{\hat{r}}$, score optimism was $L_{\hat{r}} - T_{\hat{r}}$, and substantive failure was prespecified as oracle regret greater than 0.02. Full-bank mean Jaccard quantified representativeness. Primary results are equal-scenario means: the scenario grid is not a population prior. The regret-constrained medoid had zero configured empirical-regret violation in all 4,995,000 banks. Exact duplicate masks retained multiplicity as voting candidates, and source-run provenance remained post-selection metadata. The stable hash was evaluated only as the final deterministic ordering stage; it was not a scientific score.

Randomness used counter-based SplitMix64 streams keyed to the frozen master seed, scenario stream, and replicate identifier, making candidate generation independent of chunk boundaries, execution order, restart history, and worker count. All 1,296 completed chunks passed checksum and finite-value validation, and 18 prespecified deterministic regenerations matched exactly. The protocol's engineering list named chunk sizes of 1,000 or larger; the $R = 100$ sensitivity used chunks of 250 to limit peak memory. This execution-level deviation changed no scientific parameter, and all chunk-independence and integrity checks passed. Peak resident memory was 313.7 MiB. Figure and table source CSVs, their hashes, and the extraction provenance accompany the manuscript. Results characterize locking behavior under the specified utility. The claim boundary in Supplementary Methods S1.4 applies; the simulation does not establish predictive superiority or globally optimal feature selection.

##### S1.11 Computational complexity and implementation correspondence

Let $R$ be the total retained-candidate count, $E$ the regret-eligible count, $p$ the common canonical-universe size, $s$ the average selected-set size, and $w$ a packed machine-word width. Finding $L_{\text{best}}$, calculating all regrets, and forming $E_\delta$ each require $O(R)$ time; stored regrets and eligible indices require $O(R)$ and $O(E)$ memory, respectively, although a score-only pass can use constant working memory. Dense canonical masks and their stable mask identifiers require $O(Rp)$ time and $O(Rp)$ retained memory, or $O(p)$ mask scratch plus $O(R)$ digests if streamed. If no universe is supplied, deriving the package's sorted union adds $O(Rs + p \log p)$ time. A supplied common universe is preferred because it makes the scientific mask domain explicit.

A dense bit-vector Jaccard comparison costs $O(p)$ time, a sparse hash-set comparison costs expected $O(s)$ time (or $O(|F_r| + |F_s|)$ for sorted-index merging), and a packed-bit comparison costs $O(\lceil p/w \rceil)$ word operations with $O(E\lceil p/w \rceil)$ packed-mask storage. Thus all eligible comparisons cost $O(E^2 c_J)$, where $c_J$ is the representation-specific pair cost. Retaining the complete ordered $E \times E$ matrix requires $O(E^2)$ similarity memory. Streaming each unordered pair into $E$ running sums has the same asymptotic time but only $O(E)$ accumulator memory beyond the masks; a complete pairwise audit must then be streamed to storage or omitted.

After hashes exist, deterministic tie-breaking costs $O(E)$ with a best-key scan or $O(E \log E)$ by sorting. Canonically ordering the complete audit costs $O(R \log R)$. Audit serialization requires $O(Rp + E^2)$ time and output size when canonical masks and the full pairwise table are retained. The package's dense-mask, sparse-Jaccard, complete-audit implementation therefore has expected time $O(Rp + E^2 s + R \log R)$ and memory $O(Rp + E^2)$. With the manuscript's $R = 5$, there are at most 10 unordered off-diagonal comparisons or 25 ordered entries including the diagonal. This small use case does not establish scalability to large banks. Detailed operation-level bounds are supplied in `regret_locking_formalization/REGRET_LOCKING_COMPLEXITY.md`; implementation correspondence is summarized in Supplementary Table S25.

#### S2 Supplementary Figures

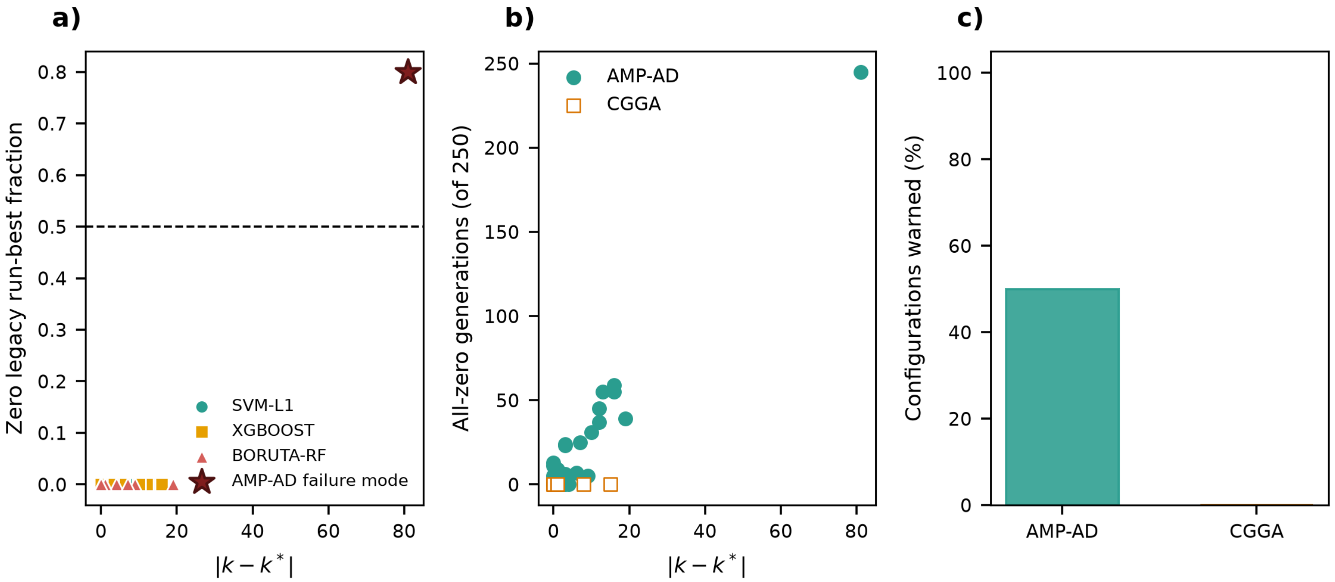


**Supplementary Figure S1.** Original-configuration zero-truncation diagnostics. a) Absolute target-size deviation versus the fraction of zero run-best fitness values. Circles, squares, and triangles denote SVM-L1, XGBoost, and Boruta-RF; the star marks the AMP-AD Rush/SVM-L1/Small failure mode, and the dashed line marks a zero-fraction value of 0.5. b) Absolute target-size deviation versus all-zero generations. Filled teal circles denote AMP-AD and open orange squares denote CGGA. c) Percentage of configurations with a flattening warning in each dataset.

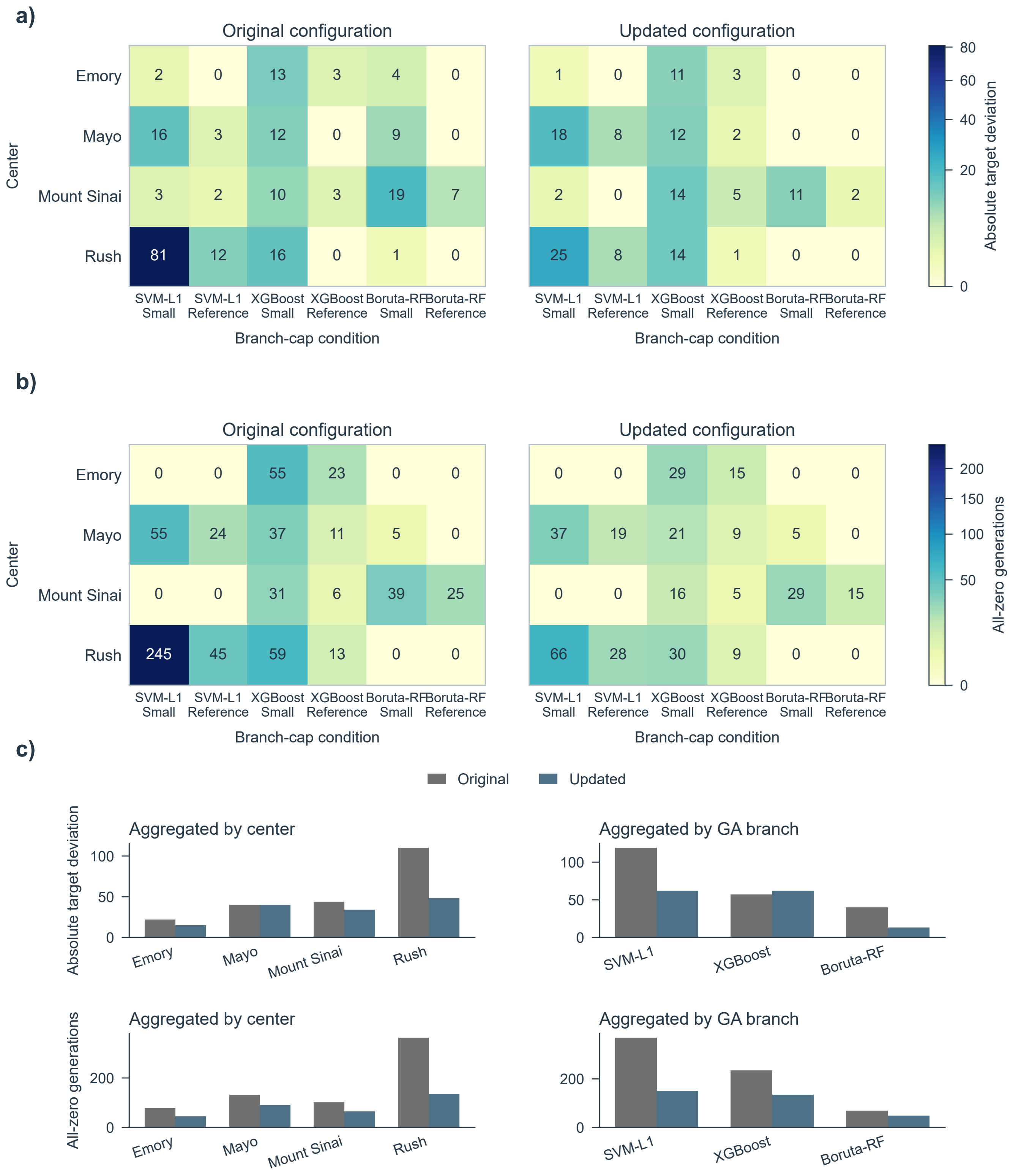


**Supplementary Figure S2.** Complete development-only original-versus-corrected-objective AMP-AD comparison across 24 combinations of center, branch, and cap and 120 full GA runs. a) Absolute target deviation. b) All-zero-generation diagnostic. c) Aggregates by center and branch. Overall values were 216 versus 137 and 673 versus 333, respectively; detailed robust summaries are in Supplementary Table S2. Held-out outcomes were not accessed.

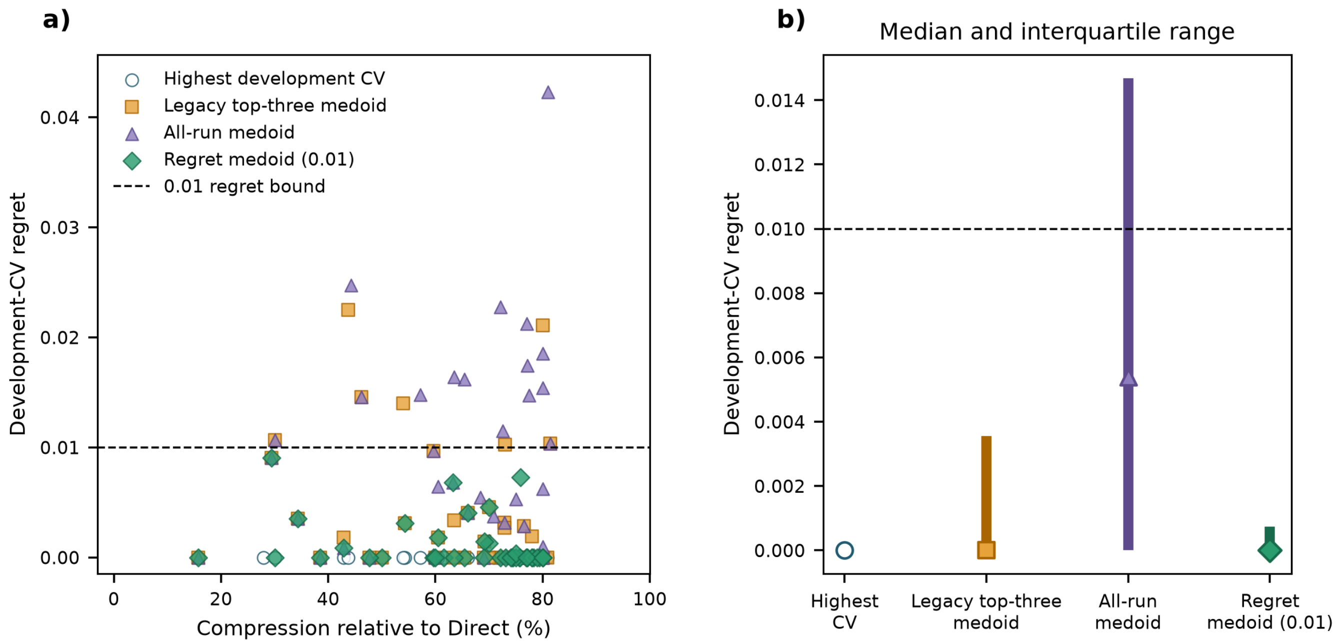


**Supplementary Figure S3.** Development-only compression and regret. a) Compression relative to each Direct candidate set versus development-CV regret for highest-locking-score selection (open circles), the legacy top-three medoid (squares), the unrestricted full-bank medoid (triangles), and the regret-constrained medoid at absolute tolerance 0.01 (diamonds). The dashed line marks the 0.01 empirical-regret bound. b) Median development-CV regret (markers) and interquartile range (vertical bars) by locking rule. Held-out outcomes were not used to define any displayed rule.

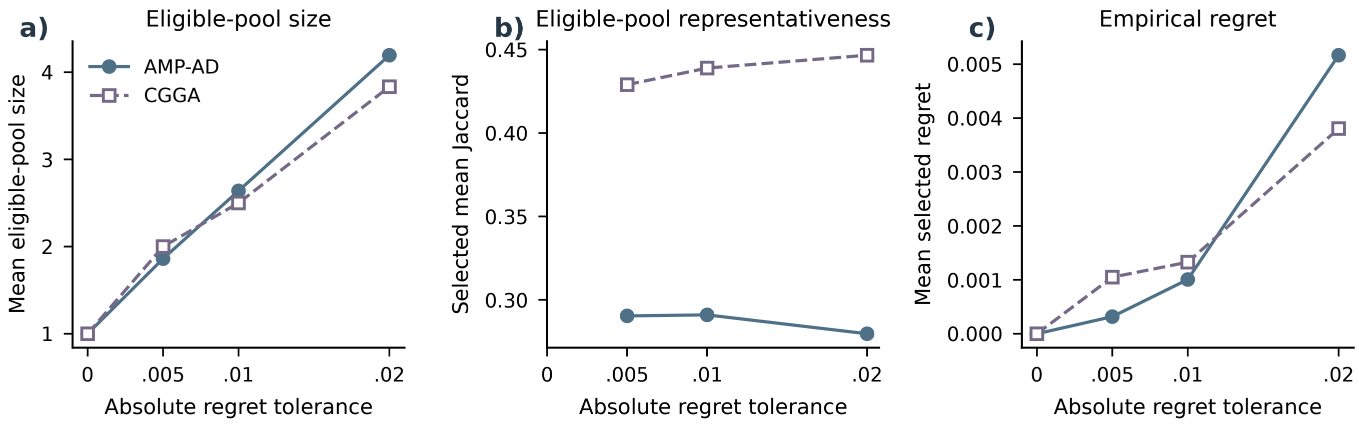


**Supplementary Figure S4.** Development-only strict locking-rule sensitivity. a) Mean eligible-pool size. b) Selected-candidate mean eligible-pool Jaccard for nonsingleton pools; singleton values are undefined and shown as missing. c) Mean selected empirical development-CV regret across absolute tolerances 0, 0.005, 0.01, and 0.02. No tolerance-expanding fallback was used.

*Supplementary Figure S5. Complete candidate-level audit of fully nested TCGA locking (continued on the next page).*

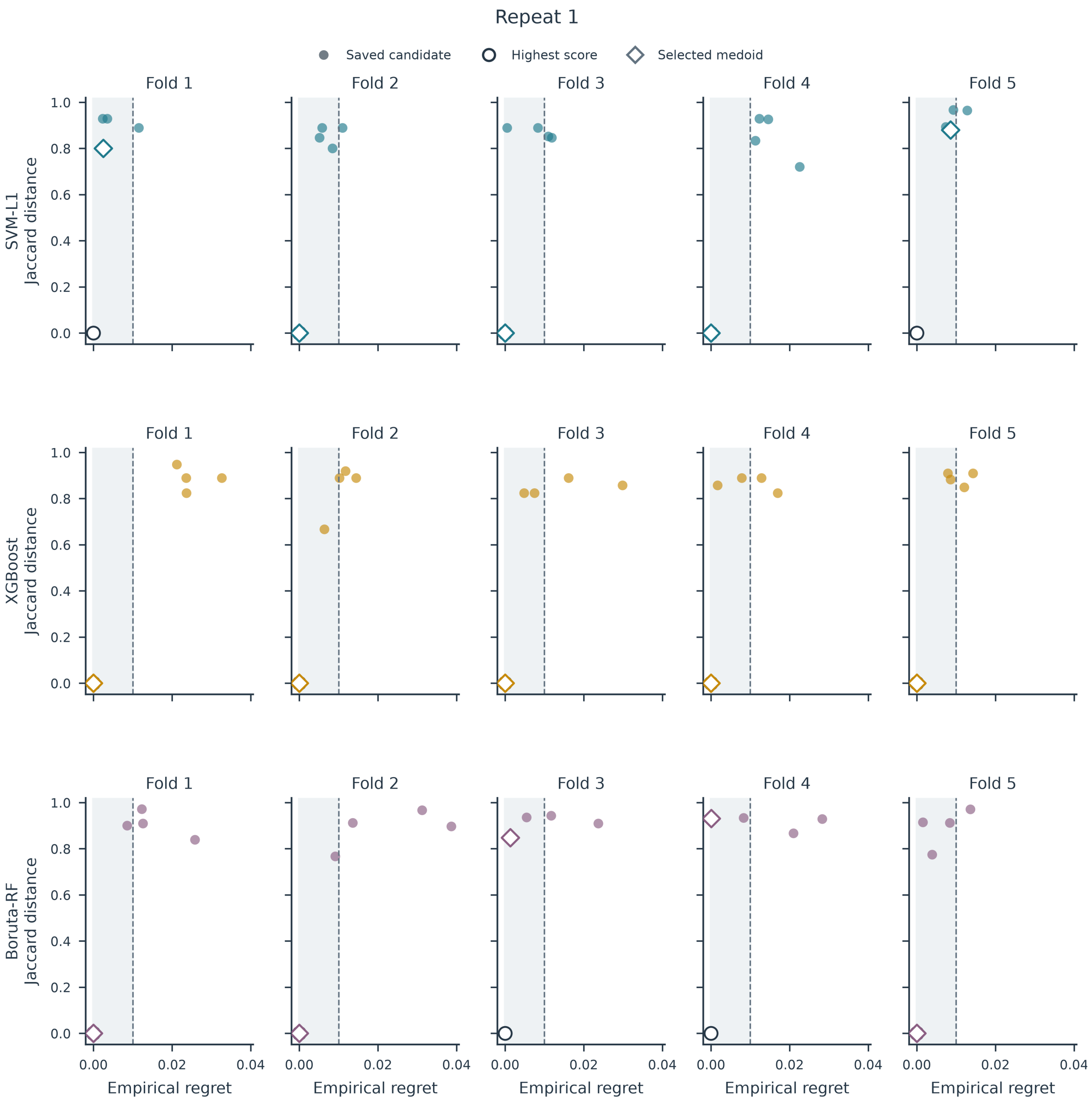

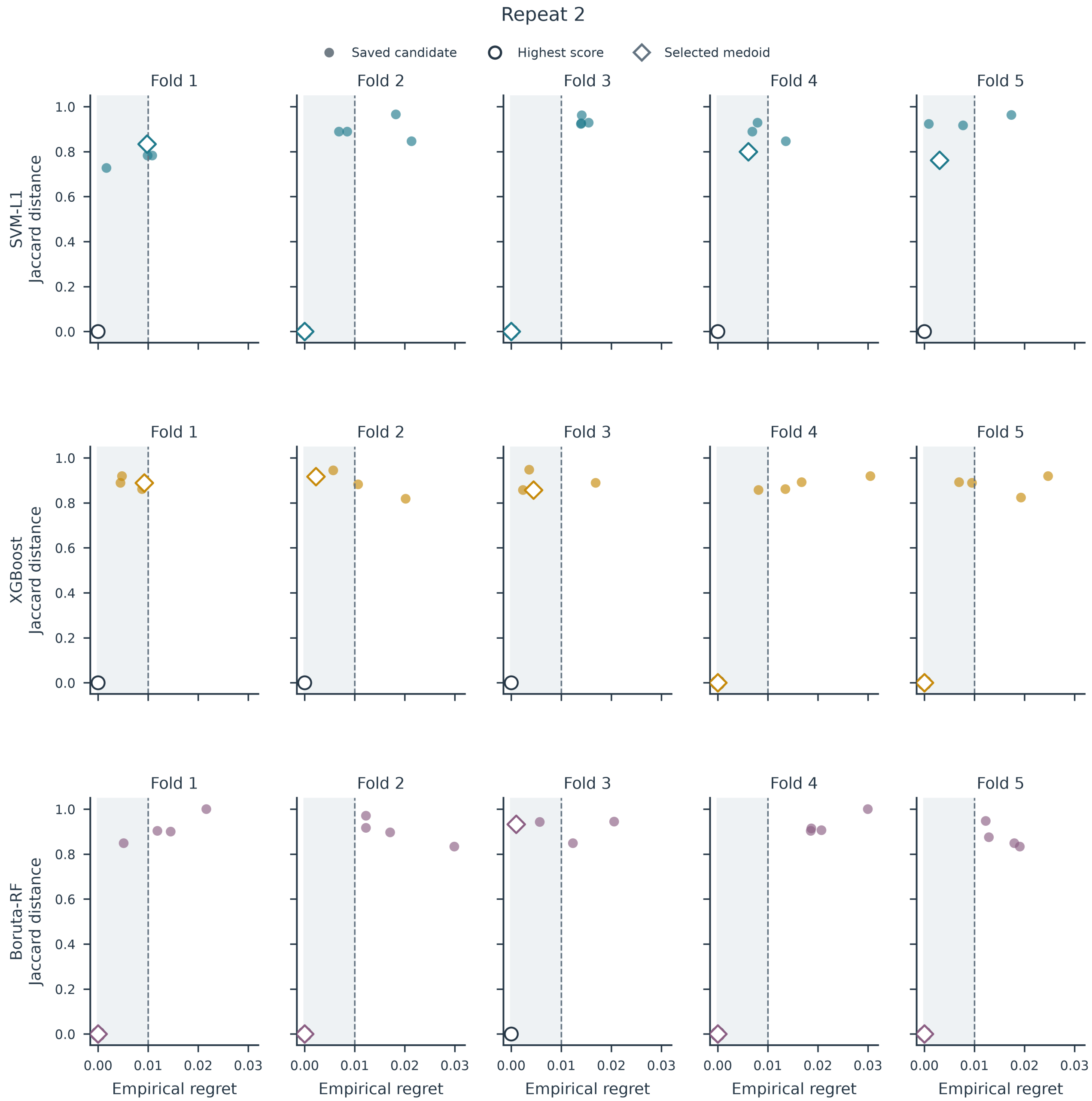


**Supplementary Figure S5.** Complete candidate-level audit of fully nested TCGA locking. The first and second pages show outer repeats 1 and 2, respectively. Each panel represents one branch and outer fold. Points are the five saved GA candidates, positioned by empirical development-CV regret and Jaccard distance from the highest-locking-score mask. Shading denotes the absolute regret-eligible region $R_r \leq 0.01$, and dashed lines mark $\delta = 0.01$. Open circles identify highest-locking-score candidates and diamonds identify selected regret-constrained medoids; markers overlap when the same candidate satisfies both roles. Selection and locking used outer-training data only.

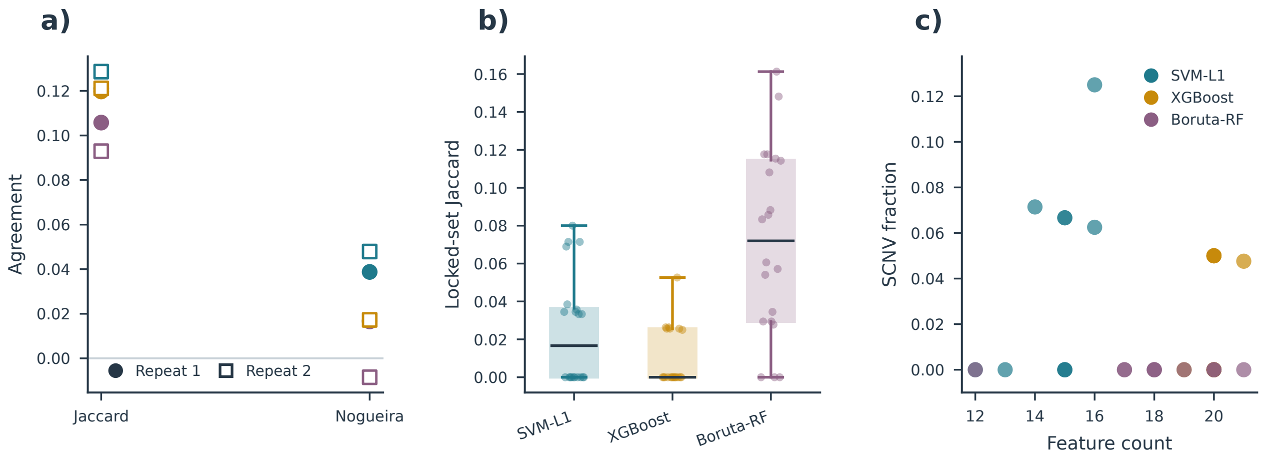


**Supplementary Figure S6.** Candidate agreement, partition sensitivity, and locked-signature composition in TCGA GBM/LGG. a) Mean pairwise Jaccard and chance-corrected Nogueira stability among five saved candidates. Each point is the unweighted mean of five fold-level bank statistics for one repeat–branch combination and is conditional on each bank's fixed post-Direct universe. Filled circles and open squares denote repeats 1 and 2. The coefficients are shown side by side and are not connected as an effect estimate; a small negative finite-sample Nogueira estimate is valid. b) Jaccard similarity for the 20 dependent outer-fold locked-signature pairs per branch, summarized descriptively. c) Selected feature count and SCNV fraction for each of 30 locked signatures. Eleven of 526 selected-feature occurrences were SCNV features; composition largely reflects upstream Direct screening and is not evidence of biological modality preference. These summaries characterize conditional algorithmic agreement and participant-partition sensitivity, not biomarker stability.

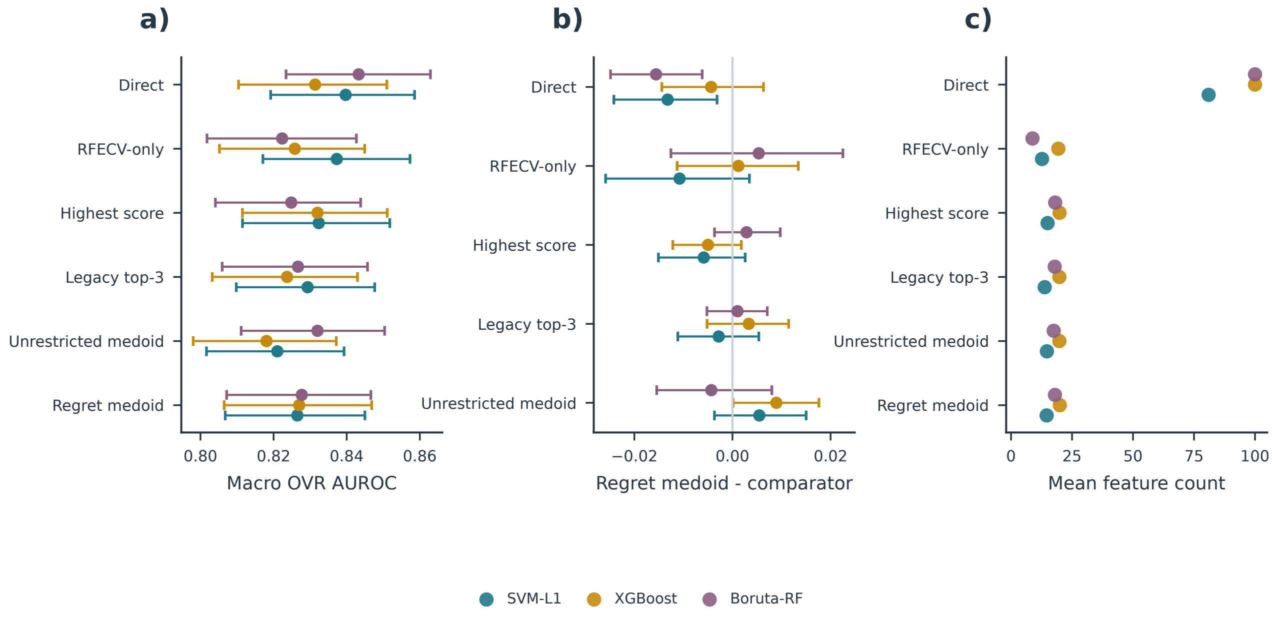


**Supplementary Figure S7.** Repeated internal out-of-fold comparator audit in TCGA GBM/LGG. a) Mean repeated-OOF macro one-vs-rest AUROC for Direct, RFECV-only, highest-locking-score selection, the legacy top-three medoid, the unrestricted full-bank medoid, and the regret-constrained medoid. b) Paired regret-constrained-medoid-minus-comparator AUROC differences. Points in a) and b) have 95% participant-clustered, class-stratified bootstrap intervals from 2,000 resamples retaining both repeat-specific predictions for each sampled participant. c) Mean feature count across ten outer folds. The protocol capped each post-Direct universe at 100 features. Colors identify branches. All three regret-constrained-medoid-versus-RFECV-only intervals included zero; equivalence was not tested.

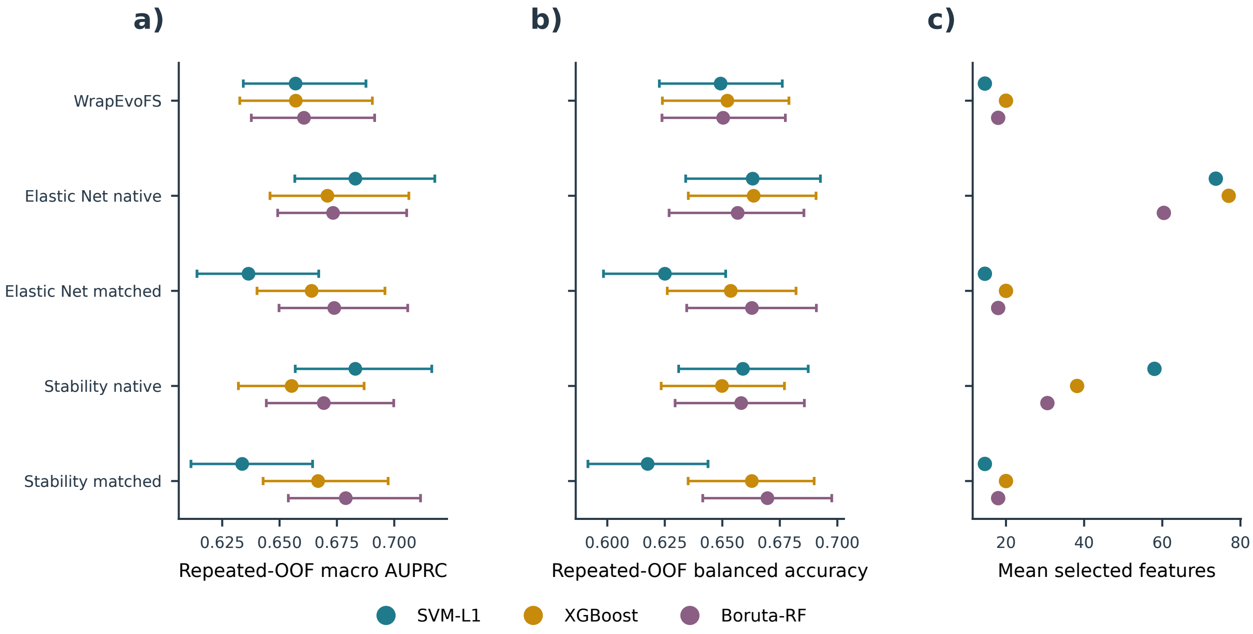


**Supplementary Figure S8.** Extended fully nested TCGA selector-layer sensitivity. a) Repeated-OOF macro AUPRC for regret-constrained-medoid WrapEvoFS, native and cardinality-matched Elastic Net, and native and cardinality-matched subsampling stability selection. b) Balanced accuracy for the same selectors. Points in a) and b) have 95% participant-clustered, class-stratified bootstrap intervals from 2,000 resamples retaining both repeat-specific predictions for each sampled participant. c) Mean selected-feature counts across ten outer folds. Colors identify Direct branches. Native and matched signatures answer different questions; the matched sets are prespecified cardinality sensitivities, not native nonzero supports. The analysis is conditional on verified post-Direct universes.

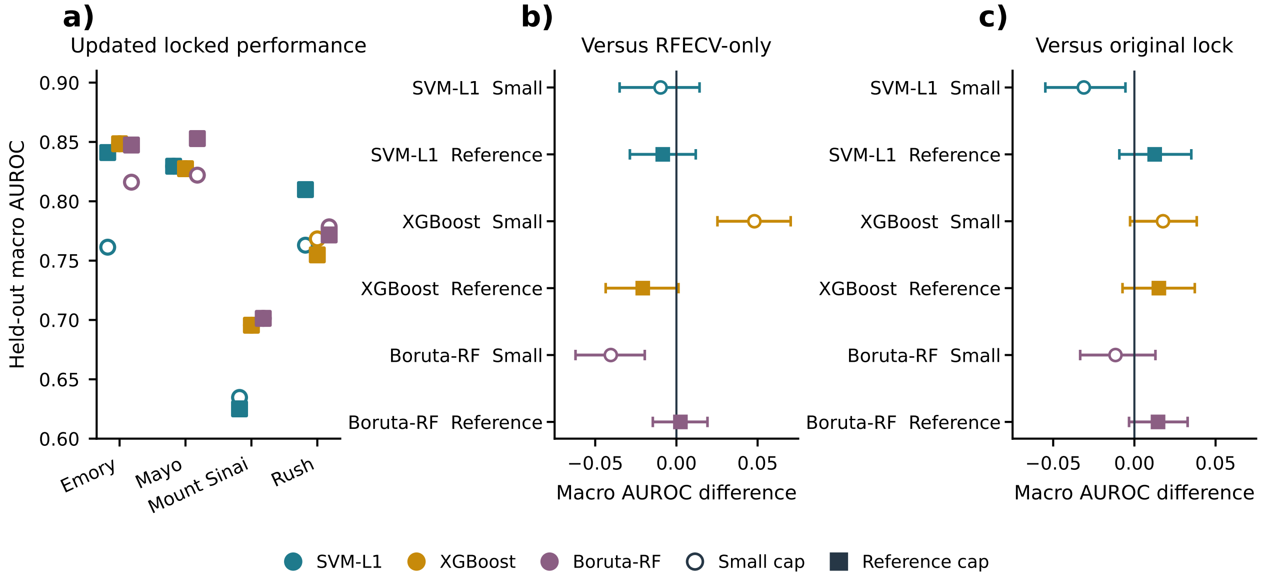


**Supplementary Figure S9.** AMP-AD one-time post-freeze held-out evaluation of the corrected objective with regret-constrained medoid locking. a) Macro one-vs-rest AUROC for 24 frozen signatures across combinations of center, branch, and cap; colors identify branches, open circles denote the Small cap, and filled squares denote the Reference cap. b) Pooled regret-constrained-medoid-minus-RFECV-only and c) regret-constrained-medoid-minus-legacy-top-three-medoid macro-AUROC differences with 95% intervals from 2,000 common resamples stratified by held-out center and true class (seed 42). All masks, parameters, and the protocol hash were frozen before held-out outcomes were loaded; evaluation triggered no reselection or parameter changes. Four of six intervals in b) and five of six in c) include zero.

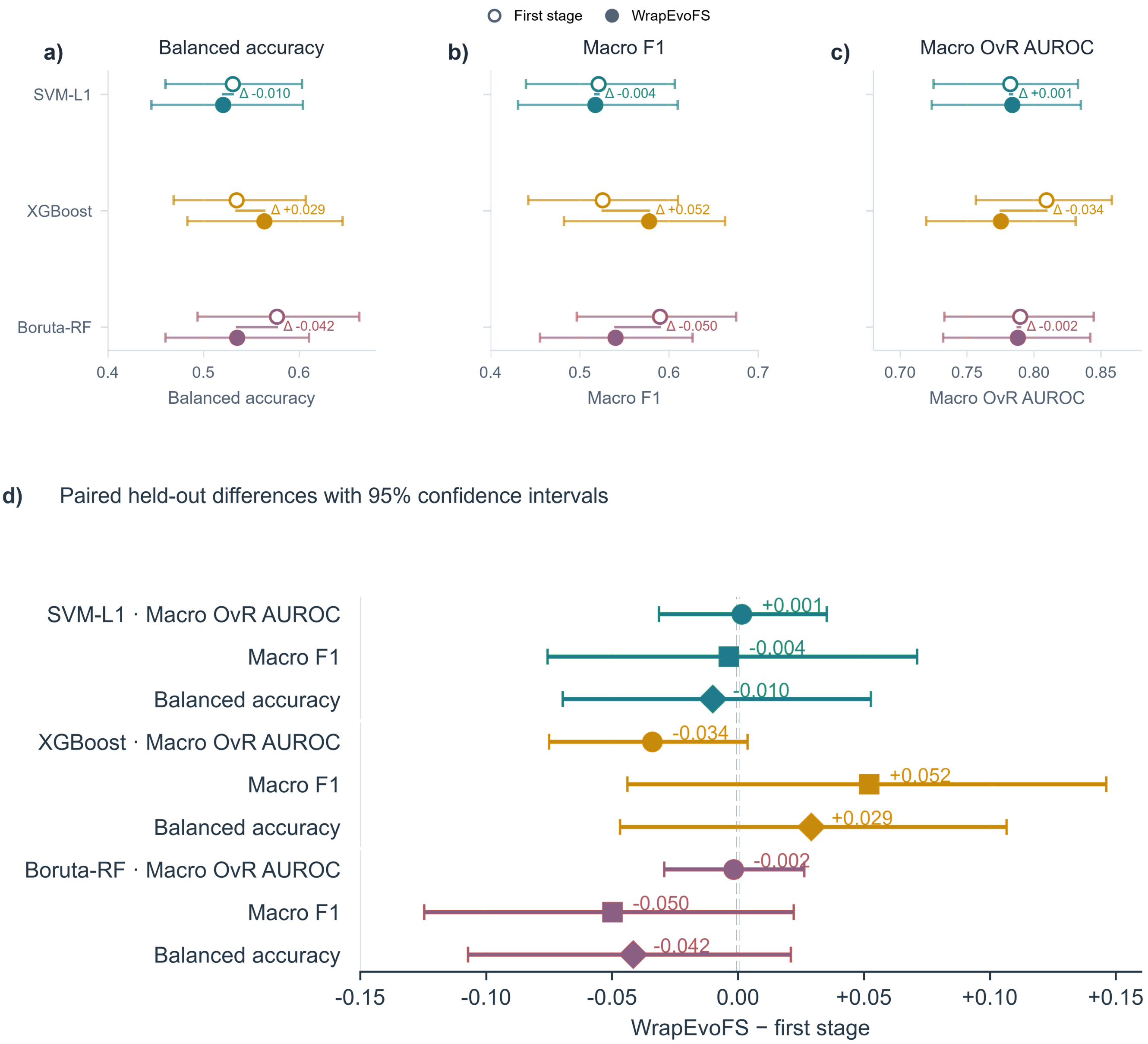


**Supplementary Figure S10.** ADNI held-out performance and paired differences. a) Balanced accuracy, b) macro F1, and c) macro one-vs-rest AUROC for the first-stage Direct signatures (open symbols) and archived legacy-top-three-medoid signatures (filled symbols), shown by branch with 95% bootstrap confidence intervals. d) Paired legacy-top-three-medoid-minus-first-stage differences from 5,000 class-stratified paired bootstrap resamples. All nine paired 95% confidence intervals include zero; equivalence and non-inferiority were not tested.

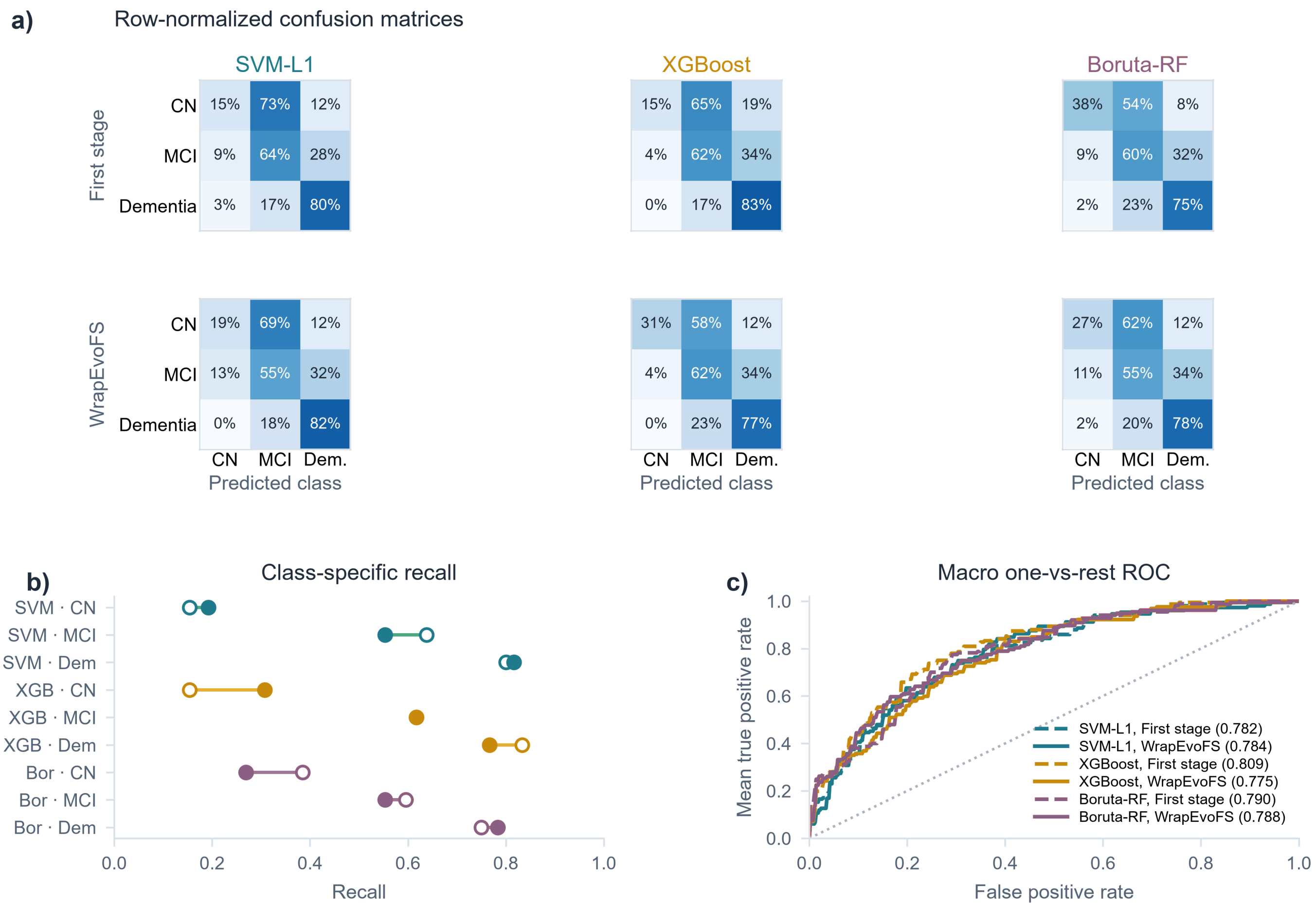


**Supplementary Figure S11.** ADNI multiclass held-out diagnostics. a) Row-normalized confusion matrices for the first-stage Direct and WrapEvoFS locked signatures in the SVM-L1, XGBoost, and Boruta-RF branches. b) Class-specific recall for CN, MCI, and dementia, with open and filled symbols denoting first-stage and WrapEvoFS signatures. c) Macro one-vs-rest ROC curves and AUROC values for the same six models.

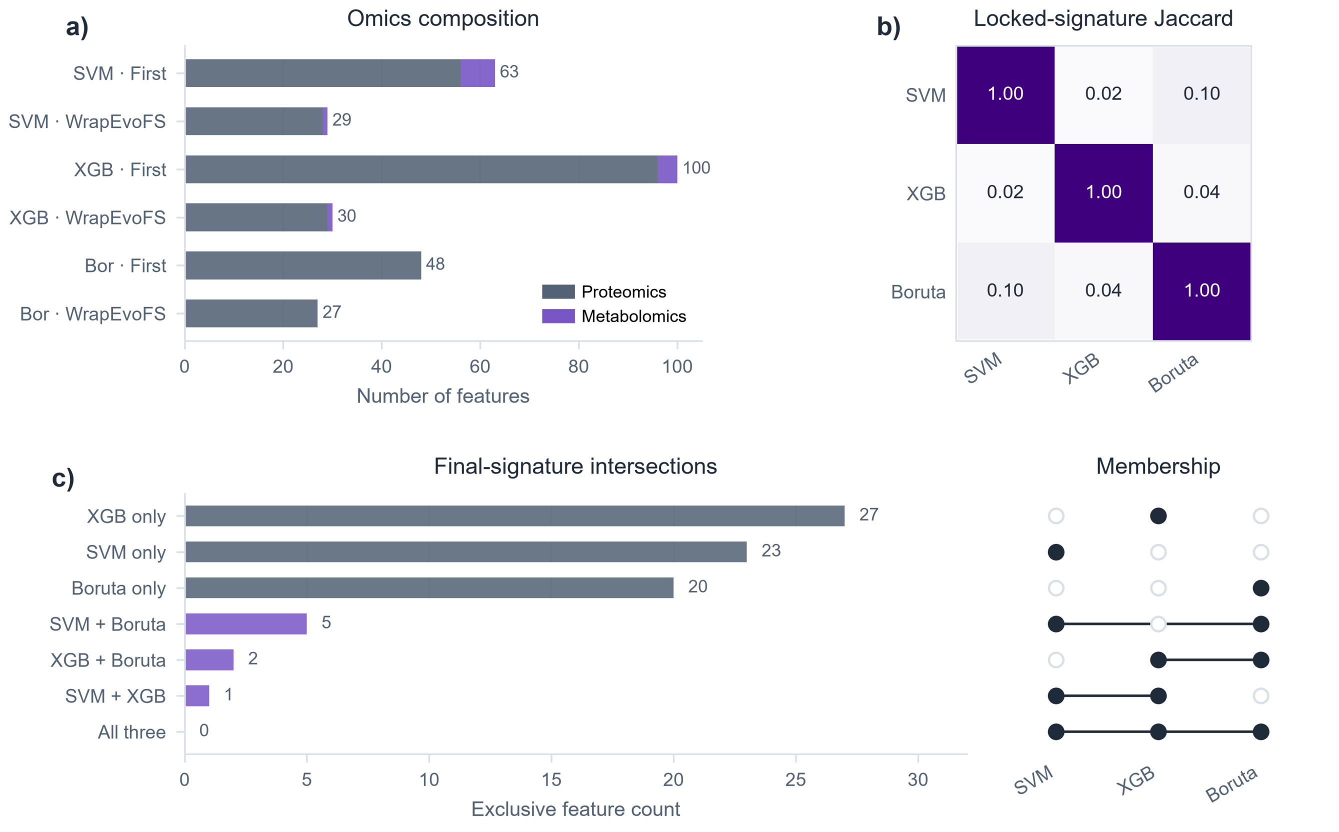


**Supplementary Figure S12.** ADNI signature composition and cross-branch overlap. a) Proteomic and metabolomic feature counts for first-stage and WrapEvoFS signatures in each branch. b) Pairwise Jaccard similarity among the three locked signatures. c) Exclusive and shared locked-signature feature counts shown with an intersection-membership matrix. The displayed sets are descriptive and are not stable biomarker claims.

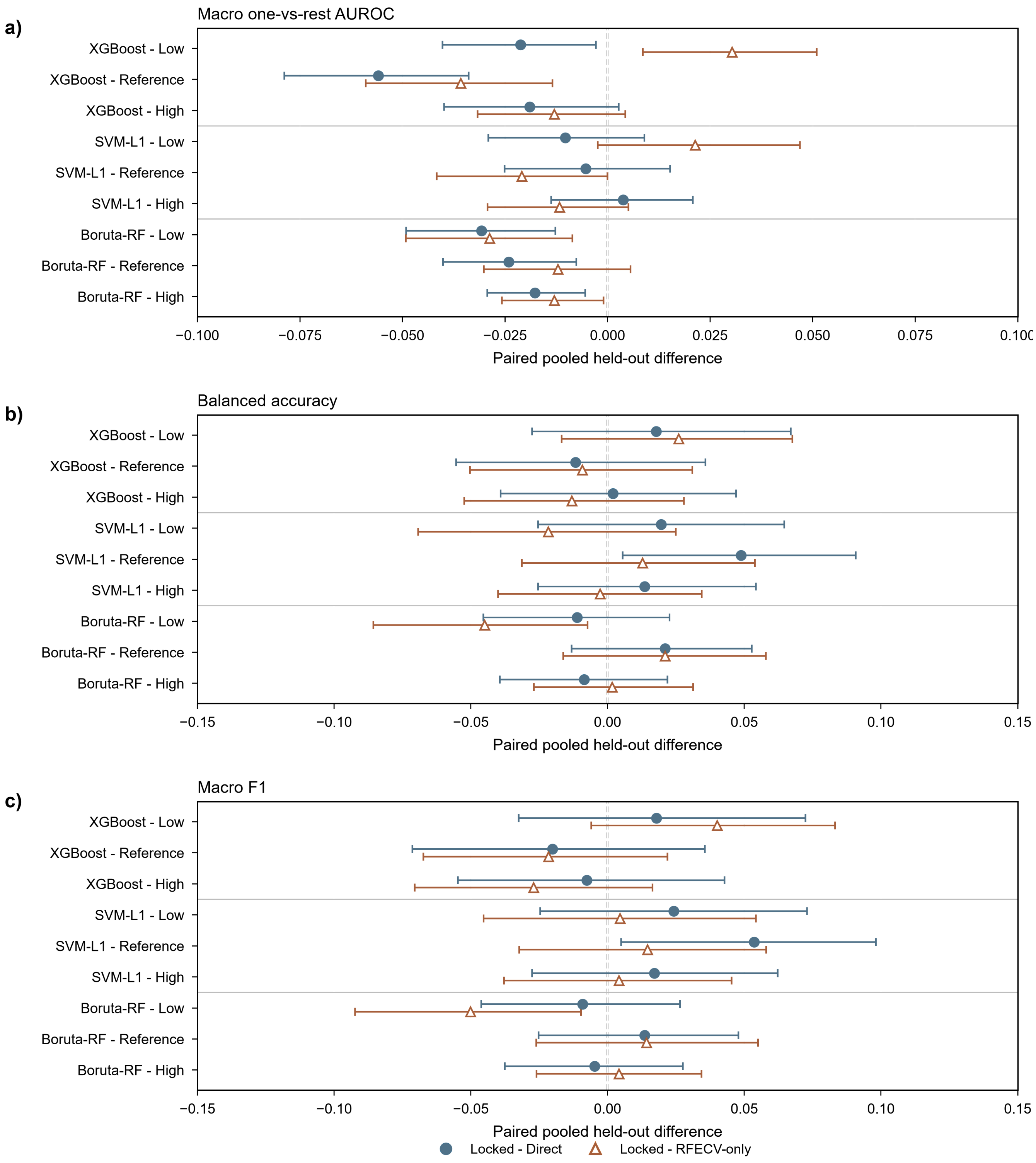


**Supplementary Figure S13.** AMP-AD paired pooled held-out differences across RFECV cap settings. a) Macro one-vs-rest AUROC, b) balanced accuracy, and c) macro F1. Filled circles show locked-minus-Direct differences and open triangles show locked-minus-RFECV-only differences; horizontal intervals are 95% paired confidence intervals. Forty of 54 exported intervals include zero; directions vary by branch, cap, metric, and comparator, and resampling conditions on four fitted center-specific models.

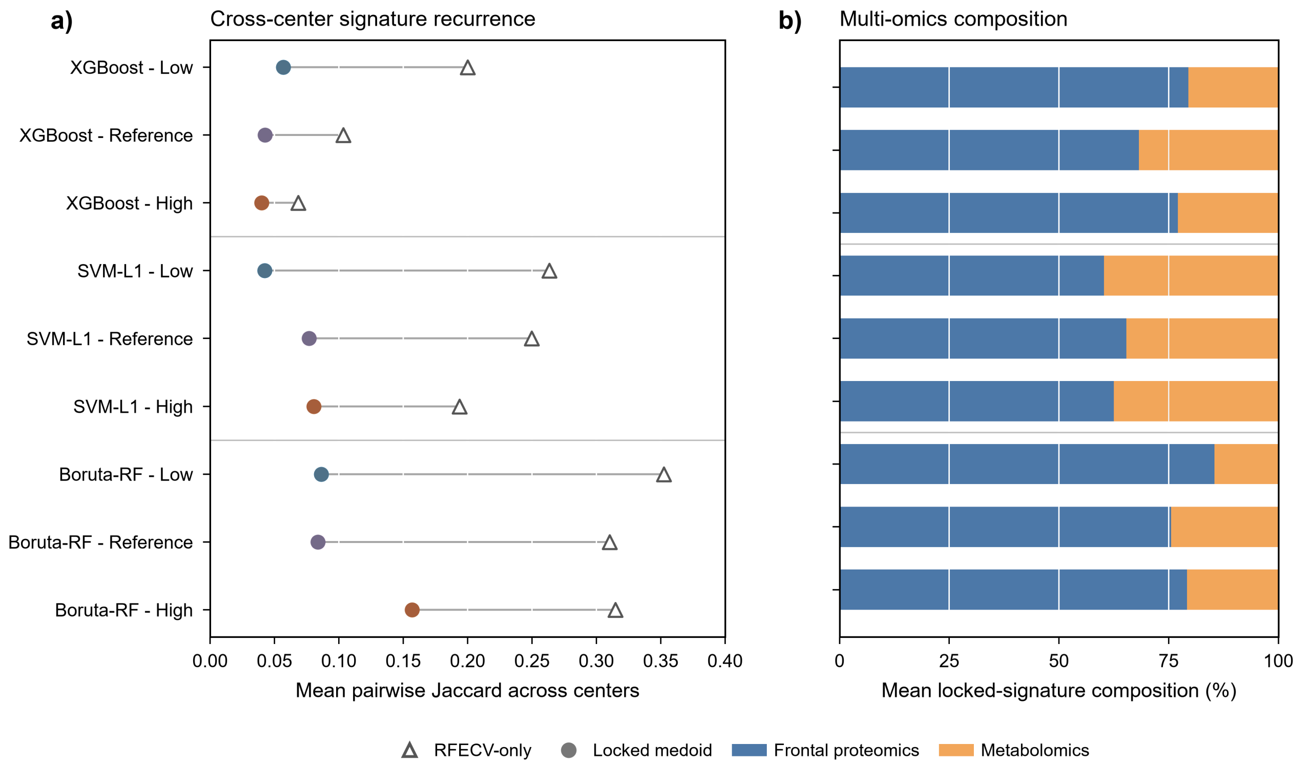


**Supplementary Figure S14.** AMP-AD cross-center recurrence and modality composition. a) Mean pairwise Jaccard similarity across centers for RFECV-only and archived legacy-top-three-medoid signatures in each branch and cap condition. b) Mean locked-signature composition split between frontal proteomics and metabolomics for the same conditions. Cross-center overlap quantifies recurrence across center-specific training sets, not biological replication.

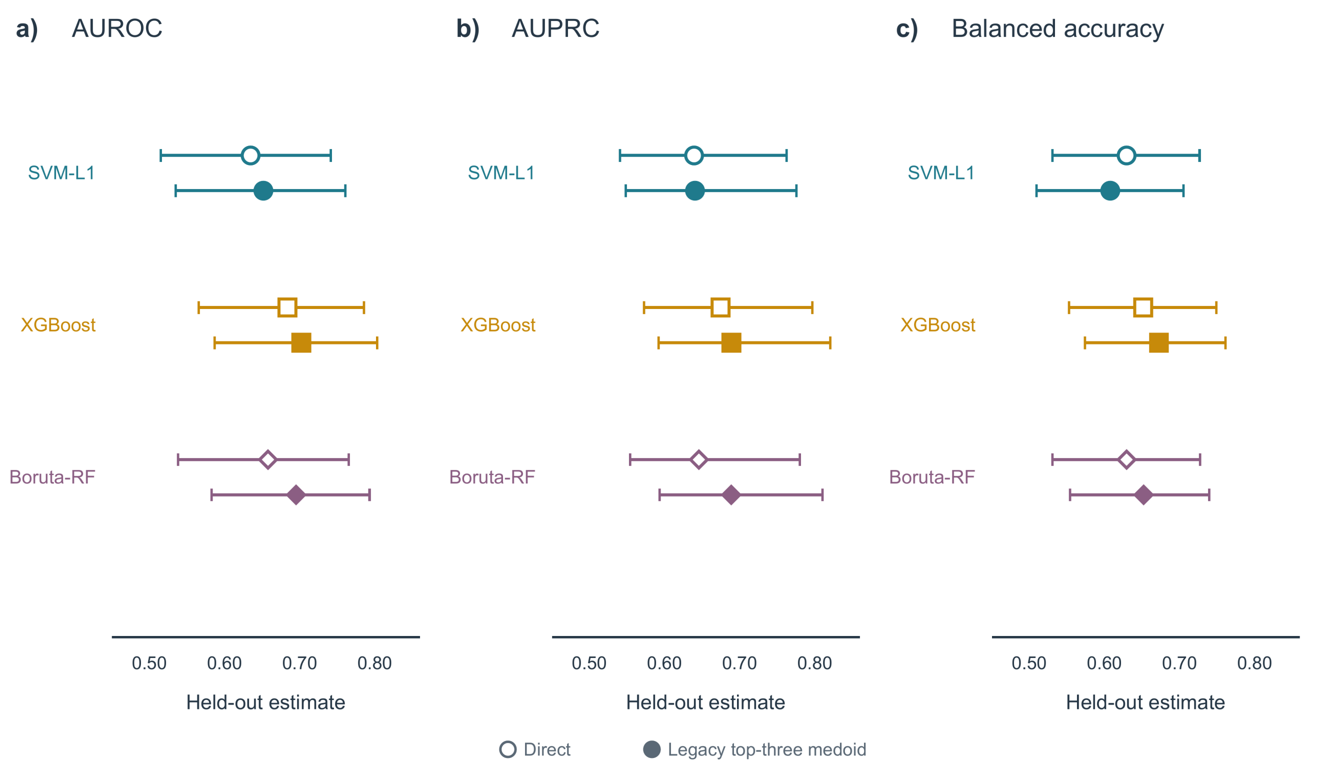


**Supplementary Figure S15.** Complete CGGA held-out comparison. a) AUROC, b) AUPRC, and c) balanced accuracy for the Direct signatures (open symbols) and legacy-top-three-medoid signatures (filled symbols) in the SVM-L1, XGBoost, and Boruta-RF branches; horizontal intervals are the corresponding held-out uncertainty intervals.

*Supplementary Figure S16. CGGA run-level stochastic agreement (continued on the next page).*

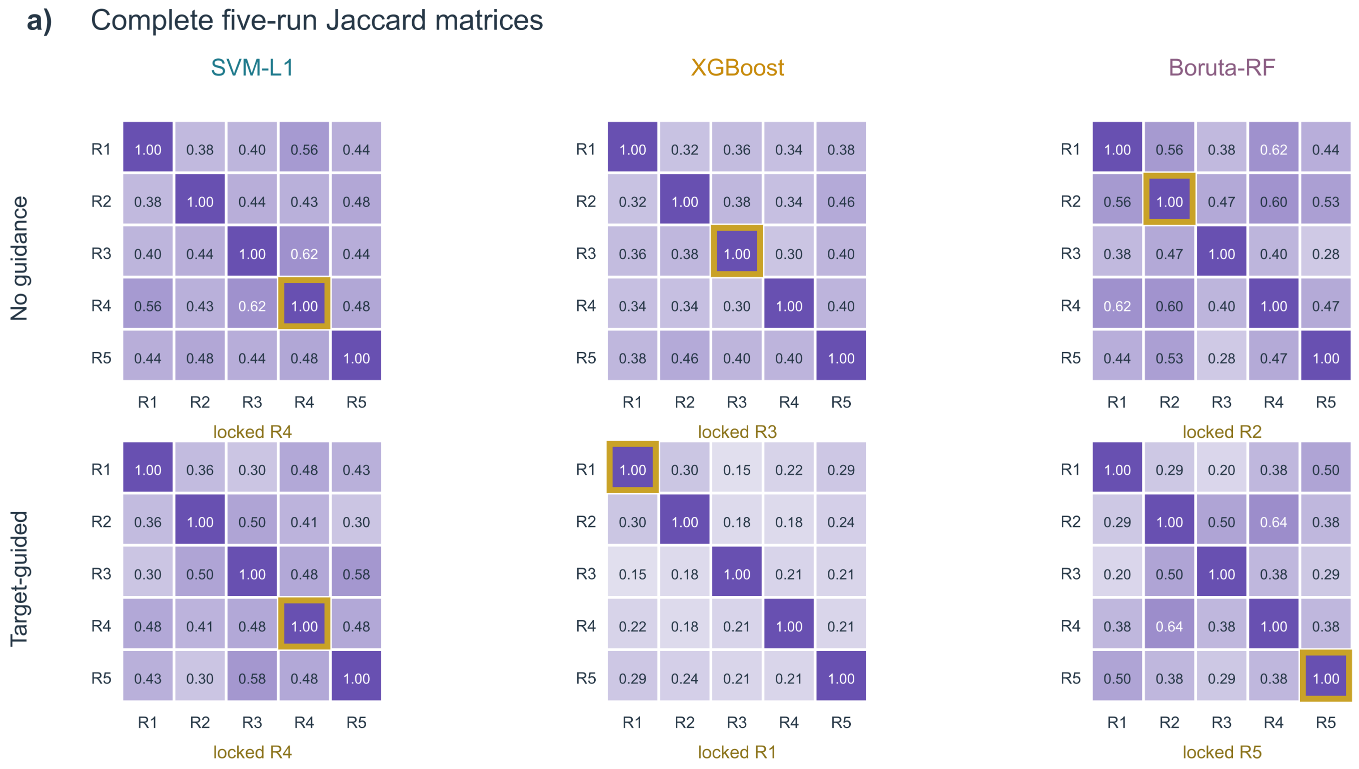

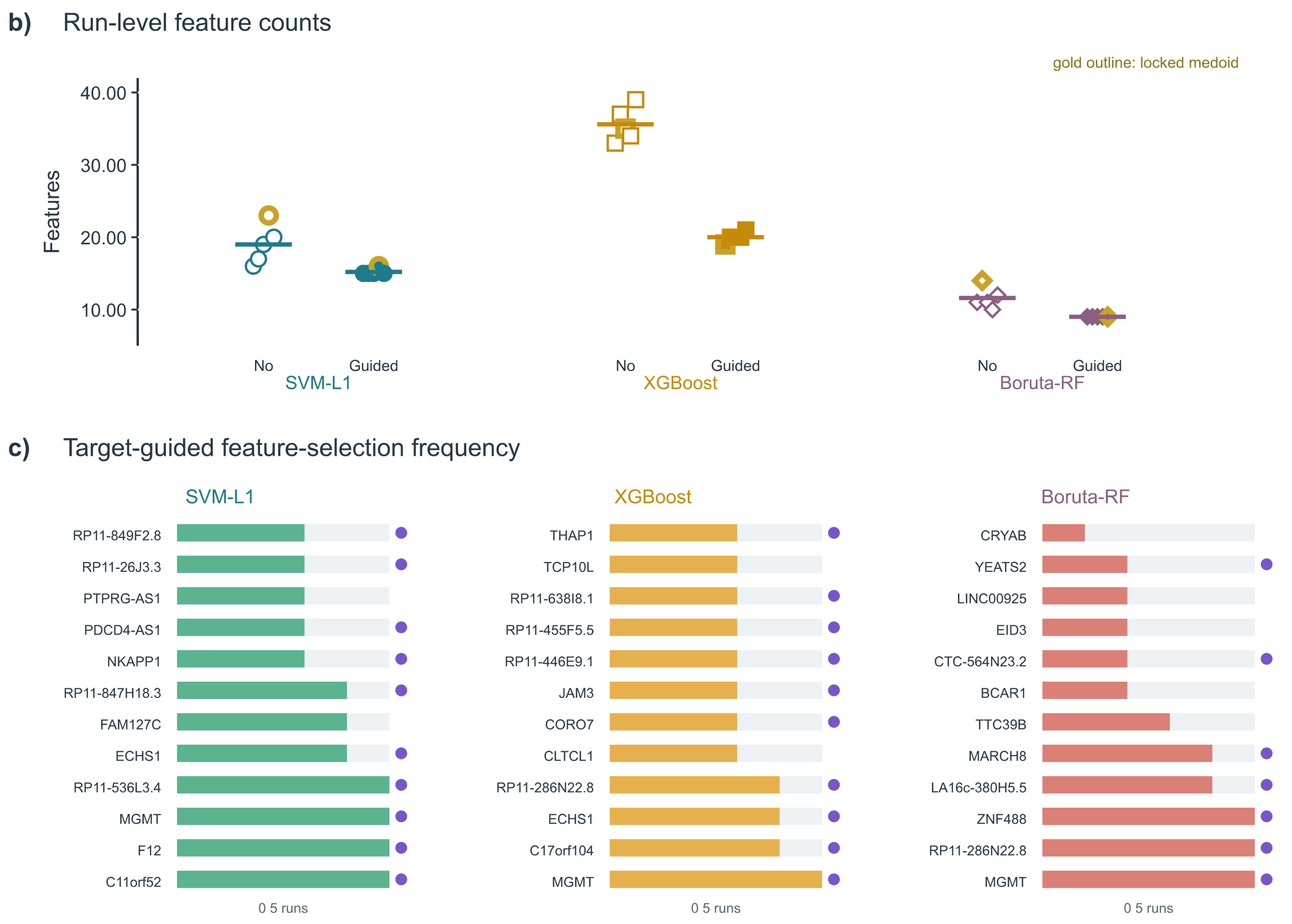


**Supplementary Figure S16.** CGGA run-level stochastic agreement. a) Five-run Jaccard matrices without and with target-size guidance; gold outlines mark the selected legacy top-three medoid run. b) Run-level feature counts without and with guidance. c) Target-guided selection frequencies; filled dots mark the legacy top-three medoid signature.

*Supplementary Figure S17. CGGA original-configuration mechanism and seed-agreement analysis (continued on the next page).*

**a)** Ablation of genetic refinement, size guidance, and run locking

| | RFECV only | Single GA run | Highest dev-CV | Legacy top-three | No target-size guidance |
|---|---|---|---|---|---|
| SVM-L1 | 99.7%<br>n=15 (−57%) | 104.7%<br>n=15 (−57%) | 104.7%<br>n=15 (−57%) | 102.7%<br>n=16 (−54%) | 98.1%<br>n=23 (−34%) |
| XGBoost | 98.7%<br>n=20 (−71%) | 102.7%<br>n=19 (−73%) | 104.8%<br>n=20 (−71%) | 102.7%<br>n=19 (−73%) | 94.4%<br>n=35 (−50%) |
| Boruta-RF | 103.7%<br>n=9 (−65%) | 104.2%<br>n=9 (−65%) | 105.7%<br>n=9 (−65%) | 105.7%<br>n=9 (−65%) | 101.2%<br>n=14 (−46%) |

Cells: AUROC/Direct ratio; n; feature reduction — Direct=100%

Single GA run = prespecified run 1

**b)** Target-size guidance and run diversity

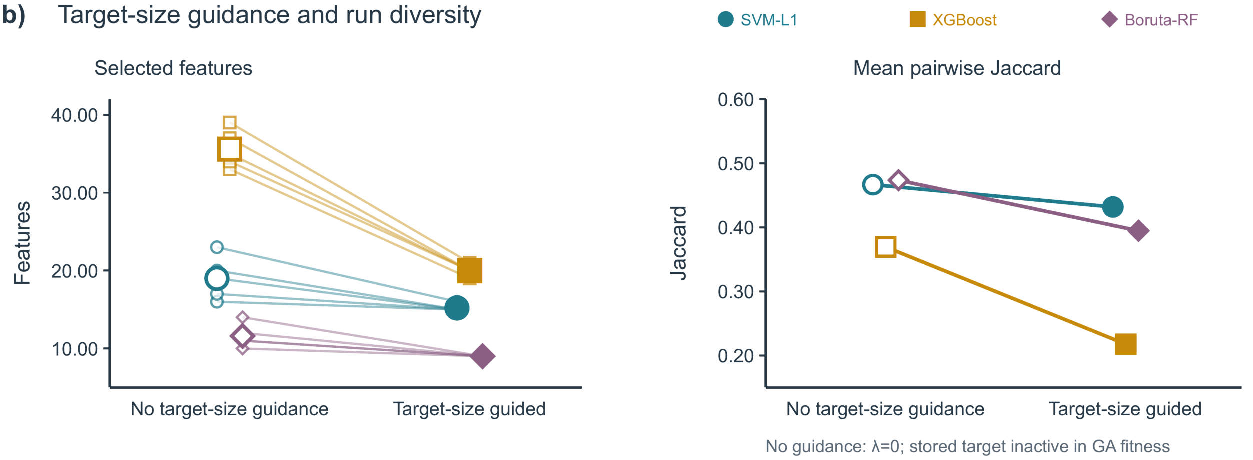

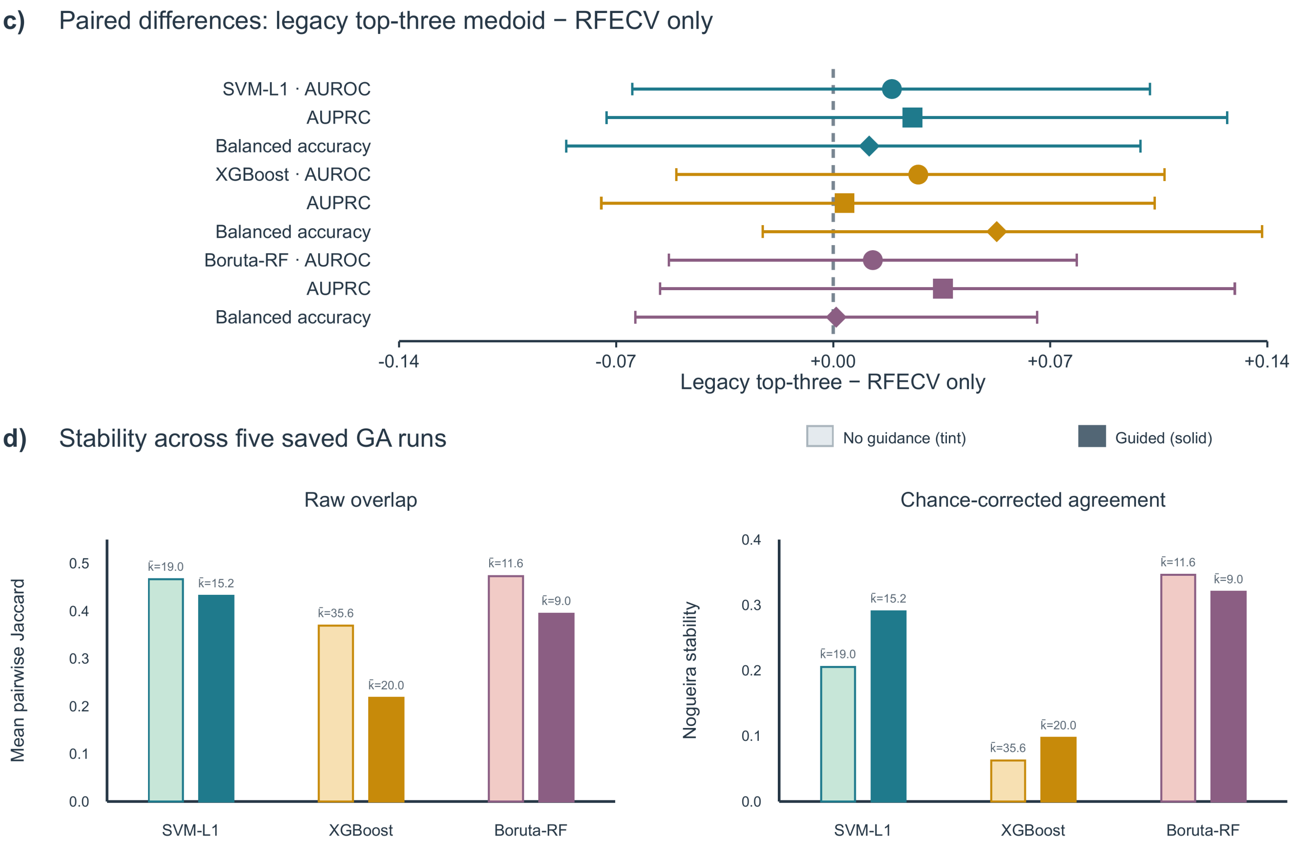


**Supplementary Figure S17.** CGGA original-configuration mechanism and seed-agreement ablation. a) Held-out AUROC relative to Direct for RFECV-only and selected-run variants. b) Five-run feature counts and mean pairwise Jaccard with and without target-size guidance. c) Paired legacy-top-three-medoid-minus-RFECV-only differences. d) Raw Jaccard and chance-corrected Nogueira agreement.

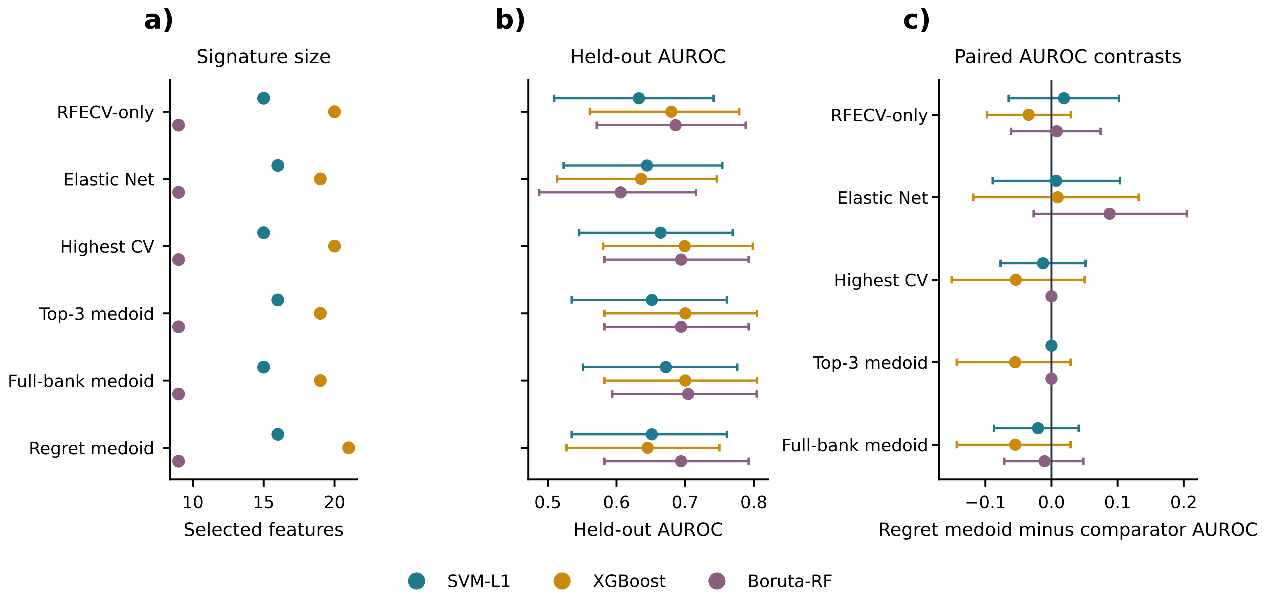


**Supplementary Figure S18.** CGGA coherent same-split, same-estimator benchmark. a) Feature counts and b) held-out AUROC with 95% class-stratified bootstrap intervals for six development-derived signatures in each branch. c) Paired regret-constrained-medoid-minus-comparator AUROC differences from 2,000 common resamples (seed 42); all 15 intervals include zero. All signatures were evaluated with the same fixed 500-tree random forest on the same 92-sample held-out partition. Held-out outcomes did not determine any feature set, locking rule, or comparator.

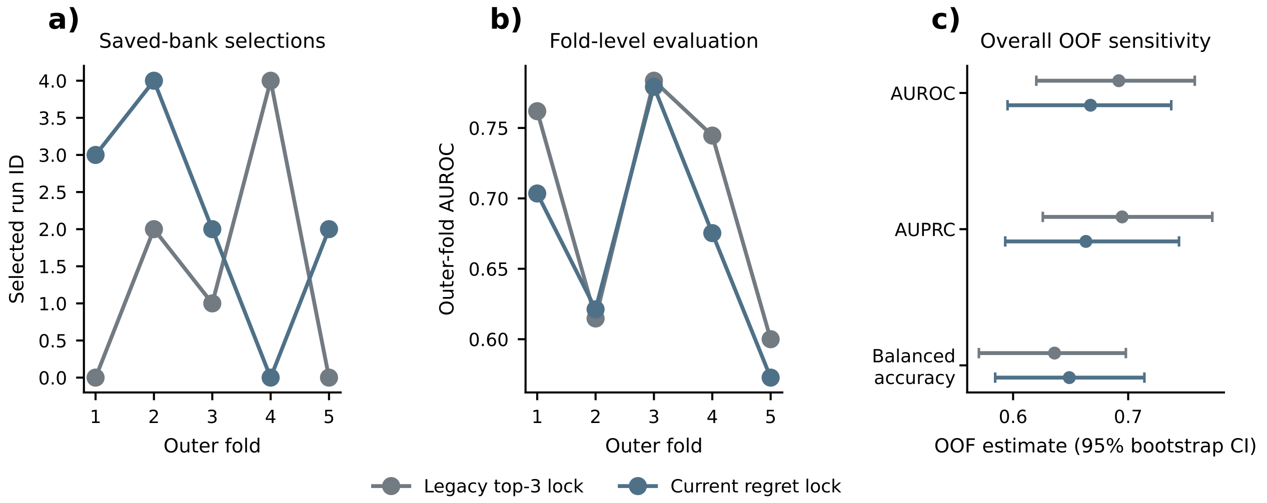


**Supplementary Figure S19.** CGGA post hoc saved-bank nested sensitivity. a) Selected run IDs under the legacy top-three medoid and regret-constrained medoid across five outer folds. b) Corresponding outer-fold AUROC. c) Overall OOF AUROC, AUPRC, and balanced accuracy with 95% class-stratified bootstrap intervals. The regret-constrained medoid changed all five selected runs; OOF AUROC was 0.692 under the legacy top-three medoid and 0.667 under the regret-constrained medoid. The source banks used a reduced historical search budget (20 individuals, 12 generations, five runs), no GA was rerun, and the original 92-sample held-out partition was not accessed.

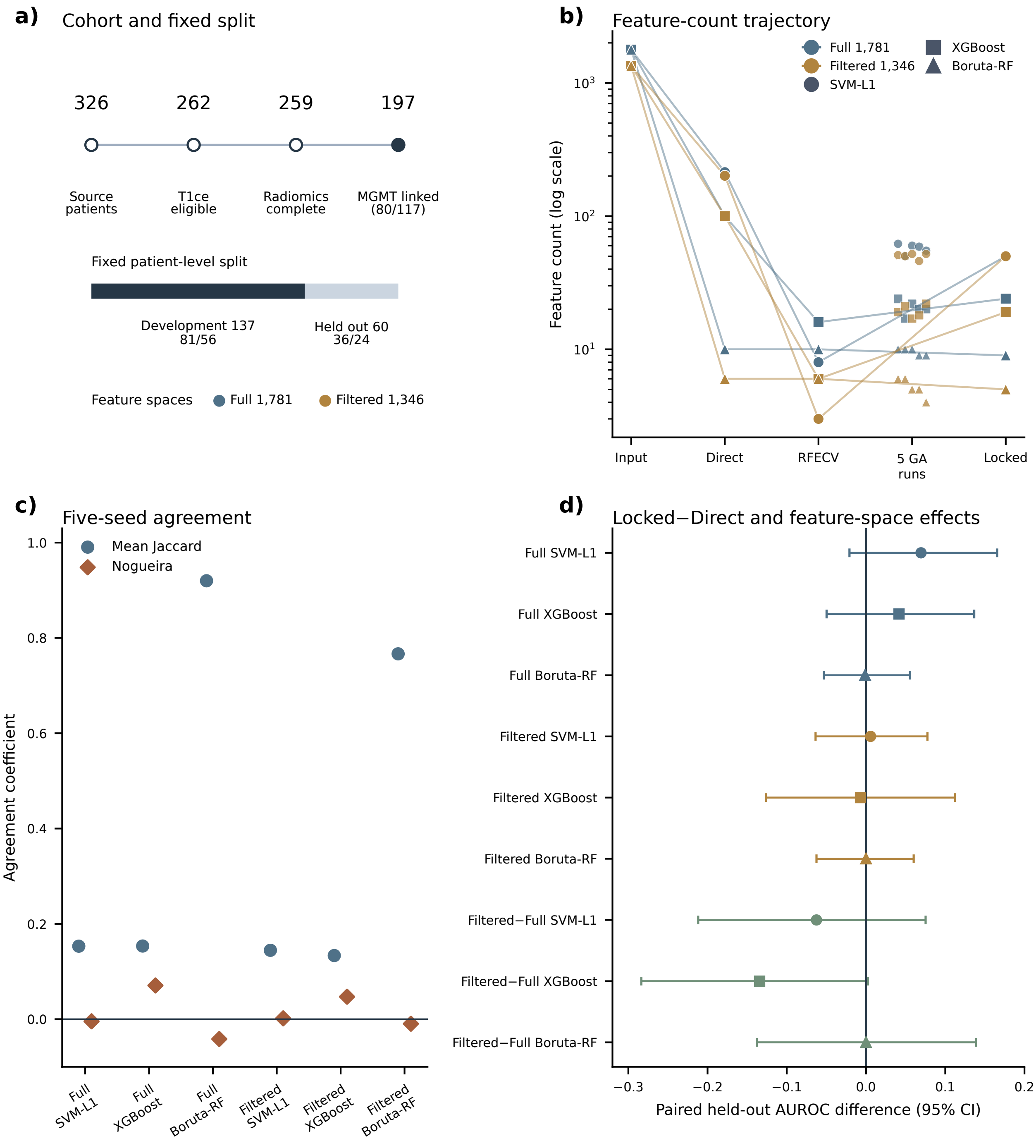


**Supplementary Figure S20.** VGH brain-tumour radiomics internal demonstration (UBC REB H20-02354) using regret-constrained locking. a) Cohort reduction, validated MGMT linkage, fixed development/held-out split, and the full and perturbation-filtered feature spaces. b) Feature counts through Direct screening, RFECV target estimation, five completed GA runs, and strict regret-constrained locking. c) Mean pairwise Jaccard and chance-corrected Nogueira agreement across five seeds. d) Paired held-out AUROC differences for locked-minus-Direct and perturbation-filtered-minus-full locked signatures, with 95% class-stratified bootstrap intervals. Both feature spaces use the same 197 participants and split; the filtered-space contrast is a label-independent feature-definition sensitivity analysis, not external validation. All paired intervals include zero. Five-seed summaries do not estimate participant-resampling or biomarker stability, and automatic tumor-core regions were not manually adjudicated.

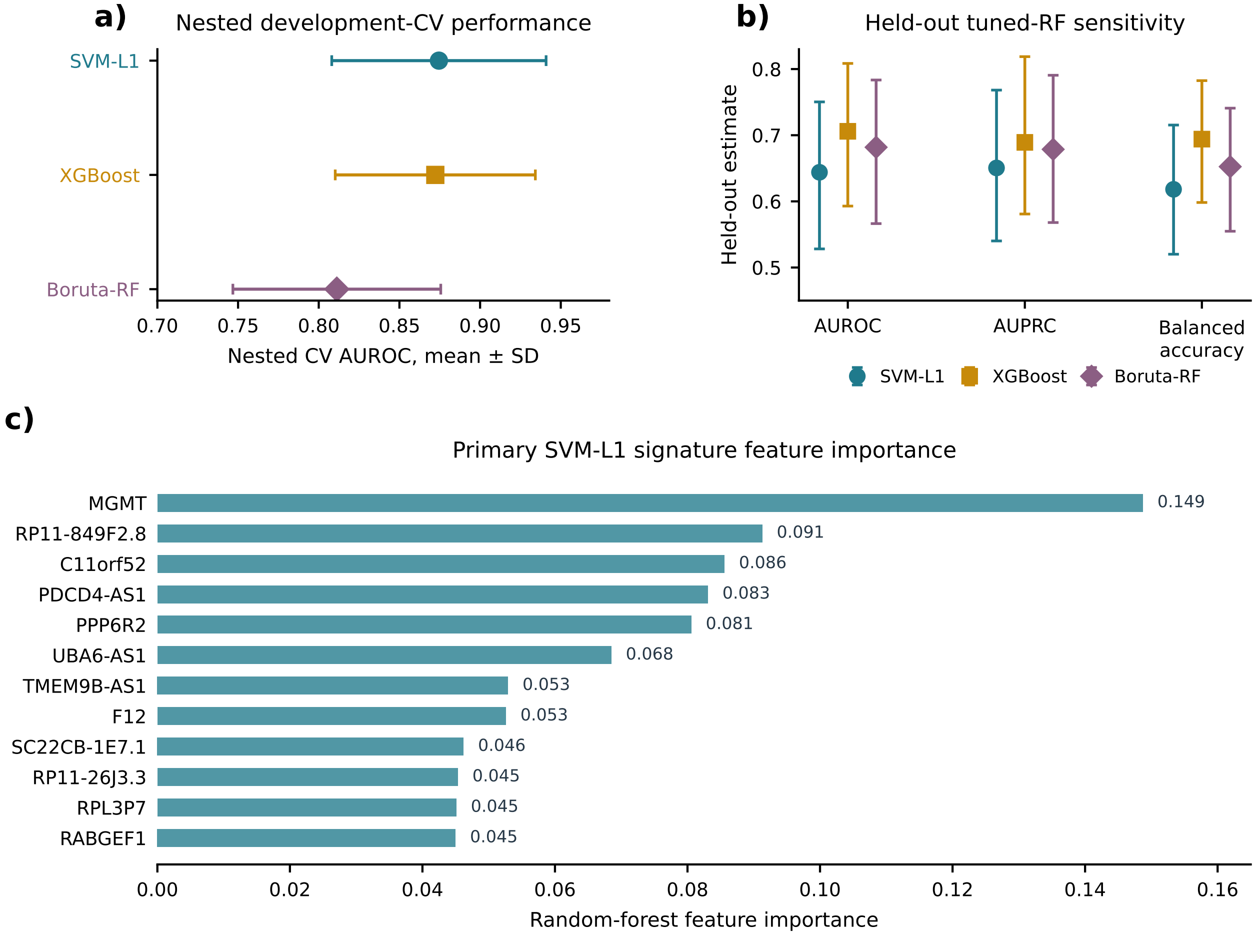


**Supplementary Figure S21.** Secondary tuned-random-forest sensitivity. a) Nested development-CV AUROC for the three branches, with the prespecified primary branch identified. b) Held-out tuned-random-forest AUROC, AUPRC, and balanced accuracy with uncertainty intervals. c) Random-forest importance values for the primary SVM-L1 signature. This analysis is descriptive and is not used to support a core claim.

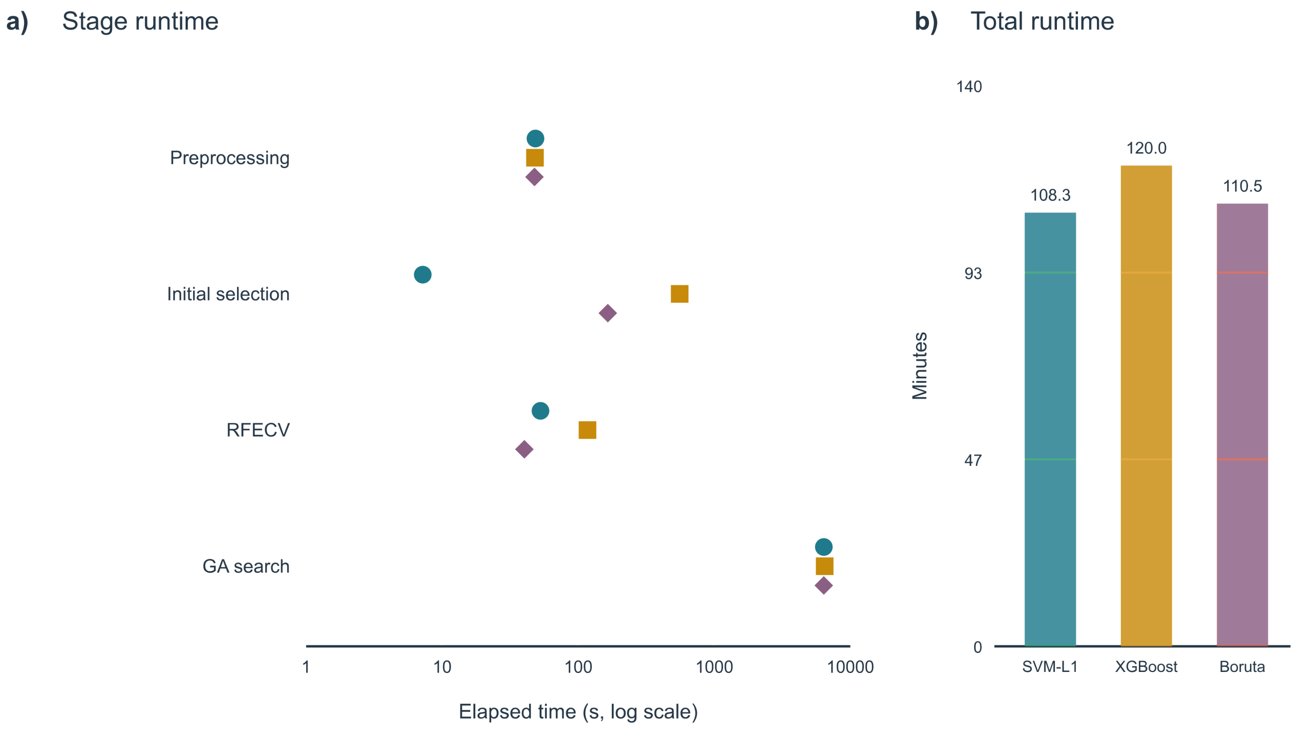


**Supplementary Figure S22.** CGGA computational characterization. a) Elapsed time for preprocessing, initial selection, RFECV, and GA search by branch on a logarithmic scale. b) Total runtime by branch. Complete runtime and peak-memory values are retained in the machine-readable computational audit.

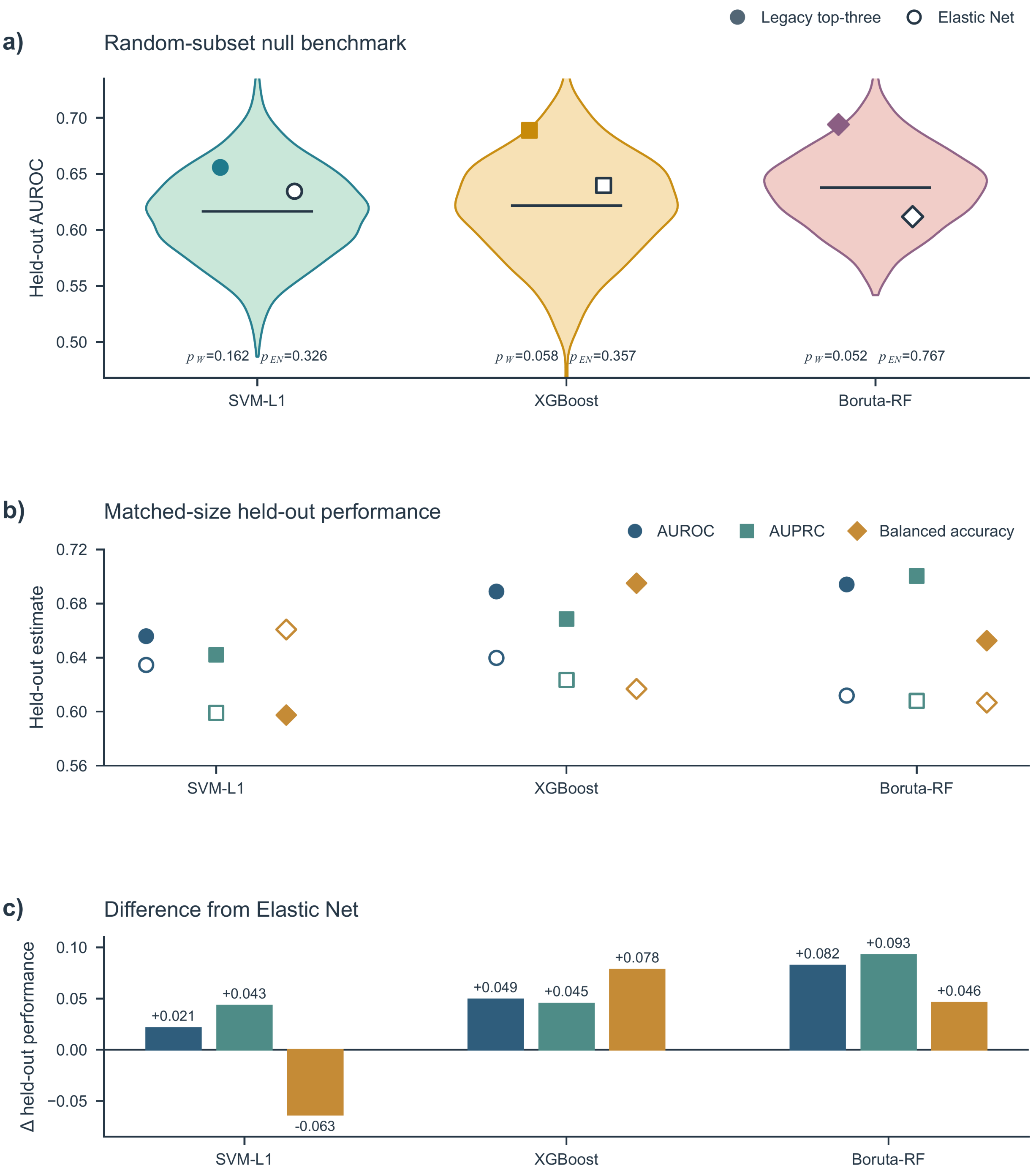


**Supplementary Figure S23.** Exploratory CGGA comparators. a) Size-matched random-subset null distributions with the legacy top-three medoid and Elastic Net estimates overlaid. b) Matched-size held-out AUROC, AUPRC, and balanced accuracy for the legacy top-three medoid and Elastic Net. c) Legacy-top-three-medoid-minus-Elastic Net differences for the same three metrics. One-sided empirical random-subset tail probabilities were not multiplicity adjusted, and all paired intervals include zero.

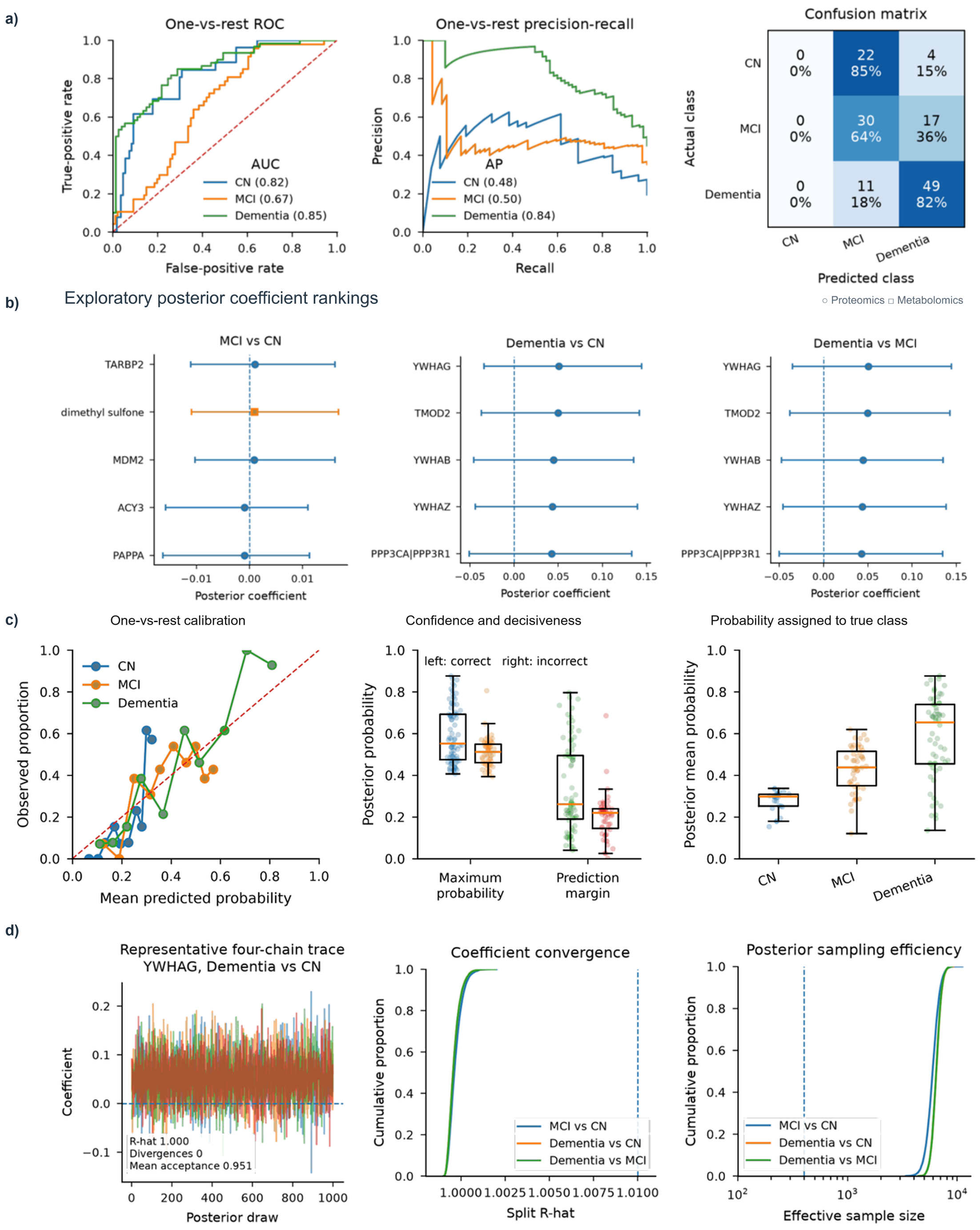


**Supplementary Figure S24.** Full-feature Bayesian predictive reference. a) One-vs-rest ROC and precision-recall curves together with the row-normalized confusion matrix. b) Exploratory posterior coefficient rankings for the three dementia contrasts; points show posterior means and intervals show 95% credible intervals. c) One-vs-rest calibration, confidence and prediction-margin distributions, and posterior probability assigned to the true class. d) A representative four-chain coefficient trace, the empirical distribution of split R-hat values, and bulk effective sample sizes. Posterior probability rankings retained discrimination while coefficient-level intervals reflected diffuse high-dimensional uncertainty.

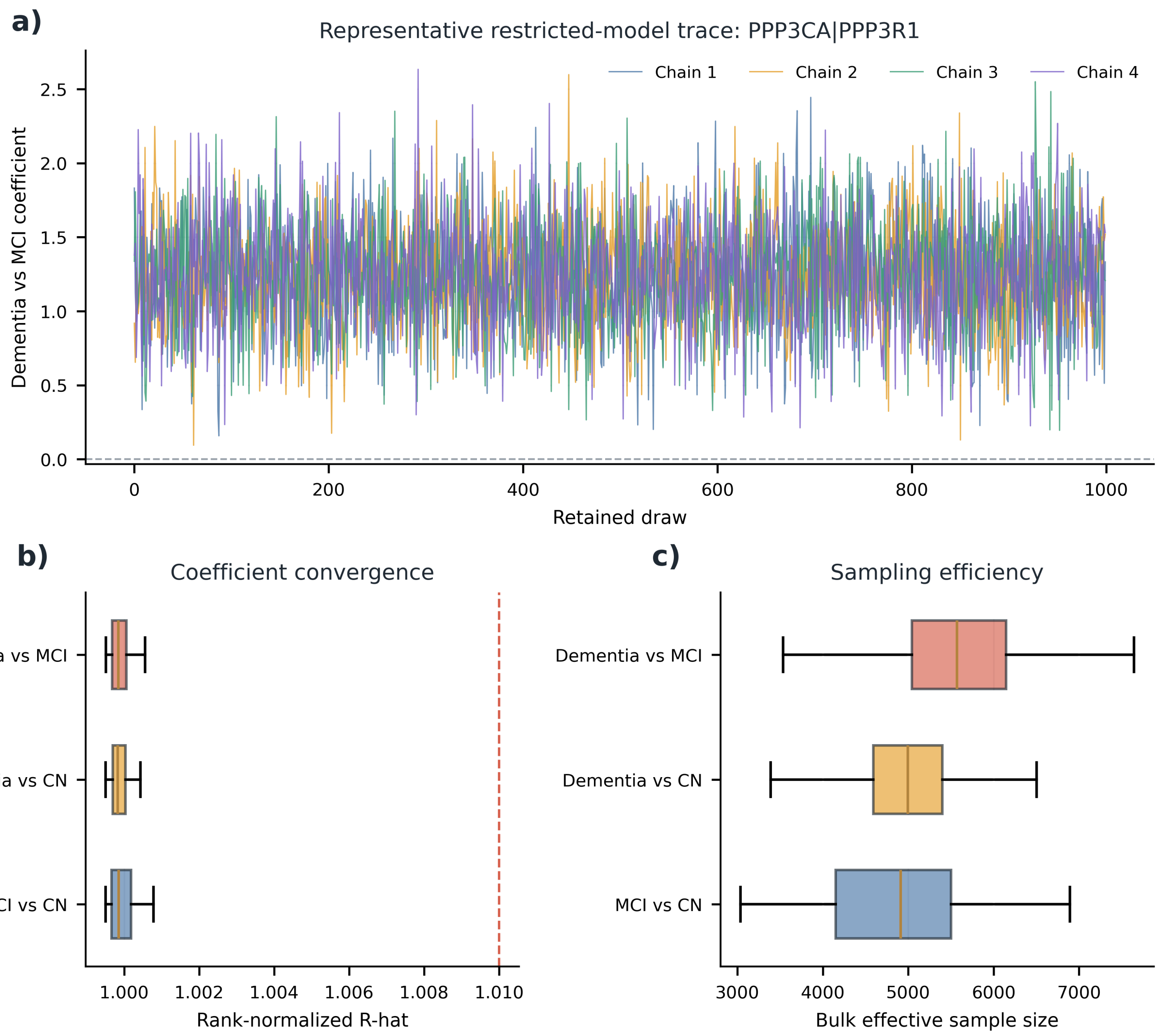


**Supplementary Figure S25.** Restricted Bayesian sampling diagnostics for the development-derived 81-candidate model. a) Representative four-chain trace for the PPP3CA|PPP3R1 dementia-versus-MCI coefficient. b) Rank-normalized R-hat distributions for the three contrasts, with the dashed line at 1.01. c) Bulk effective-sample-size distributions for the three contrasts.

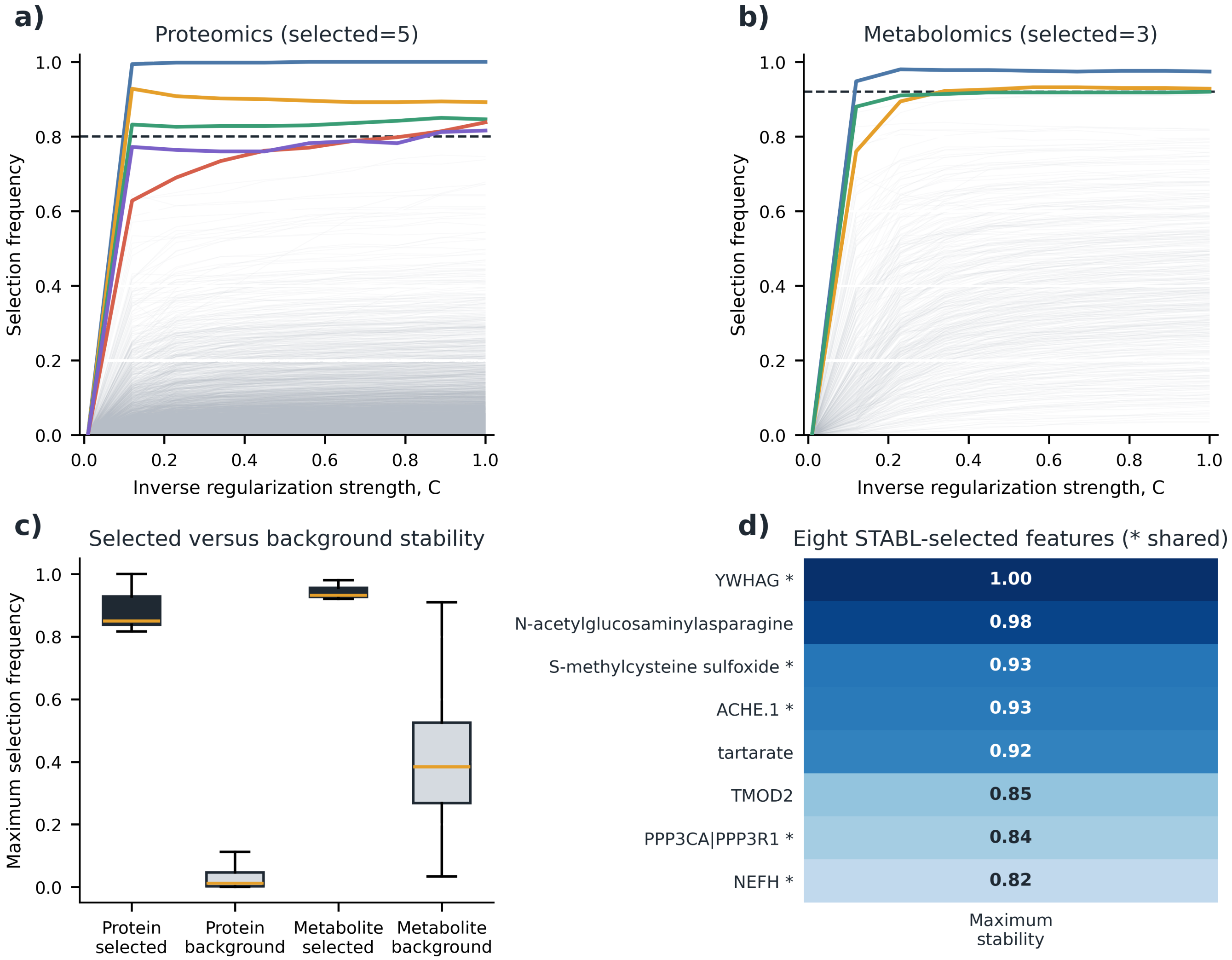


**Supplementary Figure S26.** Training-only STABL analysis. a) Proteomic and b) metabolomic stability paths across inverse regularization strength; colored paths are selected features and dashed lines are selection thresholds. c) Maximum selection-frequency distributions for selected and background features in each omic block. d) Maximum stability for the eight STABL-selected features; asterisks identify features shared with the WrapEvoFS locked-signature union.

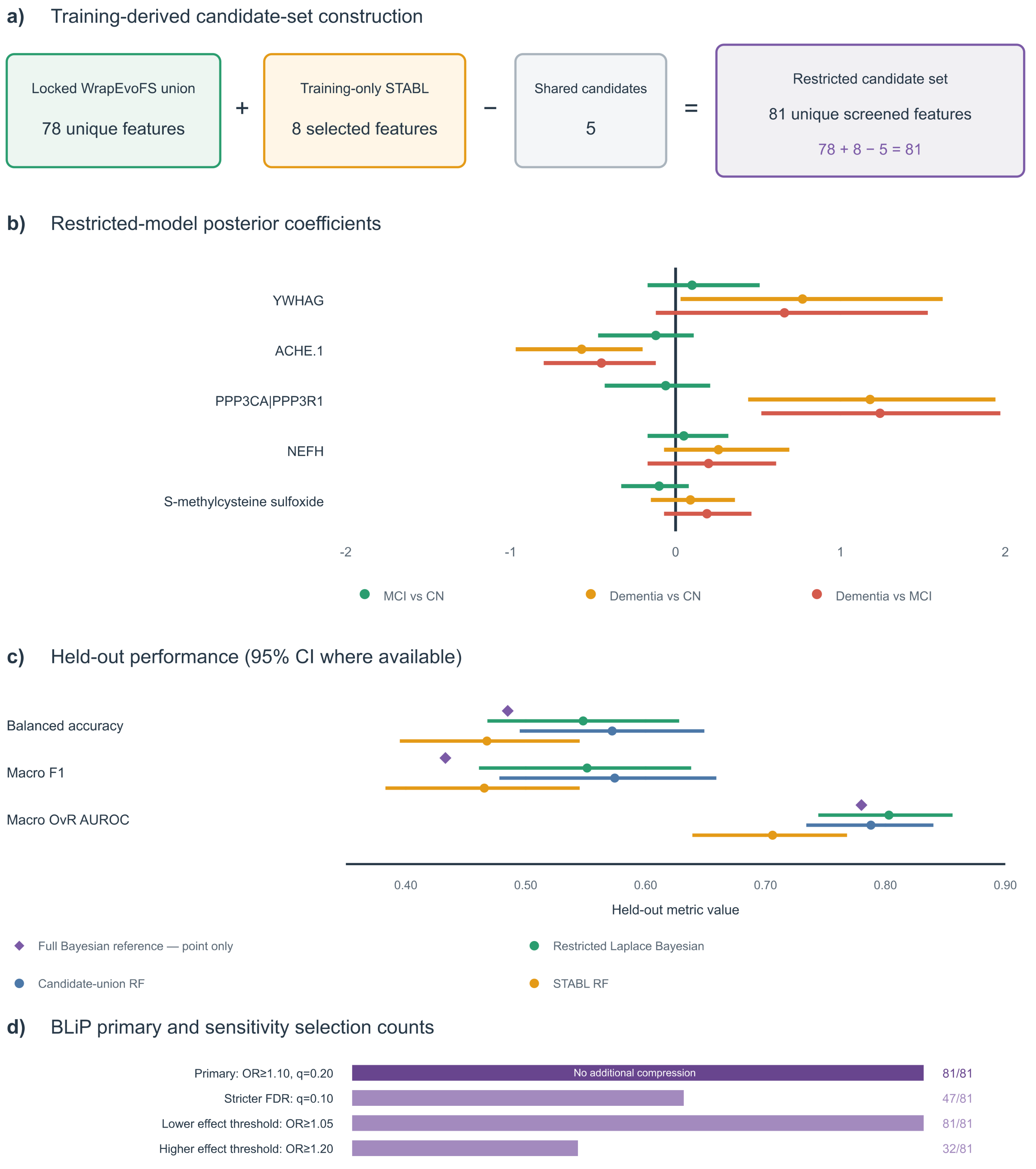


**Supplementary Figure S27.** Integrated candidate-restricted Bayesian/STABL/BLiP analysis. a) Training-derived construction of the 81-candidate restricted set from the 78-feature WrapEvoFS union and eight STABL selections with five shared candidates. b) Restricted-model posterior coefficient summaries for the five shared candidates across three dementia contrasts. c) Held-out balanced accuracy, macro F1, and macro one-vs-rest AUROC for the full Bayesian reference, restricted Laplace Bayesian model, candidate-union random forest, and STABL random forest. d) BLiP selection counts under the primary and sensitivity settings. Primary BLiP retained 81 candidates; stricter-FDR and larger-effect settings retained 47 and 32.

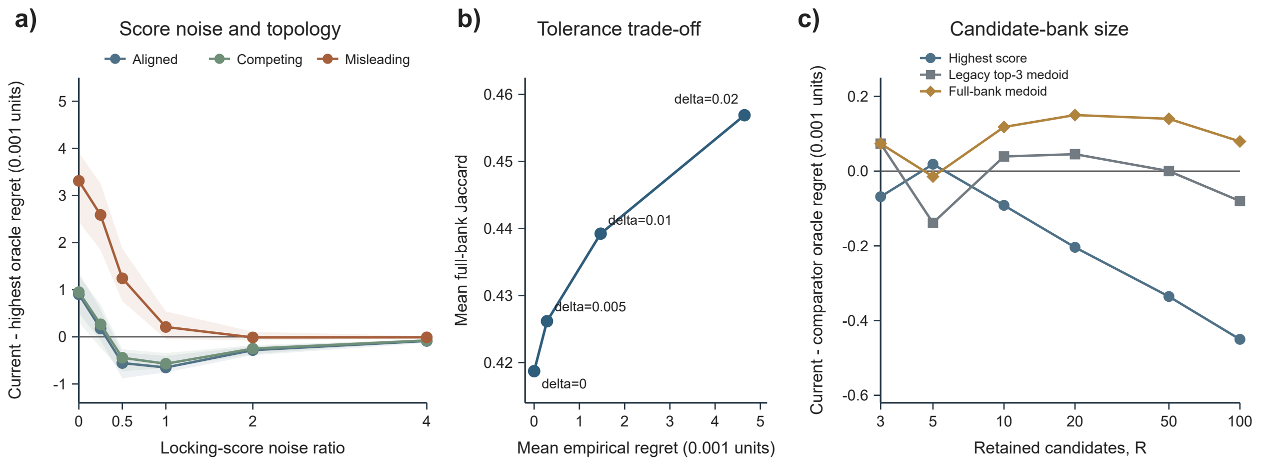


**Supplementary Figure S28.** Controlled synthetic candidate-bank locking simulation. a) Primary equal-scenario mean regret-constrained-medoid-minus-highest-score hidden-oracle regret across locking-score noise ratios and candidate topologies; shaded bands show the interquartile range across nine heterogeneity–proxy combinations. Negative differences favor the regret-constrained medoid. b) Matched 27-scenario tolerance sensitivity showing mean empirical regret against mean full-bank Jaccard; labels identify $\delta$, and increasing tolerance enlarged the eligible pool from 1.00 to 3.69 candidates on average. c) Matched 27-scenario regret-constrained-medoid-minus-comparator hidden-oracle regret across candidate-bank sizes; negative differences favor the regret-constrained medoid. The primary $R = 5$, $\delta = 0.01$ grid comprised 1,620,000 banks, and the complete sensitivity workload comprised 4,995,000. The regret-constrained medoid had no configured empirical-regret violation. Hidden utility is not AUROC; no GA or classifier was fitted.

#### S3 Supplementary Tables

**Supplementary Table S1.** Development-only AMP-AD comparison of the original configuration (zero-truncated objective and top-three Jaccard-medoid locking) with the corrected-objective configuration (untruncated objective and regret-constrained medoid locking at absolute $\delta = 0.01$) across 24 combinations of center, branch, and cap.

**a. Aggregate target fidelity and flattening diagnostics**

| Group | Absolute target deviation | | All-zero generations | |
|---|---|---|---|---|
| | Orig. | Upd. | Orig. | Upd. |
| Emory | 22 | 15 | 78 | 44 |
| Mayo | 40 | 40 | 132 | 91 |
| Mount Sinai | 44 | 34 | 101 | 65 |
| Rush | 110 | 48 | 362 | 133 |
| SVM-L1 | 119 | 62 | 369 | 150 |
| XGBoost | 57 | 62 | 235 | 134 |
| Boruta-RF | 40 | 13 | 69 | 49 |
| Overall | 216 | 137 | 673 | 333 |

**b. Complete condition-level target and flattening results**

| Center | Branch | Cap | $k^*$ | Locked $n$ | | $\|n - k^*\|$ | | Zero run-best, $n/5$ | | All-zero generations | | Upd. regret |
|---|---|---|---|---|---|---|---|---|---|---|---|---|
| | | | | Orig. | Upd. | Orig. | Upd. | Orig. | Upd. | Orig. | Upd. | |
| Emory | SVM-L1 | Small | 10 | 12 | 11 | 2 | 1 | 0/5 | 0/5 | 0 | 0 | 0.0000 |
| Emory | SVM-L1 | Reference | 19 | 19 | 19 | 0 | 0 | 0/5 | 0/5 | 0 | 0 | 0.0083 |
| Emory | XGBoost | Small | 9 | 22 | 20 | 13 | 11 | 0/5 | 0/5 | 55 | 29 | 0.0000 |
| Emory | XGBoost | Reference | 17 | 20 | 20 | 3 | 3 | 0/5 | 0/5 | 23 | 15 | 0.0000 |
| Emory | Boruta-RF | Small | 12 | 16 | 12 | 4 | 0 | 0/5 | 0/5 | 0 | 0 | 0.0000 |
| Emory | Boruta-RF | Reference | 25 | 25 | 25 | 0 | 0 | 0/5 | 0/5 | 0 | 0 | 0.0000 |
| Mayo | SVM-L1 | Small | 10 | 26 | 28 | 16 | 18 | 0/5 | 0/5 | 55 | 37 | 0.0000 |
| Mayo | SVM-L1 | Reference | 20 | 23 | 28 | 3 | 8 | 0/5 | 0/5 | 24 | 19 | 0.0000 |
| Mayo | XGBoost | Small | 10 | 22 | 22 | 12 | 12 | 0/5 | 0/5 | 37 | 21 | 0.0000 |
| Mayo | XGBoost | Reference | 20 | 20 | 22 | 0 | 2 | 0/5 | 0/5 | 11 | 9 | 0.0000 |
| Mayo | Boruta-RF | Small | 13 | 22 | 13 | 9 | 0 | 0/5 | 0/5 | 5 | 5 | 0.0000 |
| Mayo | Boruta-RF | Reference | 24 | 24 | 24 | 0 | 0 | 0/5 | 0/5 | 0 | 0 | 0.0020 |
| Mount Sinai | SVM-L1 | Small | 10 | 13 | 12 | 3 | 2 | 0/5 | 0/5 | 0 | 0 | 0.0000 |
| Mount Sinai | SVM-L1 | Reference | 17 | 19 | 17 | 2 | 0 | 0/5 | 0/5 | 0 | 0 | 0.0037 |
| Mount Sinai | XGBoost | Small | 10 | 20 | 24 | 10 | 14 | 0/5 | 0/5 | 31 | 16 | 0.0000 |
| Mount Sinai | XGBoost | Reference | 19 | 22 | 24 | 3 | 5 | 0/5 | 0/5 | 6 | 5 | 0.0000 |
| Mount Sinai | Boruta-RF | Small | 13 | 32 | 24 | 19 | 11 | 0/5 | 0/5 | 39 | 29 | 0.0060 |
| Mount Sinai | Boruta-RF | Reference | 22 | 29 | 24 | 7 | 2 | 0/5 | 0/5 | 25 | 15 | 0.0060 |
| Rush | SVM-L1 | Small | 3 | 84 | 28 | 81 | 25 | 4/5 | 0/5 | 245 | 66 | 0.0000 |
| Rush | SVM-L1 | Reference | 18 | 30 | 26 | 12 | 8 | 0/5 | 0/5 | 45 | 28 | 0.0000 |
| Rush | XGBoost | Small | 7 | 23 | 21 | 16 | 14 | 0/5 | 0/5 | 59 | 30 | 0.0000 |
| Rush | XGBoost | Reference | 20 | 20 | 21 | 0 | 1 | 0/5 | 0/5 | 13 | 9 | 0.0000 |
| Rush | Boruta-RF | Small | 12 | 13 | 12 | 1 | 0 | 0/5 | 0/5 | 0 | 0 | 0.0000 |
| Rush | Boruta-RF | Reference | 23 | 23 | 23 | 0 | 0 | 0/5 | 0/5 | 0 | 0 | 0.0000 |

**Supplementary Table S1.** Development-only AMP-AD configuration comparison (continued).

**c. Fixed development-CV locking score, eligibility, and agreement**

| Center | Branch | Cap | Locking score | | Difference | Eligible $E$ | Upd. regret | Upd. mean Jaccard |
|---|---|---|---|---|---|---|---|---|
| | | | Orig. | Upd. | Upd. − original | | | |
| Emory | SVM-L1 | Small | 0.8633 | 0.8501 | -0.0132 | 3 | 0.0000 | 0.1721 |
| Emory | SVM-L1 | Reference | 0.8629 | 0.8609 | -0.0020 | 4 | 0.0083 | 0.3114 |
| Emory | XGBoost | Small | 0.8194 | 0.8235 | 0.0041 | 3 | 0.0000 | 0.1319 |
| Emory | XGBoost | Reference | 0.8207 | 0.8235 | 0.0028 | 3 | 0.0000 | 0.1476 |
| Emory | Boruta-RF | Small | 0.8447 | 0.8488 | 0.0041 | 3 | 0.0000 | 0.1941 |
| Emory | Boruta-RF | Reference | 0.8585 | 0.8629 | 0.0044 | 3 | 0.0000 | 0.3007 |
| Mayo | SVM-L1 | Small | 0.8803 | 0.8803 | -0.0000 | 3 | 0.0000 | 0.1684 |
| Mayo | SVM-L1 | Reference | 0.8800 | 0.8803 | 0.0003 | 3 | 0.0000 | 0.1684 |
| Mayo | XGBoost | Small | 0.8621 | 0.8508 | -0.0113 | 2 | 0.0000 | 0.1795 |
| Mayo | XGBoost | Reference | 0.8269 | 0.8508 | 0.0239 | 2 | 0.0000 | 0.1795 |
| Mayo | Boruta-RF | Small | 0.8705 | 0.8694 | -0.0011 | 2 | 0.0000 | 0.1154 |
| Mayo | Boruta-RF | Reference | 0.8824 | 0.8767 | -0.0058 | 5 | 0.0020 | 0.2407 |
| Mount Sinai | SVM-L1 | Small | 0.8760 | 0.8796 | 0.0036 | 3 | 0.0000 | 0.3125 |
| Mount Sinai | SVM-L1 | Reference | 0.8862 | 0.8920 | 0.0058 | 5 | 0.0037 | 0.2732 |
| Mount Sinai | XGBoost | Small | 0.8613 | 0.8722 | 0.0109 | 2 | 0.0000 | 0.1250 |
| Mount Sinai | XGBoost | Reference | 0.8614 | 0.8722 | 0.0108 | 2 | 0.0000 | 0.1250 |
| Mount Sinai | Boruta-RF | Small | 0.8856 | 0.8804 | -0.0052 | 3 | 0.0060 | 0.1786 |
| Mount Sinai | Boruta-RF | Reference | 0.8821 | 0.8804 | -0.0017 | 3 | 0.0060 | 0.1786 |
| Rush | SVM-L1 | Small | 0.8701 | 0.8602 | -0.0100 | 3 | 0.0000 | 0.1561 |
| Rush | SVM-L1 | Reference | 0.8827 | 0.8735 | -0.0092 | 1 | 0.0000 | NA |
| Rush | XGBoost | Small | 0.8652 | 0.8622 | -0.0030 | 2 | 0.0000 | 0.0833 |
| Rush | XGBoost | Reference | 0.8595 | 0.8631 | 0.0036 | 3 | 0.0000 | 0.1676 |
| Rush | Boruta-RF | Small | 0.8766 | 0.8782 | 0.0017 | 3 | 0.0000 | 0.2982 |
| Rush | Boruta-RF | Reference | 0.8756 | 0.8856 | 0.0100 | 1 | 0.0000 | NA |

*Note.* Orig., original configuration; Upd., corrected-objective configuration. The original configuration used the zero-truncated objective and legacy top-three medoid. Corrected-objective values were obtained from 120 newly executed full GA runs (five seeds per condition) at the Small and Reference caps using the untruncated objective and regret-constrained medoid; High-cap conditions were not rerun. The all-zero column is the zero-truncation diagnostic evaluated along each untruncated search history. Uniform parent-sampling fallbacks were zero. Final locking used fixed development-CV scores, absolute regret tolerance 0.01, strict eligible-only pools, and no fallback expansion. Because both candidate generation and locking differ, between-column differences are configuration-level comparisons rather than isolated causal effects of either component. The two Rush singleton pools have undefined mean Jaccard (NA); all 18 newly completed pools contained two to five candidates. Held-out outcomes were not accessed or evaluated. Mean Jaccard was reduced in canonical mask-hash order. In Mount Sinai/SVM-L1/Reference, runs 3 and 5 had the same exact peer-Jaccard multiset but differed by one IEEE-754 unit in the last place under historical versus canonical reduction order; the authoritative package selected run 5 (17 features; regret 0.0037), whereas the older helper selected run 3 (17 features; regret 0). Both were eligible, and target-count summaries were unchanged.

**Supplementary Table S2.** Robust development-only summary of the 24-condition AMP-AD original-versus-corrected-objective configuration comparison.

| Scope | Diagnostic | $n$ | Original | Corrected | Reduction (%) | Median improvement | Better/unchanged/worse |
|---|---|---|---|---|---|---|---|
| All 24 conditions | Absolute target deviation | 24 | 216 | 137 | 79 (36.6%) | 0.5 | 12/6/6 |
| All 24 conditions | All-zero generations | 24 | 673 | 333 | 340 (50.5%) | 3.0 | 14/10/0 |
| Excluding Rush/SVM-L1/Small | Absolute target deviation | 23 | 135 | 112 | 23 (17.0%) | 0.0 | 11/6/6 |
| Excluding Rush/SVM-L1/Small | All-zero generations | 23 | 428 | 267 | 161 (37.6%) | 2.0 | 13/10/0 |

*Note.* The excluded prespecified stress condition is Rush/SVM-L1/Small. Positive improvement denotes a lower corrected-objective value. The stress condition accounts for 56/79 (70.9%) of the aggregate target-deviation reduction and 179/340 (52.6%) of the all-zero-generation reduction; both diagnostics remain lower after exclusion. Held-out outcomes were not accessed.

**Supplementary Table S3.** Existing-artifact 2 by 2 candidate-bank and locking-rule comparison for matched AMP-AD development conditions.

| Candidate bank | Locking rule | $n$ | Total target deviation | Mean selected $n$ | Mean score | Mean regret | Maximum regret | Mask changes |
|---|---|---|---|---|---|---|---|---|
| Original-objective bank | Original top-three | 24 | 216 | 24.12 | 0.86475 | 0.00261 | 0.02111 | 10 |
| Original-objective bank | Regret-constrained | 24 | 227 | 24.58 | 0.86676 | 0.00060 | 0.00729 | 10 |
| Corrected-objective bank | Legacy top-three | 24 | 135 | 20.75 | 0.86329 | 0.00351 | 0.01525 | 8 |
| Corrected-objective bank | Regret-constrained | 24 | 137 | 20.83 | 0.86572 | 0.00108 | 0.00835 | 8 |

*Note.* No candidate generation or held-out evaluation was rerun. All 24 original-objective banks and all 24 corrected-objective banks were complete after recovery of the 30 Rush run-best feature lists from the authoritative Drive source folder. "Mask changes" is the within-bank number of conditions for which the two locking rules selected different masks. The regret-constrained rule improved the configured development-CV score-gap summaries in both bank sets but did not universally improve target fidelity.

**Supplementary Table S4.** Eligible-pool decision paths and duplicate-mask sensitivity for the 24 corrected-objective AMP-AD development-only conditions at absolute $\delta = 0.01$.

**a. Eligible-pool and decision-path audit**

| Pool category | Conditions | Percent | Decision stage | Conditions | Percent |
|---|---|---|---|---|---|
| Singleton | 2 | 8.3% | Singleton direct | 2 | 8.3% |
| Two candidates | 6 | 25.0% | Unique Jaccard | 13 | 54.2% |
| Three or more | 16 | 66.7% | Higher score | 9 | 37.5% |
| | | | Smaller feature count | 0 | 0.0% |
| | | | Stable mask hash | 0 | 0.0% |

**b. Duplicate-retained versus deduplicated sensitivity**

| Candidate bank | Audited conditions | Banks with duplicates | Selected-mask changes | Status |
|---|---|---|---|---|
| Original-objective | 24 | 0 | 0 | Complete |
| Corrected-objective | 24 | 0 | 0 | Complete after Rush recovery |

*Note.* Exact duplicate masks are retained as multiple voting candidates by the package. No duplicates occurred in the 48 complete banks, so deduplication did not change a selected mask in this empirical sensitivity. The 30 Rush run-best feature lists were recovered from the authoritative Drive source folder; all six current selections matched both the archived summary and the frozen one-time evaluation. Source-run provenance is distinct from input-row order and is used only after scientific feature-set selection when exact duplicates require a provenance representative.

**Supplementary Table S5.** AMP-AD development-only representative-run locking sensitivity across 36 completed conditions.

| **Branch** | **Conditions** | **Legacy top-three = highest score** | **Legacy top-three = full-bank medoid** | **Legacy CV regret, median [IQR]** | **Legacy median features** |
|---|---|---|---|---|---|
| XGBoost | 12 | 8/12 | 4/12 | 0.0000 [0.0000, 0.0025] | 22.0 |
| SVM-L1 | 12 | 5/12 | 5/12 | 0.0016 [0.0000, 0.0037] | 28.0 |
| Boruta-RF | 12 | 8/12 | 5/12 | 0.0000 [0.0000, 0.0021] | 27.0 |
| Overall | 36 | 21/36 | 14/36 | 0.0000 [0.0000, 0.0030] | 24.5 |

**Supplementary Table S6.** Fully nested TCGA selector-layer comparator and random-bank sensitivity.

| Branch | Selector | Mean features | Macro AUROC | 95% CI | WrapEvoFS minus selector |
|---|---|---|---|---|---|
| SVM-L1 | WrapEvoFS | 14.6 | 0.827 | 0.807–0.845 | Reference |
| | Elastic Net, native | 73.7 | 0.841 | 0.821–0.861 | −0.0146 [−0.0251, −0.0045] |
| | Elastic Net, matched | 14.6 | 0.809 | 0.789–0.827 | +0.0176 [+0.0008, +0.0341] |
| | Stability, native | 58.0 | 0.840 | 0.820–0.859 | −0.0136 [−0.0245, −0.0029] |
| | Stability, matched | 14.6 | 0.800 | 0.780–0.820 | +0.0261 [+0.0082, +0.0434] |
| XGBoost | WrapEvoFS | 20.0 | 0.827 | 0.806–0.847 | Reference |
| | Elastic Net, native | 77.0 | 0.834 | 0.814–0.854 | −0.0071 [−0.0173, +0.0037] |
| | Elastic Net, matched | 20.0 | 0.829 | 0.809–0.848 | −0.0017 [−0.0146, +0.0111] |
| | Stability, native | 38.2 | 0.826 | 0.807–0.845 | +0.0008 [−0.0102, +0.0119] |
| | Stability, matched | 20.0 | 0.829 | 0.809–0.847 | −0.0015 [−0.0136, +0.0105] |
| Boruta-RF | WrapEvoFS | 18.0 | 0.828 | 0.807–0.847 | Reference |
| | Elastic Net, native | 60.4 | 0.838 | 0.819–0.857 | −0.0108 [−0.0199, −0.0012] |
| | Elastic Net, matched | 18.0 | 0.839 | 0.819–0.857 | −0.0109 [−0.0236, +0.0024] |
| | Stability, native | 30.6 | 0.834 | 0.814–0.852 | −0.0066 [−0.0179, +0.0057] |
| | Stability, matched | 18.0 | 0.841 | 0.821–0.860 | −0.0128 [−0.0260, +0.0012] |

Macro one-vs-rest AUROC estimates average two repeated five-fold outer-CV OOF estimates. Intervals and paired differences use 2,000 participant-clustered, class-stratified resamples retaining both repeat predictions for each sampled participant. Elastic Net and stability selection were fitted inside each verified post-Direct outer-training universe. Native support is the prespecified primary comparator; matched support uses the same outer-condition cardinality as WrapEvoFS. The five random-bank means were 0.828, 0.817, and 0.832 for SVM-L1, XGBoost, and Boruta-RF, respectively; mean WrapEvoFS-minus-random differences were −0.0018, +0.0102, and −0.0043. Random controls match five-candidate bank size and locking-evaluation budget, not full GA compute. These retrospective selector-layer sensitivities are conditional on the verified post-Direct outer-training universes; equivalence and external validity were not assessed.

**Supplementary Table S7.** AMP-AD one-time post-freeze held-out evaluation of the 24 corrected-objective conditions

| Branch | Cap | Mean center AUROC | Medoid − RFECV-only | Medoid − legacy top-three |
|---|---|---|---|---|
| SVM-L1 | Small | 0.747 | -0.010 [-0.035, +0.014] | -0.031 [-0.055, -0.006] |
| SVM-L1 | Reference | 0.776 | -0.008 [-0.029, +0.012] | +0.012 [-0.009, +0.035] |
| XGBoost | Small | 0.785 | +0.048 [+0.025, +0.070] | +0.018 [-0.003, +0.038] |
| XGBoost | Reference | 0.782 | -0.021 [-0.044, +0.001] | +0.015 [-0.007, +0.037] |
| Boruta-RF | Small | 0.779 | -0.040 [-0.062, -0.019] | -0.012 [-0.033, +0.013] |
| Boruta-RF | Reference | 0.793 | +0.002 [-0.015, +0.019] | +0.015 [-0.003, +0.033] |

Point estimates in the fourth and fifth columns are pooled four-center macro-AUROC differences with 95% intervals from 2,000 common resamples stratified by held-out center and true class (seed 42). The 24 development locks and protocol hash were frozen before held-out labels or outcomes were loaded. The evaluation was performed once and did not trigger reselection or retuning. This is post-freeze cross-center evaluation, not prospective clinical or independent external validation.

**Supplementary Table S8.** Archived ADNI locking audit and agreement summary. Section a reports the five run-level development-CV candidates under legacy and regret-constrained locking; section b reports the selected-run agreement summary. The larger-is-better locking score is mean balanced accuracy, and absolute regret is measured from the branch maximum. At $\delta = 0.01$, each branch had one eligible candidate, marked $\delta$-selected.

**a. Five-run locking audit**

| **Method** | **GA rank** | **n** | **Balanced accuracy** | **Macro F1** | **Weighted OvR AUROC** | **Lock rank** | **Abs. regret** | **Status** |
|---|---|---|---|---|---|---|---|---|
| SVM-L1 | 1 | 29 | 0.563 (0.028) | 0.569 (0.033) | 0.814 (0.011) | 3 | 0.02365 | Legacy top 3 |
| SVM-L1 | 2 | 29 | 0.572 (0.055) | 0.579 (0.056) | 0.817 (0.031) | 2 | 0.01401 | Legacy selected |
| SVM-L1 | 3 | 29 | 0.562 (0.053) | 0.568 (0.049) | 0.818 (0.023) | 4 | 0.02463 | — |
| SVM-L1 | 4 | 29 | 0.586 (0.052) | 0.595 (0.049) | 0.808 (0.040) | 1 | 0 | Legacy top 3; $\delta$-selected |
| SVM-L1 | 5 | 29 | 0.560 (0.029) | 0.565 (0.035) | 0.820 (0.018) | 5 | 0.02596 | — |
| XGBoost | 1 | 30 | 0.521 (0.032) | 0.520 (0.036) | 0.768 (0.008) | 3 | 0.02168 | Legacy top 3 |
| XGBoost | 2 | 30 | 0.542 (0.054) | 0.545 (0.055) | 0.784 (0.034) | 1 | 0 | Legacy and $\delta$-selected |
| XGBoost | 3 | 31 | 0.517 (0.038) | 0.509 (0.043) | 0.767 (0.019) | 4 | 0.02570 | — |
| XGBoost | 4 | 31 | 0.530 (0.047) | 0.534 (0.052) | 0.772 (0.015) | 2 | 0.01232 | Legacy top 3 |
| XGBoost | 5 | 31 | 0.516 (0.031) | 0.513 (0.034) | 0.768 (0.027) | 5 | 0.02618 | — |
| Boruta-RF | 1 | 27 | 0.574 (0.021) | 0.580 (0.026) | 0.788 (0.020) | 1 | 0 | Legacy top 3; $\delta$-selected |
| Boruta-RF | 2 | 27 | 0.557 (0.020) | 0.563 (0.025) | 0.791 (0.015) | 2 | 0.01713 | Legacy top 3 |
| Boruta-RF | 3 | 27 | 0.551 (0.024) | 0.555 (0.025) | 0.783 (0.023) | 3 | 0.02252 | Legacy selected |
| Boruta-RF | 4 | 27 | 0.530 (0.030) | 0.533 (0.029) | 0.780 (0.025) | 5 | 0.04439 | — |
| Boruta-RF | 5 | 27 | 0.548 (0.016) | 0.554 (0.024) | 0.780 (0.017) | 4 | 0.02553 | — |

**b. Locked-medoid agreement summary**

| Method | Locked run | Mean Jaccard with top three |
|---|---|---|
| SVM-L1 | 2 | 0.382 |
| XGBoost | 2 | 0.247 |
| Boruta-RF | 3 | 0.385 |

**Supplementary Table S9.** ADNI held-out performance and paired archived legacy-top-three-medoid-minus-Direct differences. Sections a–d report development-CV summaries, held-out point estimates, class-specific sensitivity, and paired held-out differences, respectively. In this table, "Locked medoid" denotes the archived legacy top-three medoid. The development-CV summaries here reproduce the archived final-evaluation fold summary; Supplementary Table S8 reports the separately recovered run-level locking-evaluator scores used for re-locking. Values may therefore differ for the same archived mask and should not be interchanged.

**a. Development-CV summary**

| Method | Signature | n | Balanced accuracy | Macro F1 | Weighted OvR AUROC | Macro OvR AUROC |
|---|---|---|---|---|---|---|
| SVM-L1 | Direct | 63 | 0.547 (0.044) | 0.535 (0.058) | 0.825 (0.022) | 0.821 (0.016) |
| SVM-L1 | Locked medoid | 29 | 0.572 (0.055) | 0.579 (0.056) | 0.817 (0.031) | 0.813 (0.025) |
| XGBoost | Direct | 100 | 0.509 (0.041) | 0.502 (0.048) | 0.783 (0.023) | 0.781 (0.023) |
| XGBoost | Locked medoid | 30 | 0.539 (0.038) | 0.539 (0.037) | 0.787 (0.032) | 0.785 (0.024) |
| Boruta-RF | Direct | 48 | 0.549 (0.028) | 0.555 (0.036) | 0.786 (0.016) | 0.784 (0.016) |
| Boruta-RF | Locked medoid | 27 | 0.560 (0.058) | 0.565 (0.063) | 0.779 (0.032) | 0.779 (0.027) |

**b. Held-out point estimates**

| Method | Signature | Balanced accuracy | Macro F1 | Accuracy | Weighted OvR AUROC | Macro OvR AUROC |
|---|---|---|---|---|---|---|
| SVM-L1 | Direct | 0.531 [0.460, 0.603] | 0.521 [0.440, 0.606] | 0.617 [0.549, 0.684] | 0.790 [0.734, 0.843] | 0.782 [0.725, 0.833] |
| SVM-L1 | Locked medoid | 0.521 [0.446, 0.604] | 0.517 [0.431, 0.610] | 0.602 [0.526, 0.677] | 0.790 [0.729, 0.841] | 0.784 [0.724, 0.835] |
| XGBoost | Direct | 0.535 [0.469, 0.607] | 0.525 [0.442, 0.610] | 0.624 [0.556, 0.692] | 0.810 [0.754, 0.862] | 0.809 [0.757, 0.858] |
| XGBoost | Locked medoid | 0.564 [0.483, 0.646] | 0.578 [0.482, 0.663] | 0.624 [0.549, 0.699] | 0.775 [0.717, 0.831] | 0.775 [0.720, 0.831] |
| Boruta-RF | Direct | 0.577 [0.494, 0.663] | 0.590 [0.497, 0.675] | 0.624 [0.549, 0.699] | 0.794 [0.739, 0.849] | 0.790 [0.733, 0.845] |
| Boruta-RF | Locked medoid | 0.535 [0.460, 0.610] | 0.540 [0.455, 0.626] | 0.602 [0.526, 0.677] | 0.795 [0.740, 0.849] | 0.788 [0.732, 0.842] |

**c. Class-specific sensitivity**

| Method | Signature | CN | MCI | Dementia |
|---|---|---|---|---|
| SVM-L1 | Direct | 0.154 | 0.638 | 0.800 |
| SVM-L1 | Locked medoid | 0.192 | 0.553 | 0.817 |
| XGBoost | Direct | 0.154 | 0.617 | 0.833 |
| XGBoost | Locked medoid | 0.308 | 0.617 | 0.767 |
| Boruta-RF | Direct | 0.385 | 0.596 | 0.750 |
| Boruta-RF | Locked medoid | 0.269 | 0.553 | 0.783 |

**d. Paired held-out differences**

| Method | Metric | Direct | Locked medoid | Difference | 95% paired CI |
|---|---|---|---|---|---|
| SVM-L1 | Balanced accuracy | 0.531 | 0.521 | -0.010 | [-0.070, +0.053] |
| SVM-L1 | Macro F1 | 0.521 | 0.517 | -0.004 | [-0.076, +0.071] |
| SVM-L1 | Macro OvR AUROC | 0.782 | 0.784 | +0.001 | [-0.031, +0.035] |
| XGBoost | Balanced accuracy | 0.535 | 0.564 | +0.029 | [-0.047, +0.107] |
| XGBoost | Macro F1 | 0.525 | 0.578 | +0.052 | [-0.044, +0.146] |
| XGBoost | Macro OvR AUROC | 0.809 | 0.775 | -0.034 | [-0.075, +0.004] |
| Boruta-RF | Balanced accuracy | 0.577 | 0.535 | -0.042 | [-0.107, +0.021] |
| Boruta-RF | Macro F1 | 0.590 | 0.540 | -0.050 | [-0.125, +0.022] |
| Boruta-RF | Macro OvR AUROC | 0.790 | 0.788 | -0.002 | [-0.029, +0.026] |

**Supplementary Table S10.** AMP-AD pooled performance, paired-interval audit, target tracking, and cross-center recurrence. Sections a–d report pooled point estimates, interval coverage, target fidelity, and locked-signature recurrence, respectively.

**a. Pooled point estimates**

| Branch | Cap | Mean features | Macro OvR AUROC [95% CI] | Balanced accuracy [95% CI] | Macro F1 [95% CI] |
|---|---|---|---|---|---|
| XGBoost | Small | 21.8 | 0.782 [0.755, 0.810] | 0.558 [0.518, 0.600] | 0.559 [0.515, 0.603] |
| XGBoost | Reference | 20.5 | 0.747 [0.717, 0.776] | 0.528 [0.488, 0.566] | 0.521 [0.477, 0.562] |
| XGBoost | High | 33.0 | 0.784 [0.759, 0.811] | 0.542 [0.503, 0.582] | 0.534 [0.493, 0.577] |
| SVM-L1 | Small | 33.8 | 0.769 [0.741, 0.796] | 0.527 [0.486, 0.569] | 0.531 [0.487, 0.574] |
| SVM-L1 | Reference | 22.8 | 0.774 [0.745, 0.802] | 0.556 [0.518, 0.595] | 0.560 [0.519, 0.601] |
| SVM-L1 | High | 36.2 | 0.783 [0.756, 0.809] | 0.520 [0.481, 0.564] | 0.524 [0.480, 0.571] |
| Boruta-RF | Small | 20.8 | 0.763 [0.737, 0.790] | 0.523 [0.486, 0.560] | 0.520 [0.480, 0.559] |
| Boruta-RF | Reference | 25.2 | 0.770 [0.741, 0.797] | 0.555 [0.520, 0.593] | 0.543 [0.505, 0.583] |
| Boruta-RF | High | 47.0 | 0.776 [0.751, 0.801] | 0.526 [0.488, 0.564] | 0.525 [0.485, 0.564] |

**b. Paired-interval audit**

| Comparison | Metric | Intervals including zero | Conditions |
|---|---|---|---|
| Locked - Direct | macro auroc | 4 | 9 |
| Locked - Direct | balanced accuracy | 8 | 9 |
| Locked - Direct | macro f1 | 8 | 9 |
| Locked - RFECV-only | macro auroc | 4 | 9 |
| Locked - RFECV-only | balanced accuracy | 8 | 9 |
| Locked - RFECV-only | macro f1 | 8 | 9 |

**c. Target fidelity and success counts**

| Cap | Absolute deviation, features | Target attainment, n/12 |
|---|---|---|
| Small | Mean 15.50; median 11.0 [3.75, 16.0] | Exact 0; <=1: 1; <=3: 3 |
| Reference | Mean 2.50; median 1.0 [0, 3.0] | Exact 6; <=1: 6; <=3: 10 |
| High | Mean 0.75; median 0 [0, 1.0] | Exact 8; <=1: 11; <=3: 11 |

**d. Locked-signature cross-center recurrence**

| Branch | Cap | Mean pairwise Jaccard | Minimum | Maximum |
|---|---|---|---|---|
| XGBoost | Small | 0.057 | 0.047 | 0.075 |
| XGBoost | Reference | 0.043 | 0.024 | 0.053 |
| XGBoost | High | 0.040 | 0.028 | 0.052 |
| SVM-L1 | Small | 0.042 | 0.000 | 0.087 |
| SVM-L1 | Reference | 0.077 | 0.039 | 0.188 |
| SVM-L1 | High | 0.080 | 0.013 | 0.143 |
| Boruta-RF | Small | 0.086 | 0.023 | 0.143 |
| Boruta-RF | Reference | 0.084 | 0.040 | 0.143 |
| Boruta-RF | High | 0.157 | 0.094 | 0.218 |

**Supplementary Table S11.** AMP-AD ordinal outcome sensitivity using existing pooled leave-one-center-out predictions. Sections a and b report locked-medoid point estimates and paired locked-medoid-minus-Direct differences, respectively.

**a. Locked-medoid point estimates**

| Branch | Cap | Mean features | QWK | Ordinal MAE | Severe-error rate |
|---|---|---|---|---|---|
| XGBoost | Small | 21.8 | 0.598 | 0.412 | 0.042 |
| XGBoost | Reference | 20.5 | 0.521 | 0.463 | 0.072 |
| XGBoost | High | 33.0 | 0.585 | 0.429 | 0.049 |
| SVM-L1 | Small | 33.8 | 0.537 | 0.452 | 0.042 |
| SVM-L1 | Reference | 22.8 | 0.555 | 0.427 | 0.055 |
| SVM-L1 | High | 36.2 | 0.499 | 0.459 | 0.057 |

*Supplementary Table S11 (continued)*

| Branch | Cap | Mean features | QWK | Ordinal MAE | Severe-error rate |
|---|---|---|---|---|---|
| Boruta-RF | Small | 20.8 | 0.559 | 0.448 | 0.049 |
| Boruta-RF | Reference | 25.2 | 0.598 | 0.423 | 0.049 |
| Boruta-RF | High | 47.0 | 0.573 | 0.444 | 0.042 |

**b. Paired locked-medoid-minus-Direct differences**

| Branch | Cap | Delta QWK [95% CI] | Delta ordinal MAE [95% CI] | Delta severe-error rate [95% CI] |
|---|---|---|---|---|
| XGBoost | Small | +0.094 [+0.035, +0.153] | -0.017 [-0.059, +0.027] | -0.027 [-0.044, -0.009] |
| XGBoost | Reference | +0.017 [-0.048, +0.081] | +0.034 [-0.009, +0.080] | +0.004 [-0.013, +0.023] |
| XGBoost | High | +0.081 [+0.025, +0.140] | +0.000 [-0.042, +0.040] | -0.019 [-0.036, -0.002] |
| SVM-L1 | Small | +0.070 [+0.014, +0.129] | -0.038 [-0.082, +0.006] | -0.038 [-0.055, -0.021] |
| SVM-L1 | Reference | +0.088 [+0.033, +0.144] | -0.063 [-0.102, -0.021] | -0.025 [-0.042, -0.009] |
| SVM-L1 | High | +0.032 [-0.030, +0.093] | -0.030 [-0.072, +0.009] | -0.023 [-0.040, -0.006] |
| Boruta-RF | Small | -0.018 [-0.060, +0.020] | +0.013 [-0.019, +0.047] | +0.002 [-0.009, +0.013] |
| Boruta-RF | Reference | +0.020 [-0.016, +0.055] | -0.011 [-0.042, +0.021] | +0.002 [-0.008, +0.011] |
| Boruta-RF | High | -0.005 [-0.050, +0.040] | +0.009 [-0.023, +0.044] | -0.006 [-0.021, +0.008] |

*Note.* Low, Moderate, and High were encoded 0, 1, and 2. Severe errors are direct Low-to-High or High-to-Low classifications. Intervals use 2,000 common resamples stratified by held-out center and true class (seed 42). Negative MAE and severe-error differences favor the locked signature.

**Supplementary Table S12.** AMP-AD cohort characteristics and ordered bScore mapping. Sections a and b report center-level composition and the source-to-analysis bScore mapping, respectively.

**a. Center-level composition and demographics**

| Center | n | Low / Moderate / High | Age at death, median [IQR] | Female, n (%) | APOE4 positive / known | PMI, median [IQR] |
|---|---|---|---|---|---|---|
| Overall | 527 | 111 / 156 / 260 | 81.7 [73.0, 88.0] | 307 (58.3%) | 225/516 (43.6%) | 8.0 [5.1, 14.5]; n=428 |
| Emory | 80 | 19 / 5 / 56 | 75.0 [61.0, 82.0] | 47 (58.8%) | 47/78 (60.3%) | 7.0 [4.5, 15.1]; n=80 |
| Mayo | 217 | 51 / 53 / 113 | 79.0 [71.0, 85.0] | 114 (52.5%) | 91/217 (41.9%) | 11.0 [5.0, 20.8]; n=118 |
| Mount Sinai | 52 | 9 / 15 / 28 | 82.0 [76.8, 90.0] | 28 (53.8%) | 20/52 (38.5%) | 6.9 [4.1, 10.0]; n=52 |
| Rush | 178 | 32 / 83 / 63 | 85.2 [78.7, 90.0] | 118 (66.3%) | 67/169 (39.6%) | 7.6 [5.8, 12.5]; n=178 |

**b. Source-to-analysis bScore mapping**

| Source bScore label | Analysis class | Ordinal code |
|---|---|---|
| None (Stage 0) | Low | 0 |
| Low (Stage I-II) | Low | 0 |
| Moderate (Stage III-IV) | Moderate | 1 |
| High (Stage V-VI) | High | 2 |

*Note.* The local raw-to-analysis bundle contained 529 common frontal-proteomic/metabolomic profiles; two participants with unavailable or unrecognized bScore were excluded. Age values recorded as 90+ were represented by the lower-bound value 90 for median and IQR summaries. APOE4 percentages use participants with known yes/no status; PMI availability is reported in the final column.

**Supplementary Table S13.** CGGA archived-candidate sensitivity comparing the original top-three locking pool with alternative retrospective rules.

| Method | Condition | Rule | Run | n | Dev-CV AUROC | CV regret | Mean Jaccard | Matches recorded |
|---|---|---|---|---|---|---|---|---|
| SVM-L1 | No guidance | Highest dev-CV | 1 | 16 | 0.880 | 0.000 | - | No |
| SVM-L1 | No guidance | Top-three medoid | 4 | 23 | 0.876 | 0.004 | 0.588 | Yes |
| SVM-L1 | No guidance | All-five medoid | 4 | 23 | 0.876 | 0.004 | 0.522 | Yes |
| SVM-L1 | Guided | Highest dev-CV | 1 | 15 | 0.873 | 0.000 | - | No |
| SVM-L1 | Guided | Top-three medoid | 4 | 16 | 0.870 | 0.003 | 0.476 | Yes |
| SVM-L1 | Guided | All-five medoid | 3 | 15 | 0.859 | 0.015 | 0.465 | No |
| XGBoost | No guidance | Highest dev-CV | 3 | 35 | 0.868 | 0.000 | - | Yes |
| XGBoost | No guidance | Top-three medoid | 3 | 35 | 0.868 | 0.000 | 0.372 | Yes |
| XGBoost | No guidance | All-five medoid | 5 | 39 | 0.843 | 0.025 | 0.412 | No |
| XGBoost | Guided | Highest dev-CV | 2 | 20 | 0.849 | 0.000 | - | No |

*Supplementary Table S13 (continued)*

| Method | Condition | Rule | Run | n | Dev-CV AUROC | CV regret | Mean Jaccard | Matches recorded |
|---|---|---|---|---|---|---|---|---|
| XGBoost | Guided | Top-three medoid | 1 | 19 | 0.846 | 0.003 | 0.295 | Yes |
| XGBoost | Guided | All-five medoid | 1 | 19 | 0.846 | 0.003 | 0.239 | Yes |
| Boruta-RF | No guidance | Highest dev-CV | 4 | 10 | 0.817 | 0.000 | - | No |
| Boruta-RF | No guidance | Top-three medoid | 2 | 14 | 0.803 | 0.015 | 0.535 | Yes |
| Boruta-RF | No guidance | All-five medoid | 2 | 14 | 0.803 | 0.015 | 0.541 | Yes |
| Boruta-RF | Guided | Highest dev-CV | 5 | 9 | 0.815 | 0.000 | - | Yes |
| Boruta-RF | Guided | Top-three medoid | 5 | 9 | 0.815 | 0.000 | 0.442 | Yes |
| Boruta-RF | Guided | All-five medoid | 2 | 9 | 0.799 | 0.016 | 0.452 | No |

*Note.* All displayed rules used only the five saved development-run candidates. The highest-development-CV rule ignores overlap; the unrestricted full-bank medoid expands the pool from the three candidates used by the legacy top-three medoid to all five. For every row, CV regret is the highest saved development-CV macro-AUROC minus the score of the candidate selected by that displayed rule. Detailed condition-level records are supplied as a machine-readable companion.

**Supplementary Table S14.** CGGA condition-level feature recurrence across five saved GA runs. Feature-level records are machine-readable companions.

| Method | Condition | Candidate universe | Mean n/run | 5/5 | >=4/5 | >=3/5 | Any run |
|---|---|---|---|---|---|---|---|
| SVM-L1 | No guidance | 35 | 19.0 | 5 | 13 | 18 | 33 |
| SVM-L1 | Guided | 35 | 15.2 | 4 | 7 | 16 | 27 |
| XGBoost | No guidance | 70 | 35.6 | 3 | 15 | 40 | 65 |
| XGBoost | Guided | 70 | 20.0 | 1 | 4 | 13 | 53 |
| Boruta-RF | No guidance | 26 | 11.6 | 3 | 7 | 12 | 19 |
| Boruta-RF | Guided | 26 | 9.0 | 3 | 5 | 6 | 20 |

**Supplementary Table S15.** CGGA component-ablation summary with held-out intervals under the fixed random forest. In this table, "Locked medoid" denotes the archived legacy top-three medoid. The archived evaluation protocol is retained; Supplementary Table S17 provides the coherent same-estimator comparison.

| Method | Variant | Features, n | AUROC [95% CI] | AUPRC [95% CI] | Balanced accuracy [95% CI] |
|---|---|---|---|---|---|
| SVM-L1 | Direct | 35 | 0.635 [0.515, 0.741] | 0.639 [0.540, 0.762] | 0.629 [0.531, 0.726] |
| SVM-L1 | RFECV-only | 15 | 0.633 [0.509, 0.741] | 0.615 [0.526, 0.751] | 0.596 [0.498, 0.694] |
| SVM-L1 | Single GA run | 15 | 0.664 [0.546, 0.769] | 0.663 [0.560, 0.788] | 0.629 [0.530, 0.726] |
| SVM-L1 | Best-development-CV run | 15 | 0.664 [0.546, 0.769] | 0.663 [0.560, 0.788] | 0.629 [0.530, 0.726] |
| SVM-L1 | Locked medoid | 16 | 0.652 [0.535, 0.761] | 0.640 [0.548, 0.775] | 0.608 [0.509, 0.705] |
| SVM-L1 | No target-size guidance | 23 | 0.623 [0.497, 0.733] | 0.598 [0.512, 0.737] | 0.618 [0.520, 0.715] |
| XGBoost | Direct | 70 | 0.684 [0.565, 0.785] | 0.675 [0.573, 0.797] | 0.652 [0.552, 0.749] |
| XGBoost | RFECV-only | 20 | 0.675 [0.553, 0.773] | 0.685 [0.582, 0.789] | 0.620 [0.512, 0.716] |
| XGBoost | Single GA run | 19 | 0.702 [0.587, 0.803] | 0.689 [0.592, 0.820] | 0.672 [0.574, 0.761] |
| XGBoost | Best-development-CV run | 20 | 0.716 [0.599, 0.811] | 0.732 [0.630, 0.854] | 0.684 [0.586, 0.772] |
| XGBoost | Locked medoid | 19 | 0.702 [0.587, 0.803] | 0.689 [0.592, 0.820] | 0.672 [0.574, 0.761] |
| XGBoost | No target-size guidance | 35 | 0.645 [0.525, 0.753] | 0.632 [0.539, 0.766] | 0.673 [0.575, 0.762] |
| Boruta-RF | Direct | 26 | 0.658 [0.538, 0.765] | 0.646 [0.554, 0.780] | 0.629 [0.531, 0.727] |
| Boruta-RF | RFECV-only | 9 | 0.682 [0.565, 0.785] | 0.653 [0.558, 0.781] | 0.651 [0.552, 0.748] |
| Boruta-RF | Single GA run | 9 | 0.685 [0.574, 0.790] | 0.732 [0.635, 0.825] | 0.673 [0.576, 0.762] |
| Boruta-RF | Best-development-CV run | 9 | 0.695 [0.582, 0.793] | 0.689 [0.593, 0.810] | 0.652 [0.554, 0.739] |
| Boruta-RF | Locked medoid | 9 | 0.695 [0.582, 0.793] | 0.689 [0.593, 0.810] | 0.652 [0.554, 0.739] |
| Boruta-RF | No target-size guidance | 14 | 0.666 [0.550, 0.773] | 0.661 [0.565, 0.779] | 0.663 [0.565, 0.751] |

**Supplementary Table S16.** Exploratory CGGA same-size random-subset and Elastic Net comparator results. Sections a–c report matched-size point estimates, random-subset tail summaries, and paired locked-medoid-minus-Elastic Net differences, respectively. In this table, "Locked medoid" denotes the archived legacy top-three medoid. The archived evaluation protocol is retained; Supplementary Table S17 provides the coherent same-estimator comparison.

**a. Matched-size point estimates**

| Method | Comparator | n | AUROC | AUPRC | Balanced accuracy |
|---|---|---|---|---|---|
| SVM-L1 | Locked medoid | 16 | 0.656 | 0.642 | 0.597 |
| SVM-L1 | Elastic Net | 16 | 0.635 | 0.599 | 0.661 |
| XGBoost | Locked medoid | 19 | 0.689 | 0.669 | 0.695 |
| XGBoost | Elastic Net | 19 | 0.640 | 0.623 | 0.617 |
| Boruta-RF | Locked medoid | 9 | 0.694 | 0.700 | 0.652 |
| Boruta-RF | Elastic Net | 9 | 0.612 | 0.608 | 0.607 |

**b. Random-subset tail summaries**

| Method | Metric | Comparator | Observed | Null mean | Percentile | p (one-sided) |
|---|---|---|---|---|---|---|
| SVM-L1 | AUROC | Locked medoid | 0.656 | 0.615 | 84.3 | 0.162 |
| SVM-L1 | AUROC | Elastic Net | 0.635 | 0.615 | 67.5 | 0.326 |
| SVM-L1 | AUPRC | Locked medoid | 0.642 | 0.619 | 71.5 | 0.286 |
| SVM-L1 | AUPRC | Elastic Net | 0.599 | 0.619 | 32.8 | 0.672 |
| SVM-L1 | Balanced accuracy | Locked medoid | 0.597 | 0.585 | 67.0 | 0.348 |
| SVM-L1 | Balanced accuracy | Elastic Net | 0.661 | 0.585 | 98.2 | 0.019 |
| XGBoost | AUROC | Locked medoid | 0.689 | 0.620 | 94.3 | 0.058 |
| XGBoost | AUROC | Elastic Net | 0.640 | 0.620 | 64.7 | 0.357 |
| XGBoost | AUPRC | Locked medoid | 0.669 | 0.624 | 83.8 | 0.163 |
| XGBoost | AUPRC | Elastic Net | 0.623 | 0.624 | 48.9 | 0.511 |
| XGBoost | Balanced accuracy | Locked medoid | 0.695 | 0.592 | 99.3 | 0.008 |
| XGBoost | Balanced accuracy | Elastic Net | 0.617 | 0.592 | 68.3 | 0.319 |
| Boruta-RF | AUROC | Locked medoid | 0.694 | 0.637 | 95.0 | 0.052 |
| Boruta-RF | AUROC | Elastic Net | 0.612 | 0.637 | 23.4 | 0.767 |
| Boruta-RF | AUPRC | Locked medoid | 0.700 | 0.631 | 93.5 | 0.066 |
| Boruta-RF | AUPRC | Elastic Net | 0.608 | 0.631 | 31.5 | 0.685 |
| Boruta-RF | Balanced accuracy | Locked medoid | 0.652 | 0.619 | 87.0 | 0.135 |
| Boruta-RF | Balanced accuracy | Elastic Net | 0.607 | 0.619 | 33.0 | 0.684 |

**c. Paired locked-medoid-minus-Elastic Net differences**

| Method | Metric | Difference | 95% bootstrap CI | Resamples |
|---|---|---|---|---|
| SVM-L1 | AUROC | +0.021 | [-0.083, +0.131] | 2,000 |
| SVM-L1 | AUPRC | +0.043 | [-0.060, +0.149] | 2,000 |
| SVM-L1 | Balanced accuracy | -0.063 | [-0.173, +0.047] | 2,000 |
| XGBoost | AUROC | +0.049 | [-0.059, +0.170] | 2,000 |
| XGBoost | AUPRC | +0.045 | [-0.062, +0.171] | 2,000 |
| XGBoost | Balanced accuracy | +0.078 | [-0.031, +0.196] | 2,000 |
| Boruta-RF | AUROC | +0.082 | [-0.030, +0.200] | 2,000 |
| Boruta-RF | AUPRC | +0.093 | [-0.035, +0.200] | 2,000 |
| Boruta-RF | Balanced accuracy | +0.046 | [-0.075, +0.165] | 2,000 |

**Supplementary Table S17.** CGGA coherent same-split, same-estimator benchmark

| Branch | Development-derived signature | Features | Held-out AUROC (95% CI) | AUPRC | Balanced accuracy |
|---|---|---|---|---|---|
| SVM-L1 | RFECV-only | 15 | 0.633 [0.509, 0.741] | 0.615 | 0.596 |
| SVM-L1 | Elastic Net | 16 | 0.644 [0.523, 0.754] | 0.611 | 0.649 |
| SVM-L1 | Highest CV | 15 | 0.664 [0.546, 0.769] | 0.663 | 0.629 |
| SVM-L1 | Legacy top-three medoid | 16 | 0.652 [0.535, 0.761] | 0.640 | 0.608 |
| SVM-L1 | Full-bank medoid | 15 | 0.672 [0.551, 0.776] | 0.672 | 0.652 |
| SVM-L1 | Regret-constrained medoid | 16 | 0.652 [0.535, 0.761] | 0.640 | 0.608 |
| XGBoost | RFECV-only | 20 | 0.680 [0.561, 0.779] | 0.677 | 0.663 |
| XGBoost | Elastic Net | 19 | 0.636 [0.513, 0.746] | 0.622 | 0.639 |
| XGBoost | Highest CV | 20 | 0.699 [0.581, 0.799] | 0.719 | 0.684 |
| XGBoost | Legacy top-three medoid | 19 | 0.700 [0.582, 0.805] | 0.685 | 0.695 |
| XGBoost | Full-bank medoid | 19 | 0.700 [0.582, 0.805] | 0.685 | 0.695 |
| XGBoost | Regret-constrained medoid | 21 | 0.645 [0.527, 0.750] | 0.686 | 0.554 |
| Boruta-RF | RFECV-only | 9 | 0.686 [0.571, 0.788] | 0.669 | 0.629 |
| Boruta-RF | Elastic Net | 9 | 0.606 [0.487, 0.716] | 0.598 | 0.596 |
| Boruta-RF | Highest CV | 9 | 0.694 [0.582, 0.792] | 0.700 | 0.652 |
| Boruta-RF | Legacy top-three medoid | 9 | 0.694 [0.582, 0.792] | 0.700 | 0.652 |
| Boruta-RF | Full-bank medoid | 9 | 0.704 [0.594, 0.804] | 0.674 | 0.695 |
| Boruta-RF | Regret-constrained medoid | 9 | 0.694 [0.582, 0.792] | 0.700 | 0.652 |

All signatures use the same fixed 214/92 split and the same 500-tree random forest (maximum depth 12, minimum leaf size 2, balanced class weighting, random state 42). Feature selection was completed within development data before held-out evaluation. Intervals use 2,000 class-stratified resamples (seed 42). Elastic Net signatures were size matched within branch. No comparison establishes predictive superiority; paired contrasts are shown in Supplementary Fig. S18c and supplied machine-readable outputs.

**Supplementary Table S18.** CGGA saved nested-bank re-locking using the regret-constrained medoid

| Fold | Legacy run | Selected run | Features | Eligible | Regret | Legacy AUROC | Selected AUROC |
|---|---|---|---|---|---|---|---|
| 1 | 0 | 3 | 15 | 2 | 0.0000 | 0.762 | 0.703 |
| 2 | 2 | 4 | 16 | 2 | 0.0000 | 0.615 | 0.621 |
| 3 | 1 | 2 | 15 | 1 | 0.0000 | 0.784 | 0.779 |
| 4 | 4 | 0 | 15 | 1 | 0.0000 | 0.745 | 0.675 |
| 5 | 0 | 2 | 15 | 2 | 0.0000 | 0.600 | 0.573 |

This post hoc internal sensitivity reuses five archived candidate banks from a reduced search budget (20 individuals, 12 generations, five runs) and does not rerun the GA or access the original 92-participant held-out cohort. The regret-constrained medoid changed the selected run in all five outer folds. OOF AUROC was 0.692 for the legacy top-three medoid and 0.667 for the regret-constrained medoid; this adverse result is retained because the regret guarantee concerns the configured development score gap.

**Supplementary Table S19.** VGH brain-tumour radiomics internal demonstration (UBC REB H20-02354) under the corrected-objective WrapEvoFS configuration.

**a. Development-only feature counts and strict locking audit**

| Feature space | Branch | Direct | RFECV $k^*$ | Five-run range | Locked | Compression | $\lvert E_{0.01}\rvert$ | Regret |
|---|---|---|---|---|---|---|---|---|
| Full 1,781 | SVM-L1 | 214 | 8 | 50–62 | 50 | 76.6% | 1 | 0.0000 |
| Full 1,781 | XGBoost | 100 | 16 | 17–24 | 24 | 76.0% | 1 | 0.0000 |
| Full 1,781 | Boruta-RF | 10 | 10 | 9–10 | 9 | 10.0% | 2 | 0.0000 |
| Filtered 1,346 | SVM-L1 | 201 | 3 | 46–52 | 50 | 75.1% | 1 | 0.0000 |
| Filtered 1,346 | XGBoost | 100 | 6 | 17–22 | 19 | 81.0% | 1 | 0.0000 |
| Filtered 1,346 | Boruta-RF | 6 | 6 | 4–6 | 5 | 16.7% | 1 | 0.0000 |

**b. Five-seed agreement and one-time held-out evaluation**

| Feature space | Branch | Mean Jaccard | Nogueira Φ | Cross-space locked Jaccard | Locked AUROC (95% CI) | Locked–Direct Δ (95% CI) | Locked–RFECV Δ (95% CI) |
|---|---|---|---|---|---|---|---|
| Full 1,781 | SVM-L1 | 0.153 | -0.005 | 0.053 | 0.567 (0.420, 0.709) | 0.069 (-0.021, 0.166) | -0.066 (-0.190, 0.053) |
| Full 1,781 | XGBoost | 0.153 | 0.071 | 0.049 | 0.565 (0.414, 0.715) | 0.042 (-0.050, 0.137) | -0.060 (-0.139, 0.013) |
| Full 1,781 | Boruta-RF | 0.920 | -0.042 | 0.273 | 0.565 (0.407, 0.708) | -0.001 (-0.053, 0.056) | -0.001 (-0.053, 0.056) |
| Filtered 1,346 | SVM-L1 | 0.144 | 0.002 | 0.053 | 0.505 (0.345, 0.652) | 0.006 (-0.064, 0.078) | -0.012 (-0.167, 0.141) |
| Filtered 1,346 | XGBoost | 0.133 | 0.047 | 0.049 | 0.431 (0.279, 0.590) | -0.007 (-0.126, 0.112) | -0.068 (-0.226, 0.087) |
| Filtered 1,346 | Boruta-RF | 0.767 | -0.010 | 0.273 | 0.565 (0.413, 0.715) | 0.000 (-0.062, 0.060) | 0.000 (-0.062, 0.060) |

*Note.* The same 197 participants and frozen 70:30 split were used for both feature spaces. The 1,346-feature space is a label-independent feature-definition sensitivity analysis, not an external cohort. Regret is the selected empirical development-CV score gap under absolute $\delta = 0.01$. Mean Jaccard and Nogueira Φ summarize five stochastic seeds on one development sample and do not measure participant-resampling or biomarker stability. Held-out intervals use 2,000 class-stratified bootstrap resamples. Every paired interval includes zero. Exact duplicate masks occurred only within the two Boruta-RF candidate banks; the package-retained multiplicity policy was used, and no duplicate mask was selected through a hash-dependent tie. Automatic tumor-core regions were not manually adjudicated.

**Supplementary Table S20.** Full-feature Bayesian sampling, global-parameter, and coefficient-level diagnostic summary. Sections a–c report chain sampling, global-parameter diagnostics, and coefficient-level diagnostics, respectively.

**a. Chain-sampling diagnostics**

| Chain | Div. | Mean accept. | Median steps | Max steps | Time (min) |
|---|---|---|---|---|---|
| 1 | 0 | 0.927 | 63 | 63 | 16.10 |
| 2 | 0 | 0.958 | 63 | 63 | 16.29 |
| 3 | 0 | 0.966 | 63 | 63 | 13.33 |
| 4 | 0 | 0.953 | 63 | 63 | 10.59 |

**b. Global-parameter diagnostics**

| Contrast | Parameter | Mean | 95% CrI | R-hat | ESS |
|---|---|---|---|---|---|
| MCI vs CN | Intercept | 0.6111 | [0.3591, 0.8753] | 1.0000 | 4223.6 |
| MCI vs CN | Global shrinkage scale | 0.4007 | [0.0114, 1.2820] | 1.0086 | 438.7 |
| MCI vs CN | Effective coefficient scale | 0.0047 | [0.0001, 0.0149] | 1.0086 | 438.7 |
| Dementia vs CN | Intercept | 0.6049 | [0.2074, 1.0030] | 1.0010 | 2808.0 |
| Dementia vs CN | Global shrinkage scale | 3.8536 | [2.8487, 5.0184] | 1.0000 | 1744.0 |
| Dementia vs CN | Effective coefficient scale | 0.0448 | [0.0331, 0.0583] | 1.0000 | 1744.0 |

**c. Coefficient-level diagnostics**

| Contrast | Coefficients | Max R-hat | R>1.01 | R>1.05 | Min ESS |
|---|---|---|---|---|---|
| MCI vs CN | 7,411 | 1.0020 | 0 | 0 | 3413.9 |
| Dementia vs CN | 7,411 | 1.0018 | 0 | 0 | 4465.5 |
| Dementia vs MCI | 7,411 | 1.0020 | 0 | 0 | 4447.7 |

**Supplementary Table S21.** Candidate-restricted Bayesian, training-only STABL, and BLiP integration summary. Sections a–e report cross-method overlap, posterior summaries, model diagnostics, held-out summaries, and BLiP settings, respectively.

**a. Cross-method overlap**

| Feature | Omic | SVM-L1 | XGB | Boruta-RF | STABL | Primary | q=.10 | OR $\geq$ 1.20 | practical-effect probability |
|---|---|---|---|---|---|---|---|---|---|
| YWHAG | Proteomics | Yes | Yes | No | Yes | Yes | Yes | Yes | 0.98175 |
| ACHE.1 | Proteomics | Yes | No | Yes | Yes | Yes | Yes | Yes | 0.99725 |
| PPP3CA\|PPP3R1 | Proteomics | No | No | Yes | Yes | Yes | Yes | Yes | 0.99975 |
| NEFH | Proteomics | Yes | No | Yes | Yes | Yes | Yes | Yes | 0.89275 |
| S-methylcysteine sulfoxide | Metabolomics | Yes | No | No | Yes | Yes | Yes | No | 0.83625 |

**b. Restricted posterior summaries**

| Feature | practical-effect probability | MCI vs CN $\beta$ [95% CrI] | Dementia vs CN $\beta$ [95% CrI] | Dementia vs MCI $\beta$ [95% CrI] |
|---|---|---|---|---|
| YWHAG | 0.98175 | +0.10 [-0.17, +0.51] | +0.77 [+0.03, +1.62] | +0.66 [-0.12, +1.53] |
| ACHE.1 | 0.99725 | -0.12 [-0.47, +0.11] | -0.57 [-0.97, -0.20] | -0.45 [-0.80, -0.12] |
| PPP3CA\|PPP3R1 | 0.99975 | -0.06 [-0.43, +0.21] | +1.18 [+0.44, +1.94] | +1.24 [+0.52, +1.97] |
| NEFH | 0.89275 | +0.05 [-0.17, +0.32] | +0.26 [-0.07, +0.69] | +0.20 [-0.17, +0.61] |
| S-methylcysteine sulfoxide | 0.83625 | -0.10 [-0.33, +0.08] | +0.09 [-0.15, +0.36] | +0.19 [-0.07, +0.46] |

**c. Restricted-model diagnostics**

| Type | Item | Max R-hat | Min ESS | Divergences | Mean acceptance | Median steps |
|---|---|---|---|---|---|---|
| Chain | Chain 1 | — | — | 0 | 0.949 | 255 |
| Chain | Chain 2 | — | — | 0 | 0.937 | 255 |
| Chain | Chain 3 | — | — | 0 | 0.942 | 255 |
| Chain | Chain 4 | — | — | 0 | 0.946 | 255 |
| Coefficient | MCI vs CN | 1.00117 | 3035.55 | — | — | — |
| Coefficient | Dementia vs CN | 1.00124 | 3310.28 | — | — | — |
| Coefficient | Dementia vs MCI | 1.00175 | 3376.57 | — | — | — |
| Global | alpha: MCI vs CN | 0.99988 | 3775.91 | — | — | — |
| Global | alpha: Dementia vs CN | 1.00040 | 3929.67 | — | — | — |
| Global | tau_global: MCI vs CN | 1.00197 | 1484.87 | — | — | — |
| Global | tau_global: Dementia vs CN | 1.00086 | 1852.62 | — | — | — |

**d. Held-out summaries**

| Analysis | Features, n | Balanced accuracy (95% CI) | Macro F1 (95% CI) | Macro OvR AUROC (95% CI) |
|---|---|---|---|---|
| Full-feature Bayesian reference (Supplementary Fig. S24) | 7411 | 0.485 [CI not reported] | 0.433 [CI not reported] | 0.780 [CI not reported] |
| Restricted Laplace Bayesian model | 81 | 0.548 [0.468, 0.628] | 0.551 [0.461, 0.638] | 0.803 [0.744, 0.856] |
| Candidate-union random forest | 81 | 0.572 [0.495, 0.649] | 0.574 [0.478, 0.659] | 0.788 [0.734, 0.840] |
| STABL random forest | 8 | 0.468 [0.395, 0.545] | 0.465 [0.383, 0.545] | 0.706 [0.639, 0.768] |

**e. BLiP settings**

| Setting | OR threshold | q | Selected, n | Overlap retained, n/5 |
|---|---|---|---|---|
| Primary | 1.10 | 0.20 | 81 | 5/5 |
| Stricter FDR | 1.10 | 0.10 | 47 | 5/5 |
| Smaller effect | 1.05 | 0.20 | 81 | 5/5 |
| Larger effect | 1.20 | 0.20 | 32 | 4/5 |

**Supplementary Table S22.** Held-out predictive performance of the full-feature Bayesian multinomial logistic reference. Sections a and b report overall and class-specific performance, respectively.

**a. Overall performance**

| Metric | Value |
|---|---|
| Accuracy | 0.594 |
| Balanced accuracy | 0.485 |
| Macro F1 | 0.433 |
| Weighted F1 | 0.533 |
| Multiclass log loss | 0.848 |
| Macro one-vs-rest ROC AUC | 0.780 |

**b. Class-specific performance**

| Metric | CN | MCI | Dementia |
|---|---|---|---|
| ROC AUC | 0.819 | 0.668 | 0.852 |
| Average precision | 0.480 | 0.497 | 0.839 |
| Precision | 0.000 | 0.476 | 0.700 |
| Sensitivity | 0.000 | 0.638 | 0.817 |
| Specificity | 1.000 | 0.593 | 0.740 |
| F1 score | 0.000 | 0.545 | 0.754 |

**Supplementary Table S23.** Controlled synthetic candidate-bank locking simulation under the primary $R = 5$ and $\delta = 0.01$ design.

**a. Equal-scenario primary operating characteristics**

| Rule | Oracle regret | Group recovery | Exact recovery | Full-bank Jaccard | Score optimism | Failure (%) |
|---|---|---|---|---|---|---|
| Highest score | 0.0022 | 0.851 | 0.749 | 0.424 | 0.0143 | 0.145 |
| Legacy top-three medoid | 0.0025 | 0.847 | 0.755 | 0.460 | 0.0073 | 0.032 |
| Full-bank medoid | 0.0023 | 0.851 | 0.771 | 0.475 | 0.0006 | 0.014 |
| Regret-constrained medoid | 0.0026 | 0.846 | 0.751 | 0.446 | 0.0133 | 0.128 |

**b. Primary empirical-regret audit**

| Rule | Mean regret | Scenario means > 0.01 | Maximum scenario q95 | Individual guarantee audit |
|---|---|---|---|---|
| Highest score | 0.0000 | 0 / 162 | 0.0000 | 0 by definition |
| Legacy top-three medoid | 0.0073 | 46 / 162 | 0.0783 | Not constrained |
| Full-bank medoid | 0.0138 | 67 / 162 | 0.1226 | Not constrained |
| Regret-constrained medoid | 0.0013 | 0 / 162 | 0.0091 | 0 / 4,995,000 |

**c. Regret-constrained-medoid primary decision paths**

| Decision stage | Probability (%) |
|---|---|
| Singleton eligible pool | 28.297 |
| Unique Jaccard medoid | 45.547 |
| Higher locking score | 25.146 |
| Smaller feature count | 0.00025 |
| Stable mask hash | 0.360 |
| Exact-duplicate provenance | 0.649 |
| Hash stage reached (inclusive) | 1.009 |

Primary entries are equal-scenario means across 162 prespecified scenarios with 10,000 banks each (1,620,000 banks); the complete sensitivity workload comprised 4,995,000 banks across 513 scenarios. Hidden oracle utility $T$ is not AUROC. Lower values are preferable for oracle regret and score optimism; higher values are preferable for recovery and Jaccard. Failure denotes oracle regret $> 0.02$; q95 denotes the within-scenario 95th percentile. The regret-constrained medoid had no configured empirical-regret violation in the complete workload. Highest-score selection has zero empirical regret by definition, whereas the two medoid comparators have no metric-scale regret constraint. Section c probabilities are mutually exclusive except for the inclusive hash-stage row. Exact duplicate masks retained their multiplicity as voting candidates. No GA, classifier, or participant-level dataset was simulated or fitted.

**Supplementary Table S24.** Executable WrapEvoFS v0.2.0 method specification and corrected-objective analysis settings.

| **Stage** | **Exact implementation and operating specification** |
|---|---|
| Scope and boundary | Values below describe the v0.2.0 implementation and corrected-objective 50-by-50 analyses unless an archived experiment supplied a dataset-specific metric or RFECV cap. Archived original-analysis overrides are reported in Supplementary Table S27. Except in the fully nested TCGA experiment, preprocessing and Direct screening are fitted on the full development partition before RFECV, GA scoring, and locking rescoring and are not nested inside the later scoring folds. In TCGA, all study-fitted steps from the supplied matrix were refitted within every outer-training partition; the provider's label-independent 5,000-feature SCNV variance filter preceded outer CV. |
| SVM-L1 Direct selector | Five shuffled stratified folds (random state 42); internal `StandardScaler`; `LinearSVC` with L1 penalty, `dual=False`, tolerance $10^{-4}$, maximum 100,000 iterations, and no class weighting. The grid is $C \in \{0.001, 0.01, 0.1, 1, 10, 30, 100\}$. Auto scoring resolves to AUROC for binary outcomes and balanced accuracy for multiclass outcomes. Features with absolute coefficient greater than $10^{-6}$ are retained; multiclass coefficients are aggregated by the maximum absolute value across classes. |
| XGBoost Direct selector | Five shuffled stratified folds (random state 42); histogram tree method; learning rate 0.3; gamma 0; maximum depth 6; $L_2$ penalty 1; $L_1$ penalty 0. The search grid is subsample $\in \{0.6, 0.7, 0.8\}$, column subsample $\in \{0.7, 0.8, 1.0\}$, and trees $\in \{80, 100, 120\}$. Auto scoring resolves to AUROC for binary outcomes and macro one-vs-rest AUROC for multiclass outcomes. Positive-importance features are ordered stably and truncated to the first 100. |
| Boruta-RF Direct selector | Random forest with 1,000 trees, maximum depth 5, balanced class weights, and random state 42; `BorutaPy` uses automatic internal tree count, percentile 95, $\alpha = 0.05$, two-step correction, maximum 150 iterations, and random state 42. Confirmed and tentative features are both retained. |
| RFECV target | Random forest with 150 trees, maximum depth 12, minimum leaf size 2, bootstrap sampling, and random state 42; five shuffled stratified folds; step size 1; minimum one feature. Auto scoring is AUROC for binary outcomes and macro one-vs-rest AUROC for multiclass outcomes. The v0.2.0 compact caps are 20 features for SVM-L1 and XGBoost and 25 for Boruta-RF. The fully nested TCGA experiment used branch-specific caps of 15, 20, and 10, respectively; archived original-analysis caps are listed in Supplementary Table S27. The selected target is the smallest feature count attaining the largest mean score among eligible counts. |

*Supplementary Table S24 (continued)*

| Stage | Exact implementation and operating specification |
|---|---|
| GA evaluator and budget | Five retained runs, population $N = 50$, and $G = 50$ generations. Each chromosome is evaluated by five-fold stratified development CV using a random forest with 150 trees, maximum depth 12, minimum leaf size 2, and bootstrap sampling. Corrected-objective GPU runs use the cuML backend with 16 histogram bins; CPU fallback uses the scikit-learn estimator with the same listed scientific RF parameters. The search metric is dataset-specific and is distinct from the later locking metric when configured as such. |
| Population initialization | For candidate-universe size $p_m$, every initial chromosome starts with exactly $\lfloor 0.30p_m \rfloor$ zeros and $p_m - \lfloor 0.30p_m \rfloor$ ones; bit positions are shuffled independently. Any empty mask is repaired by activating one uniformly sampled feature. |
| Objective and ranking | Target-deviation penalty $\lambda = 0.015$. Archived compatibility mode uses `legacy_zero_truncated_linear`; corrected-objective analyses use `untruncated_shifted_linear`. Corrected-objective population ranking and run-best updating use, in order, larger raw objective, larger base score, smaller absolute target deviation, smaller feature count, and lexicographically smaller stable mask hash. |
| Elitism and parent sampling | The best $e = 2$ chromosomes are copied unchanged. The remaining $N - e = 48$ parent indices are sampled independently with replacement from normalized shifted weights. If no raw objective is finite, the finite-objective range is at most $10^{-12}$, or the weight total is nonfinite or at most $10^{-12}$, the probabilities are set uniformly and the fallback is recorded. |
| Crossover, mutation, and repair | Sampled parents are paired in list order. One-point crossover is applied with probability 0.8, with its cut sampled uniformly from internal positions $\{1, \ldots, p_m - 2\}$ when $p_m > 2$; otherwise the parents are copied. Each child bit is flipped independently with probability 0.05. An empty child is repaired by activating one uniformly sampled feature; children are appended after elites until the next population reaches size $N$. |
| GA random-seed schedule | The genetic-operator generator for zero-based run $r$ uses seed $\xi_0 + r$. For zero-based generation $g$, current population-list index $i$, and fold $\ell$, the random-forest evaluator uses $\xi_{rgi\ell} = \xi_0 + 10000r + 100g + 1009i + \ell$. Consequently, stochastic RF evaluation is indexed by run, generation, current chromosome position, and fold; the schedule is deterministic from these indices and the exported base seed. |
| Fixed locking rescoring | One shuffled stratified five-fold partition is constructed with random state 42 and reused for all retained candidates in a bank. Each candidate uses the same 150-tree random-forest specification (maximum depth 12, minimum leaf size 2, bootstrap) and a recorded run-specific estimator seed; when no explicit seed map is supplied, the seed is locking random state plus stored run ID. The mean of the five fold scores is $L_r$. |
| Representative locking | Regret-constrained analyses use absolute $\delta = 0.01$. Eligible candidates satisfy the exact condition $L_{\text{best}} - L_r \leq \delta$ without tolerance-based pool expansion. A singleton is selected directly; otherwise candidates are ordered by larger eligible-pool mean Jaccard, larger $L_r$, smaller feature count, and lexicographically smaller canonical-mask identifier. Exact duplicate masks retain voting multiplicity; source-run ID is post-selection provenance only. |
| Audit and held-out boundary | Outputs retain fold and mean scores, objectives, penalties, seeds, masks, hashes, duplicate multiplicity, regrets, eligibility, Jaccard values, tie paths, resolved backend, software/schema version, configuration hash, and input/candidate-universe fingerprints. Held-out data are not inputs to Direct selection, RFECV, GA evaluation, candidate rescoring, eligibility, locking, or audit export. |

**Supplementary Table S25.** Implementation correspondence for the formal regret-constrained locking properties.

| **Property or assumption** | **Package mechanism** | **Verification evidence** |
|---|---|---|
| Nonempty bank and feature sets | Candidate validation rejects empty banks, empty masks, duplicate feature names, and features outside the canonical universe. | Empty-bank, empty-mask, and universe-validation tests. |
| Finite larger-is-better scores | Scores and fold scores must be finite, and regret locking accepts only the explicit `larger_is_better` orientation. | Nonfinite-score and lower-orientation misuse tests. |
| Propositions 1–2 | The best score is a maximum; eligibility uses exact $R_r \leq \delta$; every selection path is restricted to the eligible indices; a postcondition verifies the selected regret. | Singleton, hidden-epsilon boundary, and generated-bank strict-feasibility tests. |
| Propositions 3–4 | $E_\delta$ is constructed directly from the configured threshold, including zero tolerance and the full-bank limit. | Unique-best and tied-best zero-tolerance tests; all-candidate-eligible test. |
| Proposition 5 | A common ordered universe defines uint8 masks; the unchanged canonical-mask encoding helper defines stable identifiers; audit rows and metadata are canonically ordered. | Cross-process hash, feature-order canonicalization, serialization/reload, and repeated-audit tests. |
| Proposition 6 | Scientific ties depend on mask-derived content, not input-row order. Source-run ID labels provenance only after an exact duplicate feature set wins. | Exhaustive record-permutation and generated-bank permutation tests. |
| Duplicate-mask policy | Duplicate masks are not deduplicated and their multiplicity contributes to eligible-pool means; duplicate multiplicity and all source-run provenance are serialized. | Duplicate-mask and identical-mask/identical-score tests. |
| Best-run-SE-scaled sensitivity boundary | AMP-AD development-fold score vectors were not archived, so this threshold was not estimated from means or standard deviations. | Unavailable-input audit; no numerical reconstruction was attempted. |
| Archived candidate-bank recovery | ADNI Drive artifacts retained all five masks and development-CV locking scores per branch, permitting strict retrospective re-locking without rerunning selection or accessing held-out outcomes. | Exact mask-to-feature decoding, legacy-lock reproduction, finite-score and duplicate audits, and all 120 input-record permutations per branch. |
| Proposition 7 and claim boundary | Separate highest-locking-score-candidate, unrestricted-full-bank-medoid, original top-$k$ (legacy-compatible), and regret-constrained-medoid modes retain their distinct feasible sets and objectives. | Definition-level proof and strategy-specific unit tests; predictive or external-validity claims are not machine-testable. |

**Supplementary Table S26.** Dataset preprocessing and application boundaries.

| **Dataset** | **Supplied matrix / preprocessing boundary** | **Development-only fitted operations** | **Held-out application** |
|---|---|---|---|
| ADNI | 7,464 raw predictors; fixed 80:20 split | >20% missingness filter; minimum-positive/2 imputation; log2; zero-variance filter | Development-derived feature rules and imputations applied unchanged |
| CGGA | 24,326 aligned expression predictors; fixed 70:30 split | >20% missingness filter and minimum-positive/2 logic were encoded in the source notebook; no feature/cell changes were required for the supplied matrices | Locked 92-participant partition accessed after model locking |
| AMP-AD | 13,136 supplied predictors; four fixed LOCO folds | >20% missingness filter; median imputation; zero-variance filter within each development fold | Fitted transformations applied unchanged to each held-out center |
| TCGA GBM/LGG | 491 participants; 20,118 supplied RNA-seq predictors and 5,000 provider-filtered SCNV predictors; the SCNV boundary used label-independent variance filtering; two stratified five-fold outer-CV repeats | All study-fitted preprocessing, Direct screening, RFECV, five GA runs, fixed candidate rescoring, and regret-constrained medoid locking fitted within each outer-training partition | Each outer-test fold was scored once after locking and was unavailable to every selection step |
| VGH brain-tumour radiomics (H20-02354) | 197 participants; fixed 70:30 split; 1,781 full-space and 1,346 perturbation-filtered predictors | Numeric validation and zero imputation; Direct screening, RFECV, five GA runs, and strict locking on development only | Six signatures locked before one evaluation on the 60-participant held-out partition |

**Supplementary Table S27.** Archived original RFECV, GA, and top-three locking settings across the three demonstrations.

| **Dataset** | **RFECV score / eligible cap** | **GA score; runs × population × generations** | **Target guidance** | **Locking** |
|---|---|---|---|---|
| ADNI | 5-fold weighted OvR AUROC; cap 30 per branch | 5-fold balanced accuracy; $5 \times 50 \times 50$ | $\lambda = 0.015$ | Development-CV balanced accuracy; legacy top-three medoid |
| CGGA | 5-fold AUROC; caps 15/20/10 | 5-fold accuracy; $5 \times 50 \times 50$ | $\lambda = 0.015$ guided; $\lambda = 0$ without guidance | Development-CV AUROC; legacy top-three medoid |
| AMP-AD | 5-fold weighted OvR AUROC; XGBoost/SVM 10/20/40, Boruta-RF 13/25/50 | 5-fold balanced accuracy; $5 \times 50 \times 50$ | $\lambda = 0.015$ | Mean 5-fold macro OvR AUROC; legacy top-three medoid |

**Supplementary Table S28.** VGH brain-tumour radiomics upstream construction, extraction, and validation provenance (UBC REB H20-02354).

| Stage | Frozen implementation | Validation and analysis boundary |
|---|---|---|
| DICOM preflight | Header-only grouping by patient, study, series, and instance identity; T1ce eligibility required at least 20 slices, complete geometry, high-confidence sequence classification, and prespecified position and spacing tolerances, with preference for a three-dimensional acquisition. | MGMT was not used for patient, pilot, session, or series selection. The source comprised 326 participants; 262 were T1ce eligible. |
| Conversion | `dcm2niix` v1.0.20260724; read-only source access; pseudonymous NIfTI output with header and extension scrubbing; source-checksum comparison before resume or skip. | Three eligible participants were excluded only after repeated non-unique or unreadable-series conversion failure. No conversion exclusion used MGMT or tumor-mask appearance. |
| Automatic regions | RaidionicsRADS v1.3.2 and RaidionicsSeg v1.5.2; frozen brain- and tumor-model manifests; automatic tumor-core candidate region, label 1. | Empty masks, geometry mismatch, nonfinite voxels, out-of-bounds masks, and execution failure were hard failures. Review flags were retained rather than used as morphology-based exclusions. Regions were not manually corrected or expert adjudicated. |
| Radiomics | PyRadiomics v3.0.1, SimpleITK v2.2.1; normalization scale 100; 1-mm isotropic B-spline image resampling; nearest-neighbor mask resampling; bin width 25; geometry tolerance $10^{-5}$; minimum 3 dimensions and 64 voxels; pre-crop true; pad distance 5. | All final predictor values were finite. Available provenance supports fixed settings but does not independently establish complete IBSI compliance for every upstream transform. |
| Feature families | First-order, shape, GLCM, GLRLM, GLSZM, GLDM, and NGTDM across Original, Wavelet, Laplacian-of-Gaussian (1–5 mm), Gradient, Square, SquareRoot, Logarithm, and Exponential images. | The full analysis boundary contained 1,781 predictors. PyRadiomics diagnostics were excluded from predictors. |
| Stability pilot | Deterministic 15-participant pilot; original, 1-mm-eroded, and 1-mm-dilated tumor-core regions; retain ICC(2,1) $\geq 0.85$ and median relative range $\leq 0.25$. | The 1,346-feature subset was frozen without MGMT access and applied unchanged; it is a segmentation-perturbation robustness subset, not an outcome-selected signature. |
| Cohort linkage | 259 participants completed original radiomics; 197 had unique validated MGMT linkage (80 methylated, 117 unmethylated); 62 remained unlinked. | MGMT was joined after all upstream choices were frozen. Unlinked outcomes were not inferred. The supervised experiment used only the 197 linked participants. |
| Automated validation | All 262 eligible participants accounted for; 259 included and 3 explicitly excluded; image–mask geometry, finite-value, privacy, checksum/resume, schema, and patient-uniqueness checks; 26 automated tests passed. | Sixty-five included masks retained review flags and 49 acquisitions had maximum spacing at least 3 mm. Automatic QC is not clinical contour validation; patient-level QC covariates and images are not redistributed. |

### References


[1] Nogueira, S., Sechidis, K. & Brown, G. On the stability of feature selection algorithms. *J. Mach. Learn. Res.* **18**, 1–54 (2018).

[2] Li, X., Morgan, P. S., Ashburner, J., Smith, J. & Rorden, C. The first step for neuroimaging data analysis: DICOM to NIfTI conversion. *J. Neurosci. Methods* **264**, 47–56 (2016).

[3] Bouget, D. *et al.* Raidionics: an open software for pre- and postoperative central nervous system tumor segmentation and standardized reporting. *Sci. Rep.* **13**, 15570 (2023).

[4] van Griethuysen, J. J. M. *et al.* Computational radiomics system to decode the radiographic phenotype. *Cancer Res.* **77**, e104–e107 (2017).
